# Association of variants within the *GST* and other genes with anti-tubercular agents related toxicity: a systematic review and meta-analysis

**DOI:** 10.1101/515817

**Authors:** Marty Richardson, Jamie Kirkham, Kerry Dwan, Derek J Sloan, Geraint Davies, Andrea L Jorgensen

**Author notes:** **Systematic review registration** PROSPERO registration number: CRD42017068448. Corresponding author Marty Richardson, Whelan Building, University of Liverpool, L69 3GB, UK.

## Abstract

**Background:** Individuals receiving treatment with anti-tuberculosis (TB) drugs may experience serious side-effects, such as anti-TB drug-induced hepatotoxicity (ATDH). Genetic variants, such as polymorphisms of the *GST* gene and other genes, may increase the risk of experiencing such toxicity events. This systematic review and meta-analysis provides a comprehensive evaluation of the evidence base for associations between variants of the *GST* gene and other genes and toxicity outcomes related to anti-TB drugs.

**Methods:** We searched for relevant studies in MEDLINE, PubMed, EMBASE, BIOSIS and Web of Science. We pooled effect estimates for each genotype on each outcome, and stratified all analyses by country. We qualitatively assessed the methodological quality of the included studies.

**Results:** We included data from 28 distinct cohorts of patients in the review. The methodological quality of included studies was variable, with several important areas of concern. For *GSTM1*, patients with the homozygous null genotype were significantly more likely to experience hepatotoxicity than patients with heterozygous or homozygous present genotype (odds ratio [OR]=1.44, 95% confidence interval [CI] 1.15, 1.82). Moderate heterogeneity was observed in this analysis (I^2^=51.2%). No significant difference was observed for the *GSTT1* null polymorphism. For the rs3814057 polymorphism of the *PXR* gene, both heterozygous genotype and homozygous mutant-type significantly increased hepatotoxicity risk compared with homozygous wild-type (heterozygous versus homozygous wild-type: OR=1.98, 95% CI 1.06, 3.69; I^2^=0%; homozygous mutant-type versus homozygous wild-type: OR=2.18, 95% CI 1.07, 4.44; I^2^=0%).

**Conclusions:** We found that it is challenging to perform robust synthesis of the evidence base for associations between *GST* and other genetic variants and toxicity related to anti-TB drugs. We identified significant associations between the *GSTM1* null and *PXR* rs3814057 polymorphisms and ATDH. To the best of our knowledge, no meta-analyses on genetic variants other than variants of the *NAT2*, *CYP2E1*, *GSTM1* and *GSTT1* genes have been published. Our results therefore add to the existing understanding of the association between genetic variants and hepatotoxicity.

## Background

Tuberculosis (TB) is the ninth most common cause of death worldwide, and is the leading cause of death from a single infectious agent (1). The World Health Organisation currently recommends that individuals with drug susceptible TB should be treated with a combination of four first-line anti-TB drugs: isoniazid, rifampicin, ethambutol and pyrazinamide (1).

Individuals receiving treatment with anti-TB drugs may experience serious side-effects, such as anti-TB drug-induced hepatotoxicity (ATDH). Reported incidence rates of ATDH among patients treated with standard combination treatment vary from 2% to 28%, depending on the treatment regimen, patient characteristics (e.g. age, race and sex), and definition of ATDH (2). In severe cases, ATDH can be fatal, with mortality rates of 6–12% reported in cases where treatment was not stopped promptly (3). ATDH and other anti-TB drug-related adverse reactions may also contribute to poor adherence to treatment, which in turn may result in treatment failure, relapse and the emergence of drug resistance (2).

It has been hypothesised that polymorphisms within the glutathione s-transferase mu 1 (*GSTM1*) and glutathione S-transferase theta-1 (*GSTT1*) genes may be associated with risk of experiencing ATDH, as these genes encode drug metabolising enzymes. *GST* gene polymorphisms may affect the activity of GST enzymes, altering the metabolic pathway of anti-TB drugs in the liver; consequently, patients may experience hepatic adverse reactions. Toxic metabolites may also cause other toxicity outcomes, such as maculopapular eruption (MPE), although hepatotoxicity is the most widely studied toxicity outcome in pharmacogenetic studies of anti-TB drugs.

Polymorphisms of the *NAT2*, *CYP* and *GST* genes have been widely studied in relation to ATDH. Studies investigating associations between *NAT2* and *CYP* genetic variants and anti-TB drug-related toxicity will be reported separately. In this paper, we focus on genetic variants in *GST* as well as variants in other genes that have been investigated previously for association with anti-TB drug-related toxicity outcomes.

The majority of research into the mechanisms of the genetic contribution to ATDH has focussed on isoniazid. It has been hypothesised that individuals with null *GSTM1* or *GSTT1* genotypes may detoxify toxic metabolites from the metabolism of isoniazid less efficiently than those without null *GSTM1* or *GSTT1* genotypes (4). Rifampicin and pyrazinamide have also been reported to be hepatotoxic (5), although the biological mechanisms for rifampicin- and pyrazinamide-induced hepatotoxicity are not well understood (6). The OATP1B1 *15 haplotype has previously been reported to be an important predictor of rifampicin-induced hepatotoxicity (7); no research into genetic predictors of pyrazinamide-induced hepatotoxicity has been reported (8). Ethambutol has not previously been reported to cause hepatotoxicity (5).

The purpose of this systematic review and meta-analysis is to comprehensively summarise the current evidence base on associations between polymorphisms of the *GST* gene and other genetic variants (apart from those in the *NAT2* and *CYP* genes) and anti-TB drug-related toxicity in patients receiving anti-TB treatment.

Meta-analyses investigating the effect of GST genetic variants and toxicity outcomes have been published previously (4, 9–12). However, there are several limitations to these reviews.

- Cai et al. (2012) (4), Li et al. (10) and Tang et al. (12) all excluded studies if data required for meta-analysis were not reported in the study publications. Ideally, meta-analysis authors should contact authors of primary studies to obtain unreported data to facilitate inclusion in the meta-analysis.
- Cai et al. (2012) (4) excluded three studies that were not randomised controlled trials (RCTs), and Cai et al. (2015) (9), Li et al. (10), Sun et al (11) and Tang et al. (12) all included only case–control studies. Pharmacogenetic studies may use RCT, case–control or cohort designs; therefore, study design is not a valid reason for excluding studies, and important evidence may have been omitted from these reviews.
- Cai et al. (2012) (4), Cai et al. (2015) (9) and Li et al. (10) did not assess study quality, which is a key component of any systematic review.

We planned to overcome these limitations in our review by contacting study authors to obtain data required for meta-analysis, by including relevant studies regardless of their design, and by assessing studies with respect to their methodological quality.

In addition, the scope of our review is wider than the previously conducted meta-analyses, all of which excluded studies that did not report hepatotoxicity. Furthermore, to the best of our knowledge, no previously conducted review has aimed to synthesise data for genetic variants other than *NAT2*, *CYP* and *GST* genetic variants. Our review and meta-analysis updates and adds to the current evidence base on associations between genetic variants and anti-TB drug-related toxicity.

## Methods

The current study forms part of a series of systematic reviews and meta-analyses evaluating the influence of different genetic variants on toxicity to anti-TB agents, the protocol for which has been published (PROSPERO registration number: CRD42017068448) (13). This review has been conducted in accordance with the PRISMA statement (14); a completed copy of the PRISMA checklist is provided in Additional file 1.

The search strategy and study selection methods were used to identify studies that investigated the effect of any genetic variant on anti-TB drug-related toxicity outcomes. However, in this article we focus only on the subset of studies that considered genetic variants other than *NAT2* and *CYP* genetic variants; all results reported in this review were obtained from this subset. Studies investigating associations between *NAT2* and *CYP* variants and anti-TB drug-related toxicity will be reported separately.

### Selection criteria

#### Types of studies

Cohort studies, case–control studies and RCTs were eligible for inclusion in our review.

#### Types of participants

We included studies that recruited TB patients who were either already established on anti-TB treatment or were commencing treatment (at least one of isoniazid, rifampicin, pyrazinamide, or ethambutol), and who had been genotyped in order to investigate the effect of genetic variants on anti-TB drug-related toxicity. If we identified a study where only a subgroup of the study patients were TB patients receiving anti-TB treatment, we would include the study if at least 50% of the total sample size belonged to this subgroup.

#### Types of outcomes

We included studies that measured any anti-TB drug-related toxicity outcomes.

### Search strategy

An information specialist (Eleanor Kotas) designed the search strategy (see Additional file 2), and searched for studies in MEDLINE, PubMed, EMBASE, BIOSIS, and Web of Science (date of search: 3 March 2016). We hand searched reference lists from relevant studies, and contacted experts in the field to identify further eligible studies. We included studies published in English only. We did not restrict by publication date or publication status.

### Study selection

The search results were imported into Covidence (15), a web-based software platform that can be used to facilitate the production of systematic reviews. We removed duplicate records, and one author (MR) screened study abstracts to remove obviously irrelevant studies. At this stage, if there was any uncertainty about the relevance of an abstract, the abstract would be included. A second author (AJ, JK or KD) independently screened a sample of 10% of study abstracts.

We obtained the full text for each potentially relevant study. One reviewer (MR) assessed eligibility of each of these studies based on the selection criteria. A second author (AJ, JK or KD) independently assessed the eligibility of a sample of 10% of studies for inclusion. Any disagreements between the two reviewers at the abstract and full-text screening stages were resolved through discussion, and by consulting a third author if necessary.

### Outcomes

The primary outcome of this review was hepatotoxicity by any definition used by the original investigators. The secondary outcomes were all other toxicity outcomes reported in the included studies.

### Data collection

We designed and piloted a data extraction form. We extracted data on study design, participant characteristics, treatment regimen and outcomes. One author (MR) extracted data, in line with methods described in the Cochrane Handbook (16) and The HuGENet HuGE Review Handbook (17). Dual data extraction was performed for all outcome data (second author: AJ, JK or KD). Any disagreements between the two reviewers were resolved through discussion, and by consulting a third author if necessary. We contacted study authors if outcome data necessary for inclusion in a meta-analysis were not reported.

We examined author lists, locations, dates of recruitment and other study characteristics to identify instances of multiple articles reporting outcomes for the same patient cohort, or for overlapping patient cohorts. We contacted the authors of such articles to determine whether patient cohorts were distinct. If an author confirmed that multiple articles reported outcomes for non-distinct patient cohorts, or if we suspected this based on reported study characteristics, we assigned a group identifier (GI) to these articles. This GI ensured that no patients were included more than once (i.e. were double-counted) in any meta-analysis.

### Quality assessment

One author (MR) applied criteria developed by Jorgensen and Williamson (18) specifically for pharmacogenetic studies, to assess the quality of each included study. A second author (AJ) independently assessed the quality of a sample of 10% of studies. Any discrepancies between the two authors were resolved through discussion. We summarised the number of studies meeting each criterion in the text.

### Data synthesis

For all analyses, the measure of treatment effect used was the odds ratio (OR).

#### Primary analyses

The primary analyses compared the risk of ATDH in the following groups:

- patients with homozygous null genotype versus those with heterozygous or homozygous present genotype at *GSTM1*
- patients with homozygous null genotype versus those with heterozygous or homozygous present genotype at *GSTT1*.

Data were pooled from studies that reported data for each genotype group separately and studies that combined homozygous present and heterozygous genotype groups. No studies combined homozygous null and heterozygous genotype groups.

Sensitivity analyses were conducted to investigate the robustness of the primary analyses. We conducted pairwise comparisons of heterozygous versus homozygous present genotype, and homozygous null versus homozygous present genotype for both *GSTM1* and *GSTT1*, using data from studies that reported on each genotype group separately.

We produced funnel plots for each of the primary analyses (where at least 10 studies were included, as recommended in the Cochrane Handbook (16)) in order to investigate the possibility of publication bias.

#### Secondary analyses

The secondary analyses compared risks in the following groups:

- risk of ATDH between genotype groups for genetic variants other than variants of the *NAT2*, *CYP*, *GSTM1* and *GSTT1* genes (henceforth referred to as “other genetic variants”)
- risk of other anti-TB drug-related toxicity outcomes between genotype groups for *GSTM1* and *GSTT1* null polymorphisms
- risk of other anti-TB drug-related toxicity outcomes between genotype groups for other genetic variants.

We performed meta-analyses for all the above associations that were investigated by at least two studies. For single-nucleotide polymorphisms (SNPs) investigated by one study only, ORs comparing genotype groups were calculated and summarised in a table, together with the pooled estimates from the meta-analyses. 95% confidence intervals (CIs) were reported for all ORs.

For SNPs where all studies presented data for each genotype group separately, we performed two pairwise comparisons; heterozygous genotype versus homozygous wild-type, and homozygous mutant-type versus homozygous wild-type. For SNPs where all studies presented data for combined genotype groups, we performed one comparison of the combined genotype groups. For SNPs where some studies presented data for each genotype group separately, and some studies presented data for combined genotype groups, we performed both pairwise comparisons (using data from studies that reported on each genotype group separately), and a comparison of the combined genotype groups (using data from all studies).

All meta-analyses were performed using the metan package in Stata 14 (19), using the random-effects model, as we expected to observe heterogeneity between studies due to differences in study design, quality of methods, ethnic background of participants, and outcome definitions. The random-effects model used the method of DerSimonian and Laird (20) with the estimate of heterogeneity being taken from the Mantel–Haenszel model (21). If zero events were observed in one of the genotype groups, a continuity correction of 0.5 was used. Data were excluded from the analysis if there were no patients in one of the genotype groups in a comparison.

The HuGENet HuGE Review Handbook recommends that meta-analyses of genetic association studies are stratified by ethnicity, and that pooling of results should only be performed if effect estimates for different ethnic groups appear sufficiently similar (17). Information on participants’ ethnicity was not commonly reported; however, in an attempt to follow this recommendation, we stratified our analyses by the countries in which studies were performed.

## Results

### Included and excluded studies

A PRISMA flow diagram, showing the selection process of studies during the literature search, is provided in Fig. 1. The search identified 77 eligible articles, from which 52 distinct cohorts of patients were identified.

**Fig. 1.**
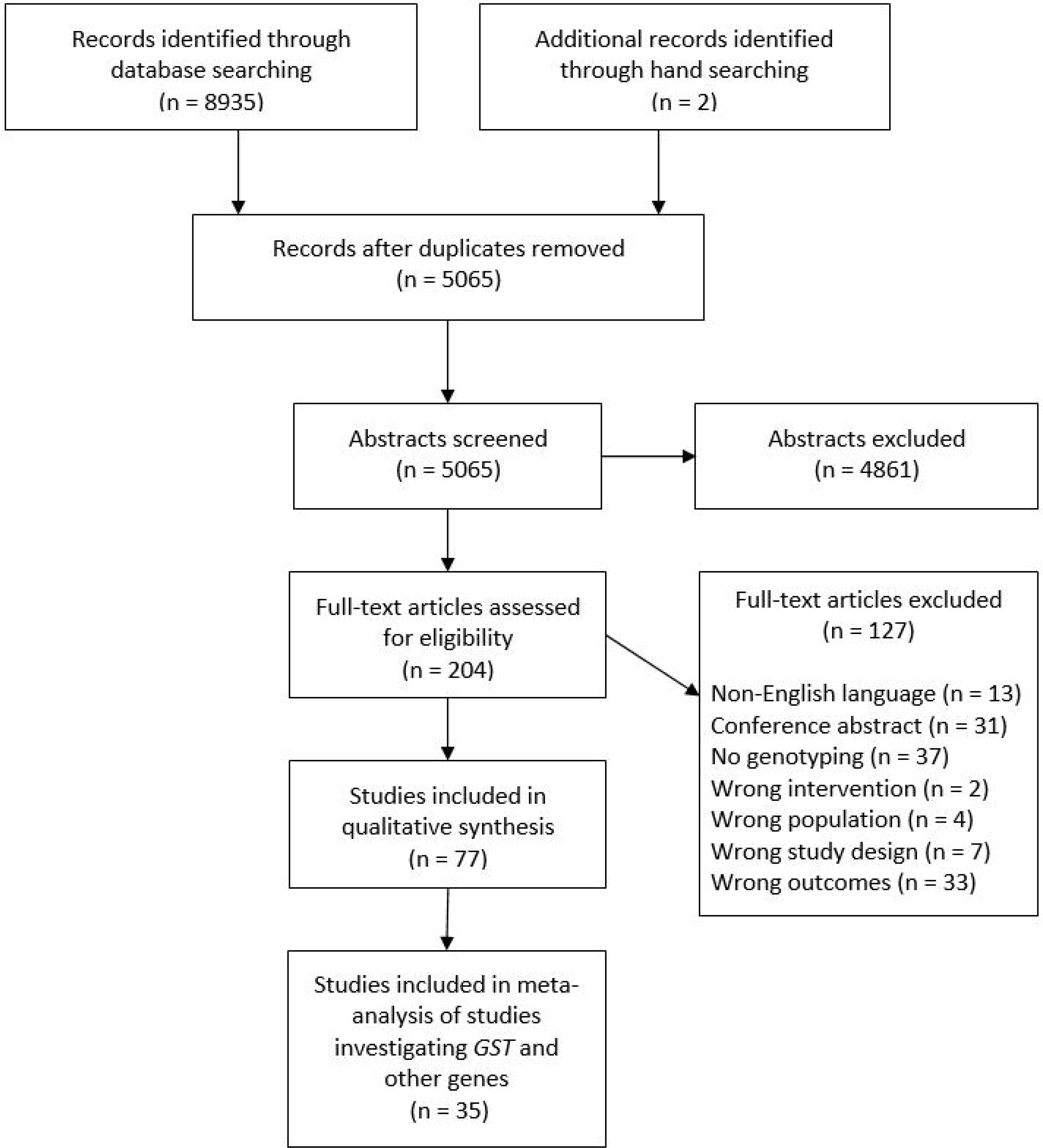
PRISMA flow diagram. From: Moher D, Liberati A, Tetzlaff J, Altman DG, The PRISMA Group. Preferred reporting items for systematic reviews and meta-analyses: the PRISMA statement. PLoS Med 2009;6(7):e1000097. doi:10.1371/journal.pmed1000097

Thirty-eight articles (29 distinct patient cohorts) investigated the association between *GSTM1/GSTT1* or other genetic variants and anti-TB drug-related toxicity. In this review, we include data from 35 articles (22–56), which included 28 distinct patient cohorts. We did not include data from two articles (57, 58) as the data presented were unclear and we were unable to clarify the data with the authors. We did not include data from a third article (59) because for the *GSTM1/GSTT1* gene, we suspected that data were reported for the same cohort of patients as that included in a more recent article by the same author (43). Data on other genetic variants presented in this third excluded article were unclear and we were unable to clarify the data with the authors. The characteristics of studies included in this review are provided in Additional file 3.

### Quality assessment

#### Choosing which genes and SNPs to genotype

Twenty-four articles provided a rationale for the choice of genes and SNPs to be investigated. Eleven articles (7, 22, 30, 43, 44, 46, 48, 50, 54–56) did not provide such information; however, none of these articles limited their reporting to only statistically significant associations. Consequently, there is no evidence to suggest that selective reporting of genes and SNPs is an issue of concern for the included studies.

#### Sample size

The median sample size was 245 (interquartile range 163–346), meaning that most studies are likely to be at risk of being underpowered (18). Only one article (40) reported the a priori power to detect pre-specified effect sizes.

#### Study design

Twenty-two articles employed a case–control design (7, 24, 25, 27, 29–38, 40, 44, 45, 48–52), and 13 articles employed a prospective cohort design (22, 23, 26, 28, 41–43, 46, 47, 53–56). Only one (29) of the 22 articles describing case-control studies reported that the two groups were genotyped in mixed batches. Separate genotyping in cases and controls could potentially bias the results (18).

#### Reliability of genotypes

Only eight articles mentioned genotype quality control procedures (7, 25, 26, 29, 30, 48, 51, 52), and therefore genotyping results from the remaining 27 articles should be interpreted with caution due to the risk of bias from incorrect genotype allocation. Only six articles (24, 32, 35, 41, 43, 47) compared genotype frequencies of all investigated SNPs with those previously published for the same population, a simple way of highlighting problems with genotyping. Only five (25, 29, 48, 51, 52) of the 22 articles describing case–control studies mentioned that genotyping personnel were blinded to outcome status; blinding minimises the risk of bias during the genotyping procedure (18).

#### Missing genotype data

For most articles (21/35, 60%), the number of participants analysed matched the study sample size; therefore, there were no missing genotype data. For the remaining 14 articles (25, 26, 31, 34–37, 45, 49, 51–55), only five articles (26, 45, 49, 53, 55) reported the extent of missing data for all genes and SNPs analysed. None of these articles described checking whether missing data were randomly distributed; therefore, 14 articles were at risk of bias from non-random missing data (18).

#### Population stratification

No articles mentioned undertaking tests for population stratification. One article reported that only patients who had been settled in the area for a minimum of three generations were eligible for inclusion in the study (46). This design ensured that all study patients were from a non-diverse ethnic group. All other studies were at risk of confounding due to population stratification.

#### Hardy–Weinberg equilibrium (HWE)

Nineteen articles (7, 23, 25–27, 30, 31, 34–37, 42, 48, 51–56) reported testing for HWE for all SNPs investigated, and a further two (43, 49) tested for HWE for a subset of SNPs. The remaining 14 articles did not report on HWE testing.

#### Mode of inheritance

Sixteen articles made a specific assumption regarding the underlying mode of inheritance (22, 24, 27–29, 32, 33, 36, 38, 41, 44, 45, 47, 49, 54, 55). Of these, only four provided a rationale for this assumption (28, 44, 45, 54); for the remaining 12 articles, there is a risk of selective reporting where several analyses under different modes of inheritance may have been conducted, with only the most statistically significant being reported (18). Six articles conducted analyses assuming different modes of inheritance (25, 31, 42, 51–53), although only three of these articles (25, 31, 51) adjusted these analyses for multiplicity of testing; therefore, there is a risk of inflating the type I error rate in the other two articles.

#### Choice and definition of outcomes

There was large variation in the definition of hepatotoxicity; a table detailing the range of definitions is available in Additional file 4. Of the 33 articles reporting hepatotoxicity data, three provided only vague definitions, and the remaining 30 articles provided 22 different definitions.

Definitions of other toxicity outcomes reported are provided in Additional file 5. These definitions were generally not sufficiently detailed.

Five articles (25, 26, 30, 41, 49) did not provide justification for the choice of outcomes, but the choice of outcomes was appropriate to address the main study aim as described in the article introduction. The remaining articles all provided justification for the choice of outcomes. Consequently, there is no evidence of selective outcome reporting.

#### Treatment adherence

Only five articles (26, 33, 34, 36, 38) mentioned assessing treatment adherence. It was not necessary to adjust for adherence in the analyses of one article, as patients were reported to have good treatment adherence (26), and the other four articles that assessed treatment adherence all excluded patients who were reported to have poor adherence. Three articles (24, 46, 56) reported that treatment was administered by directly observed therapy, short-course, so it was unnecessary to measure adherence.

### Association between *GSTM1/GSTT1* and other genetic variants and anti-TB drug-related toxicity

#### *Primary analyses*: GSTM1/GSTT1 *and hepatotoxicity*

Forest plots displaying the results of the analyses for *GSTM1* and *GSTT1* are provided in Fig. 2 and Fig. 3, respectively.

**Fig. 2.**
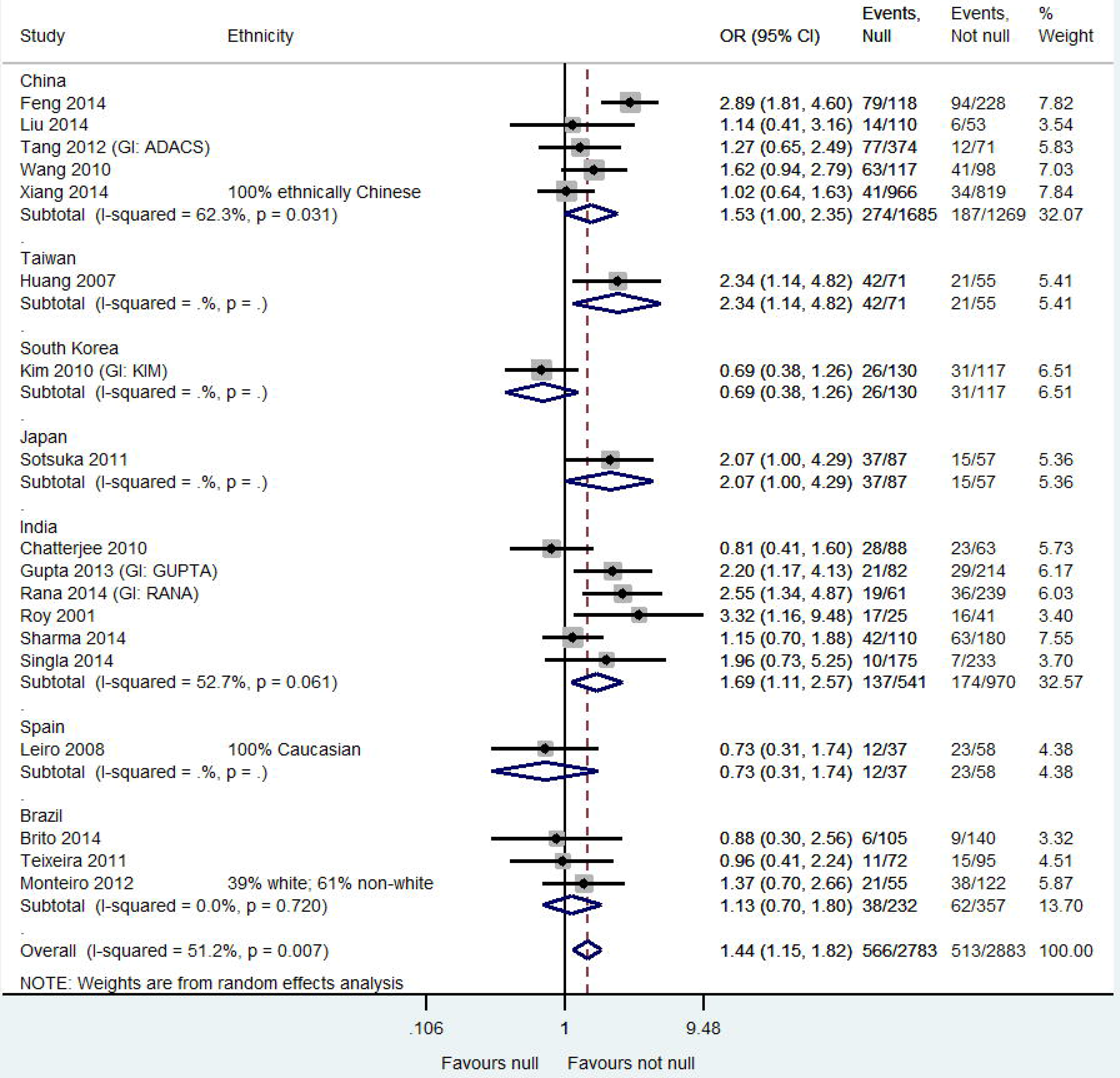
*GSTM1* null polymorphism and anti-TB drug-induced hepatotoxicity: homozygous null versus heterozygous or homozygous present genotype CI: confidence interval; GI: group identifier; OR: odds ratio

For *GSTM1*, patients with homozygous null genotype were significantly more likely to experience hepatotoxicity than patients with heterozygous or homozygous present genotype (OR=1.44, 95% CI 1.15, 1.82). Moderate heterogeneity was observed in this analysis (I^2^=51.2%). This heterogeneity may be due to the variable genotype frequencies in different geographic areas; Polimanti et al. (60) identified that the frequency of *GSTM1* null genotype ranges from 33.4% in African populations to 54.3% in European populations.

The results of the sensitivity analyses (two pairwise comparisons) for the *GSTM1* gene are provided in Additional file 6. As only one study reported on each genotype group separately for the *GSTM1* gene, no meta-analysis was performed. Instead, we calculated ORs and corresponding 95% CIs for this study for each comparison. No significant differences were observed for either pairwise comparison (heterozygous versus homozygous present: OR=0.42, 95% CI 0.02, 8.18; homozygous null versus homozygous present: OR=0.97, 95% CI 0.35, 2.71).

**Fig. 3.**
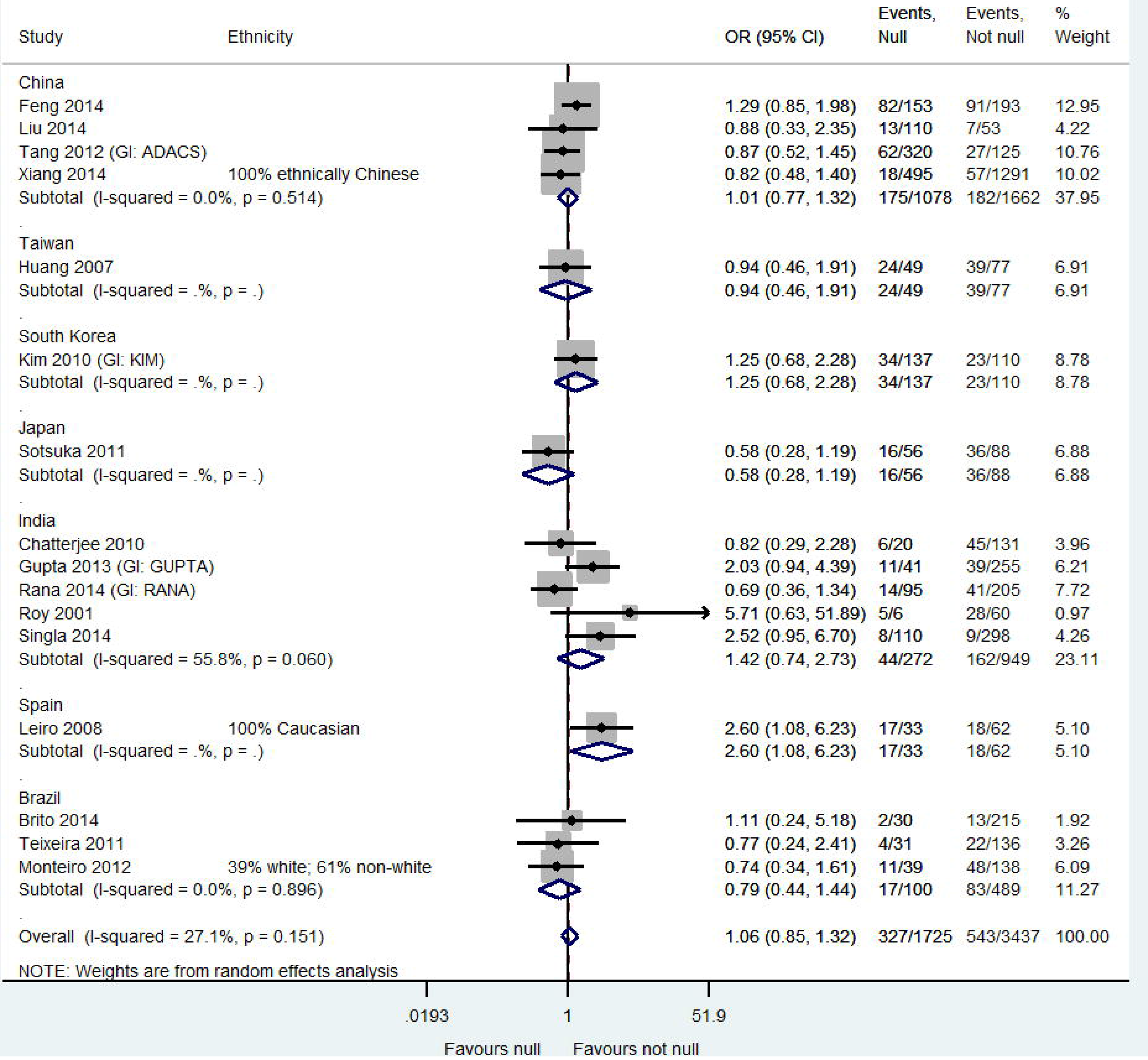
*GSTT1* null polymorphism and anti-TB drug-induced hepatotoxicity: homozygous null versus heterozygous or homozygous present genotype CI: confidence interval; GI: group identifier; OR: odds ratio

For *GSTT1*, there was no significant difference in the risk of hepatotoxicity between patients with homozygous null genotype and patients with heterozygous or homozygous present genotype (OR=1.06, 95% CI 0.85, 1.32). A relatively small amount of heterogeneity was observed in this analysis (I^2^=27.1%).

The sensitivity analyses (two pairwise comparisons) also showed no significant differences between genotype groups (heterozygous versus homozygous present: OR=0.67, 95% CI 0.29, 1.55; I^2^=0.0%; homozygous null versus homozygous present: OR=0.61, 95% CI 0.22,; I^2^=0.0%) (Additional file 6).

We produced a funnel plot for each of the primary analyses (Additional file 7). There was no evidence to suggest that publication bias was an issue of concern.

#### Secondary analyses: Other genetic variants and hepatotoxicity

The included studies reported data for 27 other genes, and 83 SNPs. A summary of all data for the association between other genetic variants and hepatotoxicity is provided in Additional file 8.

There were sufficient data to perform meta-analyses for 14 SNPs of six different genes. Forest plots showing the results of these meta-analyses are provided in Additional file 9. The findings from these meta-analyses are:

- For the rs1045642 SNP of the *ABCB1 gene*, no significant differences were observed for either pairwise comparison.
- For four SNPs of the *PXR* gene (rs3814055, rs2461823, rs7643645, rs6785049), no significant differences were observed for either pairwise comparison.
- For the rs12488820 polymorphism of the *PXR* gene, no significant difference was observed for homozygous mutant-type versus homozygous wild-type. No meta-analysis was performed for heterozygous genotype versus homozygous wild-type, as no patients had heterozygous genotype in either patient cohort.
- For the rs3814057 polymorphism of the *PXR* gene, both heterozygous genotype and homozygous mutant-type significantly increased hepatotoxicity risk compared with homozygous wild-type (heterozygous versus homozygous wild-type: OR=1.98, 95% CI 1.06, 3.69; I^2^=0%; homozygous mutant-type versus homozygous wild-type: OR=2.18, 95% CI 1.07, 4.44; I^2^=0%).
- For three SNPs of the *SLCO1B1* gene (rs4149014, rs2306283, rs4149056), no significant differences were observed for either pairwise comparison.
- For the rs4149013 polymorphism of the *SLCO1B1* gene, heterozygous genotype significantly increased hepatotoxicity risk compared with homozygous wild-type (OR=1.67, 95% CI 1.12, 2.50; I^2^=0%); however, no significant difference was observed for homozygous mutant-type versus homozygous wild-type.
- For the rs4880 polymorphism of the *SOD2* gene, no significant difference was observed for homozygous mutant-type or heterozygous genotype versus homozygous wild-type.
- For the rs4148323 polymorphism of the *UGT1A1* gene, no significant difference was observed for heterozygous genotype versus homozygous wild-type. No meta-analysis was performed for homozygous mutant-type versus homozygous wild-type, as only one study identified patients with homozygous mutant-type genotype.

For the analyses of the rs1045642 SNP of the *ABCB1* gene, and the rs4149056 and rs4149014 SNPs of the *SLCO1B1* gene, substantial heterogeneity was observed; this may be due the variable frequencies of genotypes across different geographical areas. Indeed, allele frequencies have been found to vary considerably between different ethnic populations for the rs1045642 SNP (61), for the rs4149056 SNP (also known as SLCO1B1 521T-C) (62) and the rs4149014 SNP (63).

#### Secondary analyses: GSTM1/GSTT1 and other toxicity outcomes

Data for the association between *GST* and toxicity outcomes (other than hepatotoxicity) are summarised in Additional file 10. Each reported result is based on data from a single study.

One study (Costa, 2012) investigated the association between *GSTM1/GSTT1* and adverse drug reactions (ADRs), and identified no significant association between *GSTM1* null status and ADRs, or between *GSTT1* null status and ADRs. Another study (Kim, 2010 [GI: KIM]) investigated the association between *GSTM1/GSTT1* and anti-tuberculosis drug (ATD)-induced cutaneous reactions; no significant association was identified between *GSTM1* null status and ATD-induced cutaneous reactions, or between *GSTT1* null status and ATD-induced cutaneous reactions.

#### Secondary analyses: Other genetic variants and other toxicity outcomes

Data for the association between other genetic variants and toxicity outcomes (other than hepatotoxicity) are summarised in Additional file 10. Each reported result is based on data from a single study.

One study (Kim 2012a [GI: KIM]) investigated the association between polymorphisms of the *ABCB1* and *ABCC2* genes and ATD-induced MPE. For the rs1045642 and rs10261685 polymorphisms of the *ABCB1* gene, no significant differences were observed for either pairwise comparison. For the 1774 G-del, rs717620, rs2273697, rs3740070 and rs3740066 polymorphisms of the *ABCC2* gene, no significant differences were observed for either pairwise comparison. For the rs1885301 and rs2804400 polymorphisms of the *ABCC2* gene, homozygous mutant-type was shown to significantly increase the risk of ATD-induced MPE compared with homozygous wild-type (rs1885301: OR=3.39, 95% CI 1.13, 10.14; rs2804400: OR=3.47, 95% CI 1.16, 10.36); no significant difference was observed for heterozygous genotype versus homozygous wild-type for either polymorphism.

## Discussion

### Meta-analyses

Our systematic review and meta-analysis has demonstrated that performing robust synthesis of the evidence base for associations between genetic variants of *GST* and other genes and toxicity outcomes related to anti-TB drugs is challenging. The included studies varied in terms of how participants were classified according to genotype, choice and definition of outcomes and variants, ethnicity of participants and methodological quality. While conducting our review, we carefully considered these challenges, stratifying meta-analyses by genetic variants, the combination of different genotype groups and outcomes. We also stratified further by the country in which the study was conducted as a proxy for ethnicity, which was not widely reported.

We found that for *GSTM1*, patients with homozygous null genotype were significantly more likely to experience hepatotoxicity than patients with heterozygous or homozygous present genotype. We also found that for *GSTT1*, there was no significant difference in the risk of hepatotoxicity between patients with homozygous null genotype and those with heterozygous or homozygous present genotype. These results are consistent with the findings of previously conducted meta-analyses, which all identified a significant association between *GSTM1* null genotype and hepatotoxicity, and no significant association between *GSTT1* null genotype and hepatotoxicity (4, 9–12). In particular, Cai et al. (4) and Tang et al. (12) reported very similar ORs for both the *GSTM1* and *GSTT1* genes to those reported in the current review.

We also identified that for the rs3814057 polymorphism of the *PXR* gene, both the heterozygous and homozygous mutant-type genotype significantly increased hepatotoxicity risk compared with the homozygous wild-type genotype.

### Quality assessment

Several areas of concern with regard to the quality of the included studies were identified. Most studies were significantly smaller than typically required to provide sufficient power to detect a genetic association (18); however, there was an almost universal lack of reporting of a priori power calculations (only one study reported this in the current review), and therefore readers would not be aware of the possibility of false-negative results. In addition, 77% of studies did not report on steps taken to ensure correct genotype allocation, suggesting that results should be interpreted with caution. None of the studies reported on whether the distribution of missing data had been checked. Data that are missing in a non-random fashion can introduce bias to a study. Furthermore, heterozygous genotypes are particularly difficult to identify compared with homozygous genotypes missing genotype data are not missing at random (18).

Most of the studies (34/35 [97%]) were at risk of confounding from population stratification, as they did not report on steps taken to ensure non-diverse patient ethnicity, or adjust for population stratification. Furthermore, 40% of the included studies did not report on whether HWE testing had been conducted; HWE testing can expose irregularities in genotype frequencies that could be caused by genotyping errors, population stratification and other problems (18). Nearly a third of studies (31%) did not justify their choice of mode of inheritance and may therefore be at risk of selective reporting. Finally, 77% of studies did not adjust for treatment adherence and therefore the proportion of variability explained by genetic variants may be underestimated in these studies (18).

Despite these concerns regarding quality, none of the studies were considered to be so poor methodologically to be excluded from sensitivity analyses.

### Limitations

The ethnicity of included participants was not reported widely in individual studies and this is an important limitation of the review, as the distributions of *GSTM1* and *GSTT1* genotypes vary considerably between different ethnic populations (64). As a proxy for ethnicity, we stratified our meta-analyses by the study country, making the assumption that multiple studies from a single country were likely to be relatively comparable in terms of the ethnicity of the included patients. However, this approach is not ideal, as the population of any given country is often ethnically diverse, and consequently, we are unable to make inferences about the impact of ethnicity on the associations between genetic variants of *GST* and other genes and anti-TB drug-related toxicity.

Definitions of hepatotoxicity varied substantially among the included studies (22 different definitions across 33 articles), introducing heterogeneity into the meta-analyses. Jorgensen et al. (65) and Contopoulos-Ioannidis et al. (66) made similar observations about the variability of definitions of outcomes across pharmacogenetics studies. Greater comparability of pharmacogenetic studies would help to reduce heterogeneity and increase the robustness of findings in meta-analyses. Experts in the field of TB have an important role to play in developing consensus definitions for outcomes that are commonly reported in pharmacogenetic studies of anti-TB drugs.

In most of the studies, patients were treated with a combination of anti-TB drugs, meaning that it is very difficult to link pharmacogenomic factors to specific drugs. It is possible that some studies included patients with rifampicin- or pyrazinamide-induced hepatotoxicity, for which biological mechanisms are unknown (6). If genetic variants of the investigated genes do not contribute to rifampicin- or pyrazinamide-induced hepatotoxicity, the inclusion of patients with rifampicin- or pyrazinamide-induced hepatotoxicity may have reduced the likelihood of observing significant associations.

An additional challenge was identifying distinct patient cohorts from the included articles. If multiple articles report data for the same patient cohort, data for this patient cohort must only be included in meta-analysis once, otherwise participants are “double-counted”, and the pooled effect will be overly biased toward these outcome data, and the assumption about independence between cohorts will also be violated. For two articles (43, 59), we contacted study authors for clarification about patient cohort overlap but did not receive a response. Therefore, data from the older article (59) were excluded from meta-analyses to which both articles contributed data. If the two articles reported data for distinct patient cohorts, then data has been lost by excluding one article. Furthermore, there is a possibility that there were cases of patient cohort overlap that we did not identify; if this is the case, some patients may be double-counted in the meta-analyses.

Finally, there is a lack of evidence in our review from Africa (only one study was conducted in Africa (55)). TB is endemic throughout the African continent and genotype frequencies of the *GSTM1* and *GSTT1* genes vary considerably across the region (67); however, mapping of pharmacogenomic polymorphisms is scant in African populations. Therefore, the evidence gathered and synthesised in this review is not representative of the global population most affected by TB. More pharmacogenetic studies are required in this geographic setting in order to better understand the association between genetic variants and anti-TB drug-related toxicity in African populations.

### Conclusions

We observed significant associations between the *GSTM1* null and *PXR* rs3814057 polymorphisms and ATDH. To the best of our knowledge, no meta-analyses on genetic variants other than variants of the *NAT2*, *CYP2E1*, *GSTM1* and *GSTT1* genes have been published, and therefore our results add to the existing understanding of the association between genetic variants and hepatotoxicity. A stratified medicine approach for the treatment of TB, by which patients would receive treatment according to genetic factors, would allow the risk-benefit ratio to be improved, therefore improving patient outcome and reducing healthcare costs. Although the findings from our meta-analyses alone are not sufficiently strong to support a stratified approach, they suggest, particularly for the *GSTM1* gene, that comprehensive genotyping in a wider range of populations is needed to determine the value of pharmacogenetics testing in the treatment of TB.

## List of abbreviations

ADR: adverse drug reaction
ATD: anti-tuberculosis drug
ATDH: anti-TB drug-induced hepatotoxicity
CI: confidence interval
GI: group identifier
GSTM1: glutathione s-transferase mu 1
GSTT1: glutathione S-transferase theta-1
MPE: maculopapular eruption
MT: mutant-type
N/A: not applicable
NR: not reported
OR: odds ratio
RCT: randomised controlled trial
SNP: single-nucleotide polymorphism
TB: tuberculosis
WT: wild-type

## Declarations

### Ethics approval and consent to participate

Not applicable

### Availability of data and material

The datasets used and/or analysed during the current study are available from the corresponding author on reasonable request.

### Authors’ contributions

MR is the guarantor, and drafted the manuscript. All authors contributed to development of the objectives for the review, the search strategy and selection criteria. JK and KD provided statistical expertise on meta-analysis methodology. DS and GD provided expertise on the pharmacogenetics of anti-tuberculosis drugs, and the disease area in general. AJ provided statistical expertise on pharmacogenetics studies, and contributed to the development of meta-analysis methodology tailored for pharmacogenetics studies, i.e. the quality assessment checklist. All authors read, provided feedback and approved the final manuscript.

## Acknowledgements

We would like to thank Eleanor Kotas for her assistance in drafting and implementing the search strategy.

