## Additional File 2 for "Association of variants within the *GST* and other genes with anti-tubercular agents related toxicity: a systematic review and meta-analysis"

### Additional file 2 Search history

| **Databases** | **Date searched** | **No. retrieved** |
| --- | --- | --- |
| MEDLINE (Ovid) and MEDLINE In-Process (Ovid) | 03/03/2016 | 3029 |
| EMBASE (Ovid) | 03/03/2016 | 4778 |
| PubMed | 03/03/2016 | 379 |
| Web of Science | 03/03/2016 | 421 |
| BIOSIS | 03/03/2016 | 328 |

*Search strategies*

**Database: Web of Science and BIOSIS**

| [618](http://apps.webofknowledge.com.liverpool.idm.oclc.org/summary.do?product=UA&doc=1&qid=31&SID=Y144YhJlA9U9SPEzDH1&search_mode=CombineSearches&update_back2search_link_param=yes) | #11 AND #9 AND #8 |
| --- | --- |
| [Approximately](http://apps.webofknowledge.com.liverpool.idm.oclc.org/summary.do?product=UA&doc=1&qid=30&SID=Y144YhJlA9U9SPEzDH1&search_mode=CombineSearches&update_back2search_link_param=yes)  [1,634,627](http://apps.webofknowledge.com.liverpool.idm.oclc.org/summary.do?product=UA&doc=1&qid=30&SID=Y144YhJlA9U9SPEzDH1&search_mode=CombineSearches&update_back2search_link_param=yes) | #10 OR #7 OR #6 OR #5 OR #4 |
| [Approximately](http://apps.webofknowledge.com.liverpool.idm.oclc.org/summary.do?product=UA&doc=1&qid=29&SID=Y144YhJlA9U9SPEzDH1&search_mode=GeneralSearch&update_back2search_link_param=yes)  [9,565](http://apps.webofknowledge.com.liverpool.idm.oclc.org/summary.do?product=UA&doc=1&qid=29&SID=Y144YhJlA9U9SPEzDH1&search_mode=GeneralSearch&update_back2search_link_param=yes) | **TITLE:** ((((Genetic or gene*) near/2 associat* near/2 (studies or study or analys*)))) |
| [Approximately](http://apps.webofknowledge.com.liverpool.idm.oclc.org/summary.do?product=UA&doc=1&qid=27&SID=Y144YhJlA9U9SPEzDH1&search_mode=CombineSearches&update_back2search_link_param=yes)  [93,935](http://apps.webofknowledge.com.liverpool.idm.oclc.org/summary.do?product=UA&doc=1&qid=27&SID=Y144YhJlA9U9SPEzDH1&search_mode=CombineSearches&update_back2search_link_param=yes) | #2 OR #1 |
| [Approximately](http://apps.webofknowledge.com.liverpool.idm.oclc.org/summary.do?product=UA&doc=1&qid=26&SID=Y144YhJlA9U9SPEzDH1&search_mode=GeneralSearch&update_back2search_link_param=yes)  [382,014](http://apps.webofknowledge.com.liverpool.idm.oclc.org/summary.do?product=UA&doc=1&qid=26&SID=Y144YhJlA9U9SPEzDH1&search_mode=GeneralSearch&update_back2search_link_param=yes) | **TITLE:** ((((TB or Tuberculosis* or Antitubercul*)))) |
| [Approximately](http://apps.webofknowledge.com.liverpool.idm.oclc.org/summary.do?product=UA&doc=1&qid=25&SID=Y144YhJlA9U9SPEzDH1&search_mode=GeneralSearch&update_back2search_link_param=yes)  [219,324](http://apps.webofknowledge.com.liverpool.idm.oclc.org/summary.do?product=UA&doc=1&qid=25&SID=Y144YhJlA9U9SPEzDH1&search_mode=GeneralSearch&update_back2search_link_param=yes) | **TITLE:** ((((gene* or genetic*) near/5 (mutat* or variant*)))) |
| [Approximately](http://apps.webofknowledge.com.liverpool.idm.oclc.org/summary.do?product=UA&doc=1&qid=24&SID=Y144YhJlA9U9SPEzDH1&search_mode=GeneralSearch&update_back2search_link_param=yes)  [1,388,831](http://apps.webofknowledge.com.liverpool.idm.oclc.org/summary.do?product=UA&doc=1&qid=24&SID=Y144YhJlA9U9SPEzDH1&search_mode=GeneralSearch&update_back2search_link_param=yes) | **TITLE:** ((((SNP or Genotyp* or Phenotyp* or Allele* or Pharmacogenet* or Pharmacogenom* or Polymorph*)))) |
| [Approximately](http://apps.webofknowledge.com.liverpool.idm.oclc.org/summary.do?product=UA&doc=1&qid=23&SID=Y144YhJlA9U9SPEzDH1&search_mode=GeneralSearch&update_back2search_link_param=yes)  [38,430](http://apps.webofknowledge.com.liverpool.idm.oclc.org/summary.do?product=UA&doc=1&qid=23&SID=Y144YhJlA9U9SPEzDH1&search_mode=GeneralSearch&update_back2search_link_param=yes) | **TITLE:** (((single* near/2 nucleotid* near/2 polymorph*))) |
| [Approximately](http://apps.webofknowledge.com.liverpool.idm.oclc.org/summary.do?product=UA&doc=1&qid=22&SID=Y144YhJlA9U9SPEzDH1&search_mode=GeneralSearch&update_back2search_link_param=yes)  [45,745](http://apps.webofknowledge.com.liverpool.idm.oclc.org/summary.do?product=UA&doc=1&qid=22&SID=Y144YhJlA9U9SPEzDH1&search_mode=GeneralSearch&update_back2search_link_param=yes) | **TITLE:** ((((genetic* or gene*) near/3 (suscept* or predisposit* or anticipat*)))) |
| [Approximately](http://apps.webofknowledge.com.liverpool.idm.oclc.org/summary.do?product=UA&doc=1&qid=20&SID=Y144YhJlA9U9SPEzDH1&search_mode=GeneralSearch&update_back2search_link_param=yes)  [47,961](http://apps.webofknowledge.com.liverpool.idm.oclc.org/summary.do?product=UA&doc=1&qid=20&SID=Y144YhJlA9U9SPEzDH1&search_mode=GeneralSearch&update_back2search_link_param=yes) | **TITLE:** (((aminosalicylic acid or diarylquinoline* or ethambutol* or ethionamide* or isoniazid* or prothionamide* or pyrazinamide* or thioacetazone* or capreomycin* or cycloserine* or enviomycin* or rifabutin* or rifampin* or viomycin*))) |
| [Approximately](http://apps.webofknowledge.com.liverpool.idm.oclc.org/summary.do?product=UA&doc=1&qid=19&SID=Y144YhJlA9U9SPEzDH1&search_mode=GeneralSearch&update_back2search_link_param=yes)  [49,386](http://apps.webofknowledge.com.liverpool.idm.oclc.org/summary.do?product=UA&doc=1&qid=19&SID=Y144YhJlA9U9SPEzDH1&search_mode=GeneralSearch&update_back2search_link_param=yes) | **TITLE:** ((((Antitubercul* or tuberculos* or TB) Near/4 (agent* or drug* or antibiotic* or medicine* or medication* or treatment*)))) |

**Database: MEDLINE**

| [# ▲](http://ovidsp.tx.ovid.com/sp-3.18.0b/ovidweb.cgi?&S=OPDEFPAOGIDDHCAFNCJKEHFBGBANAA00&Sort+Sets=descending) | **Searches** | **Results** |
| --- | --- | --- |
| 1 | antitubercular agents/ or aminosalicylic acid/ or diarylquinolines/ or ethambutol/ or ethionamide/ or isoniazid/ or prothionamide/ or pyrazinamide/ or thioacetazone/ or antibiotics, antitubercular/ or capreomycin/ or cycloserine/ or enviomycin/ or rifabutin/ or rifampin/ or viomycin/ | 73943 |
| 2 | ((Antitubercul* or tuberculos* or TB) adj4 (agent* or drug* or antibiotic* or medicine* or medication* or treatment*)).tw. | 29293 |
| 3 | (aminosalicylic acid or diarylquinoline* or ethambutol* or ethionamide* or isoniazid* or prothionamide* or pyrazinamide* or thioacetazone* or capreomycin* or cycloserine* or enviomycin* or rifabutin* or rifampin* or viomycin*).tw. | 26053 |
| 4 | 1 or 2 or 3 | 93357 |
| 5 | Polymorphism, Genetic/ | 103705 |
| 6 | genetic predisposition to disease/ or anticipation, genetic/ | 101390 |
| 7 | Pharmacogenetics/ | 9595 |
| 8 | Genetic Association Studies/ | 14210 |
| 9 | ((Genetic or gene*) adj2 associat* adj2 (studies or study or analys*)).tw. | 4883 |
| 10 | ((genetic* or gene*) adj3 (suscept* or predisposit* or anticipat*)).tw. | 40247 |
| 11 | Polymorphism, Single Nucleotide/ | 77811 |
| 12 | (single* adj2 nucleotid* adj2 polymorph*).tw. | 46260 |
| 13 | (SNP or Genotyp* or Phenotyp* or Allele* or Pharmacogenet* or Pharmacogenom* or Polymorph*).tw. | 774469 |
| 14 | ((gene* or genetic*) adj5 (mutat* or variant*)).tw. | 182197 |
| 15 | Genotype/ or Phenotype/ or Alleles/ | 381555 |
| 16 | or/5-15 | 1035512 |
| 17 | exp Tuberculosis/ | 175110 |
| 18 | (TB or Tuberculosis*).tw. | 153175 |
| 19 | Antitubercul*.tw. | 11635 |
| 20 | or/17-19 | 213138 |
| 21 | 4 and 16 and 20 | 2846 |
| 22 | animal/ not human/ | 4159388 |
| **23** | **21 not 22** | **2730** |

**Database: EMBASE**

| [# ▲](http://ovidsp.uk.ovid.com/sp-3.25.0a/ovidweb.cgi?&S=POIHPDPCNBHFIMKMFNGKPFDGBNGJAA00&Sort+Sets=descending) | Searches | Results |
| --- | --- | --- |
| 1 | antitubercular agents/ or aminosalicylic acid/ or diarylquinolines/ or ethambutol/ or ethionamide/ or isoniazid/ or prothionamide/ or pyrazinamide/ or thioacetazone/ or antibiotics, antitubercular/ or capreomycin/ or cycloserine/ or enviomycin/ or rifabutin/ or rifampin/ or viomycin/ | 151901 |
| 2 | ((Antitubercul* or tuberculos* or TB) adj4 (agent* or drug* or antibiotic* or medicine* or medication* or treatment*)).tw. | 40664 |
| 3 | (aminosalicylic acid or diarylquinoline* or ethambutol* or ethionamide* or isoniazid* or prothionamide* or pyrazinamide* or thioacetazone* or capreomycin* or cycloserine* or enviomycin* or rifabutin* or rifampin* or viomycin*).tw. | 34743 |
| 4 | 1 or 2 or 3 | 172588 |
| 5 | Polymorphism, Genetic/ | 102257 |
| 6 | genetic predisposition to disease/ or anticipation, genetic/ | 97585 |
| 7 | Pharmacogenetics/ | 17431 |
| 8 | Genetic Association Studies/ | 876 |
| 9 | ((Genetic or gene*) adj2 associat* adj2 (studies or study or analys*)).tw. | 7890 |
| 10 | ((genetic* or gene*) adj3 (suscept* or predisposit* or anticipat*)).tw. | 61544 |
| 11 | Polymorphism, Single Nucleotide/ | 98303 |
| 12 | (single* adj2 nucleotid* adj2 polymorph*).tw. | 75841 |
| 13 | (SNP or Genotyp* or Phenotyp* or Allele* or Pharmacogenet* or Pharmacogenom* or Polymorph*).tw. | 1171894 |
| 14 | ((gene* or genetic*) adj5 (mutat* or variant*)).tw. | 294715 |
| 15 | Genotype/ or Phenotype/ or Alleles/ | 777386 |
| 16 | or/5-15 | 1548879 |
| 17 | exp Tuberculosis/ | 197008 |
| 18 | (TB or Tuberculosis*).tw. | 187590 |
| 19 | Antitubercul*.tw. | 16330 |
| 20 | or/17-19 | 253048 |
| 21 | 4 and 16 and 20 | 5380 |
| 22 | animal/ not human/ | 1357016 |
| 23 | 21 not 22 | 5360 |
| 24 | limit 23 to em=188300-201608 | 4778 |

**Database: PubMed**

| #1 | Search (((Antitubercul* or tuberculos* or TB))) AND ((agent* or drug* or antibiotic* or medicine* or medication* or treatment*)) | 124242 |
| --- | --- | --- |
| #2 | Search ((aminosalicylic acid or diarylquinoline* or ethambutol* or ethionamide* or isoniazid* or prothionamide* or pyrazinamide* or thioacetazone* or capreomycin* or cycloserine* or enviomycin* or rifabutin* or rifampin* or viomycin*)) | 49591 |
| #3 | Search (#1 or #2) | 151329 |
| #4 | Search ((((Genetic or gene*) near/2 near/2 ))) AND associat*) AND ((studies or study or analys*)) | 3922 |
| #5 | Search (((genetic* or gene*))) AND ((suscept* or predisposit* or anticipat*)) | 235548 |
| #6 | Search ((single*) AND nucleotid*) AND polymorph* | 98071 |
| #7 | Search ((SNP or Genotyp* or Phenotyp* or Allele* or Pharmacogenet* or Pharmacogenom* or Polymorph*)) | 997538 |
| #8 | Search (((gene* or genetic*))) AND ((mutat* or variant*)) | 743819 |
| #9 | Search (#4 or #5 or #6 or #7 or #8) | 1553428 |
| #10 | Search (((((TB or Tuberculosis* or Antitubercul*))))) | 251923 |
| #11 | Search (#3 and #9 and #10) | 7671 |
| #12 | Search ("2015/08/01"[Date - Entrez] : "3000"[Date - Entrez]) | 658085 |
| #13 | Search (#11 and #12) | 379 |
