## Additional File 3 for "Association of variants within the *GST* and other genes with anti-tubercular agents related toxicity: a systematic review and meta-analysis"

**Additional file 3** Key characteristics of included studies

| **Study** | **Country** | **Study design** | **Follow-up time** | **Drugs and dosage** | **Selection criteria** | **Sample size** | **Toxicity outcomes** |
| --- | --- | --- | --- | --- | --- | --- | --- |
| Brito 2014 | Brazil | Prospective cohort | NR | Daily treatment with INH, RIF, and PZA for the first 2 months, followed by INH and RIF for an additional 4 months | Inclusion criteria: patients aged 18 years and older that were newly diagnosed with active TB and were submitted to treatment as described in “Drugs and dosage”.  Exclusion criteria: patients using anti-TB drugs prior to study enrolment, patients whose results of LFTs prior to the beginning of treatment were higher than twice the ULN and refusal to participate in the study. | 245 | Drug-induced hepatitis |
| Chang 2012 | Taiwan | Prospective cohort | NR | First-line anti-TB medications | Patients diagnosed with TB and treated with first-line anti-TB medications  Exclusion criteria were pregnancy, abnormal liver function and history of positive viral hepatitis before starting treatment with first-line anti-TB drugs. | 98 | ATDH |
| Chatterjee 2010 | India | Case-control | NR | ATD regimens comprising of INH, RIF and PZA as per the Revised National Tuberculosis Control Program of India –DOTS | Inclusion criteria: newly diagnosed cases of patients with pulmonary TB, aged 12 years or more, and of both sexes were potential study subjects. All subjects who received treatment as described in “Drugs and dosage” were screened and recruited if they had (a) normal pre-treatment serum ALT, AST, bilirubin, ALP and albumin levels (b) hepatitis B surface antigen and HIV negative.  For each case, 2 controls were recruited, within the same cohort and on the same ATD (INH, RIF, PZA) but with serum ALT levels <3 times ULN, serum bilirubin <1 mg⁄dL and no history of severe nausea, vomiting within the first 3 months of initiation of therapy. Controls were matched with cases on the basis of age, sex, disease severity and drug dosage.  Exclusion criteria: pre-existing documented liver diseases like cirrhosis, acute or chronic hepatitis, and alcoholic liver disease | 151 | ATDH |
| Chen 2015 (GI: ADACS) | China | Case-control | NR | All patients took INH 600 mg, RIF 600 mg (or 450 mg if body weight was <50 kg), PZA 2000 mg and EMB 1250 mg every other day for the first 2 months (re-treatment patients were injected with SM 750 mg each time simultaneously). Then with the same regimen, PZA and EMB were discontinued for primary patients, whereas PZA and SM were discontinued for re-treatment patients, the rest of the drugs were continued for another 4 months | Inclusion criteria: newly diagnosed patients with sputum smear positive pulmonary TB  Exclusion criteria: (i) abnormal serum ALT, AST or total bilirubin levels before anti-TB treatment; (ii) carriers of the hepatitis B or C virus; (iii) alcoholic liver disease or habitual alcohol drinking; (iv) the concomitant use of hepatotoxic drugs; and (v) a history of chronic liver disease or systemic diseases that may cause liver dysfunction. | 445 | ATDH |
| Costa 2012 | Brazil | Prospective cohort | NR | All patients were treated with the first-line anti-TB drug regimen INH (300 mg/kg/day), RIF (300 mg/kg/day), and PZA (1500 mg/kg/day) for the first 2 months, and then INH and RIF for a further 4 months. | Male or female subjects aged 18 years or over, who had no previously described renal, allergic, or hepatic diseases and were not pregnant, were considered for the study. | 129 | ADRs |
| Feng 2014 | China | Case-control | 6 months | Treatment with anti-TB drug regimens at the usual dosage, including 300 mg/day INH, 450 mg/day RIF, and 1500 mg/day PZA | Selection of cases:  The cases were selected based on liver functions, i.e., all indices of liver function were normal before anti-TB chemotherapy, and became abnormal indicating hepatic injury after 6 months of chemotherapy. Cases were patients who showed anti-TB drug-induced hepatitis based on increased serum transaminase values that were 3-fold higher than the ULN (40 IU/L ALT) and symptoms compatible with hepatitis.  Selection of controls Controls underwent the same anti-TB chemotherapy with the selected cases and were not tested with abnormality in liver functions after 6 months of the chemotherapy. The controls selected matched the criteria compared to the cases: i) same gender; ii) age discrepancy of less than 5 years; iii) living in the same regions; and iv) treatment with anti-TB drug regimens at the usual dosage, including 300 mg/day INH, 450 mg/day RIF, and 1500 mg/day PZA. | 346 | ATDH |
| Gupta 2013 (GI: GUPTA) | India | Prospective cohort | 6-9 months | INH: 5 mg/kg (max. 300 mg/day)  RIF: 10 mg/kg (max. 600 mg/day)  PZA: 25 mg/kg (max. 2000 mg/day)  EMB: 15-25 mg/kg (max. 1500 mg/day) | Inclusion criteria: i) patients showing evident lesion of TB by simple X-ray or computer tomography; ii) positive sputum smear and/or culture for acid fast bacilli in clinical samples; iii) normal ALT (4-37 IU/L), AST (4-40 IU/L) and total bilirubin levels (0.2-1.6 mg/dL)  Exclusion criteria: i) clinically and laboratory confirmed chronic liver disease such as jaundice; ii) positive serological testing for hepatitis B and/or C viruses; iii) alcoholic liver disease or habitual alcohol drinking; iv) patients using anti-TB drugs prior to enrolment in the study and/or other potentially hepatotoxic drugs; v) ALT, AST, and total bilirubin levels 2-fold above the ULN before treatment; vi) refusal to participate in the study | 296 | ATDH |
| He 2015 | China | Case-control | 6 months | Daily 2S(E)HRZ4HR: S, streptomycin; E, ethambutol; H, isoniazid, R, rifampicin; Z, pyrazinamide; dose increased for 2 months and then consolidated for 4 months | Cases:  Inclusion criteria: occurrence of liver injury after 6 months of anti-TB drug therapy.  Controls:  Inclusion criteria: absence of liver injury after 6 months of anti-TB drug therapy and a match with patients in the case group in terms of age (<5 years difference), sex, and therapeutic regimen  Exclusion criteria: presence of abnormal liver function before the administration of TB treatment; co-occurrence of other diseases that can cause liver function abnormalities such as viral hepatitis, alcoholic liver disease, autoimmune hepatitis, and hypoxemia; and consumption of other drugs that can cause liver dysfunction in patients. | 254 | ADLI |
| Huang 2007 | Taiwan | Case-control | NR | NR | Cases  Inclusion criteria: (1) an increase in ALT or AST level greater than 5 times the ULN, or an elevation in ALP greater than twice the ULN, confirmed on at least 2 consecutive blood draws; (2) if baseline ALT, AST or ALP are known and elevated, then ALT or AST greater than 5 times baseline value, or ALP greater than twice baseline level on at least 2 consecutive blood draws; (3) any elevation of serum ALT, AST, or ALP, associated with an increased serum total bilirubin (>2.5 mg/dL), in the absence of prior diagnosis of liver disease, Gilbert’s syndrome, or evidence of hemolysis; (4) a causality assessment score greater than 5  Exclusion criteria: (1) refusal of blood sampling or informed written consent; (2) patients having possible acute or chronic viral hepatitis, such as positive serum hepatitis B virus surface antigen, IgM antibody to hepatitis B core antigen, IgM antibody to hepatitis A virus, antibody to hepatitis C virus, IgM antibody to Epstein–Barr virus, cytomegalovirus and herpes simplex virus; (3) patients with chronic alcoholism or fatty liver; (4) probable autoimmune liver diseases with positive serum anti-nuclear antibody or anti-mitochondrial antibody; (5) any other major hepatic or systemic diseases that may induce elevation of liver biochemical tests, such as fatty liver, alcoholic liver disease, cholangitis, congestive heart failure, hypoxia, and bacteremia; (6) acetaminophen hepatotoxicity; (7) a causality assessment score less than 5; (8) patients with incomplete clinical or laboratory data  Controls  In- and out-patients of the Taipei Veterans General Hospital, matched with cases according to sex, age (within 10 years old), drug(s), duration of therapy (equivalent to or longer); liver biochemical tests, checked regularly during the administration of drugs, were within normal limits | 230 (63 receiving anti-TB drugs had hepatotoxicity, 63 did not) | DILI |
| Kim 2009 (GI: KIM) | Korea | Case-control | NR | All patients with pulmonary TB were treated daily with a combination regimen including INH (300-400 mg daily), RFP (450-600 mg daily), EMB (600-800 mg daily) and PZA (1000-1500 mg daily) for 2 months and then without PZA for 4 or more following months. Doses of each drug were adjusted based on body weight of the patient | Newly diagnosed and treated patients with pulmonary TB.  Exclusion criteria: patients with active or chronic hepatitis including alcoholic hepatitis, fatty liver disease, liver cirrhosis, carriers of the hepatitis B or C virus, heavy alcohol intake, decreased renal function and severe cardiac diseases requiring several medications | 226 | ATDH |
| Kim 2010 (GI: KIM) | Korea | Case-control | NR | For the 2-month initial phase, patients were given INH (300–400 mg daily), RIF (450–600 mg daily), EMB (600–800 mg daily), and PZA (1000–1500 mg daily), and doses of each drug were adjusted, depending on the body weight of the subjects. PZA was omitted in the subsequent continuation phase for 4 months or more with maintenance of INH, RIF, and EMB. The period of the continuation phase in each patient was determined by the clinician, based on clinical features and treatment response. | Cases  Inclusion criteria: patients with ATD-induced hepatitis and ATD-induced adverse cutaneous reactions among patients newly diagnosed with pulmonary TB and/or TB pleuritis and treated with first-line anti-TB medications  Exclusion criteria: (1) abnormal LFT result at baseline (elevated serum AST, ALT, or bilirubin above normal range); (2) active or chronic hepatitis, including alcoholic hepatitis, fatty liver disease and liver cirrhosis; (3) carriers of the hepatitis B or C virus; (4) heavy alcohol intake; (5) decreased renal function; (6) other chronic medical conditions requiring medication; and (7) patients with skin diseases before treatment  Controls  Patients who showed no adverse reaction to ATD during the treatment period and agreed to participate in this study | 341 | ATDH  ATD-induced cutaneous reactions (and the subgroup of MPE patients) |
| Kim 2012a (GI: KIM) | Korea | Case-control | NR | All patients with pulmonary TB and/or TB pleurisy were treated daily with a combination regimen including INH (300-400 mg daily), RIF (450-600 mg daily), EMB (600-800 mg daily), and PZA (1,000-1,500 mg daily) for 2 months and then without PZA for 4 or more following months | Inclusion criteria: patients with pulmonary TB and/or TB pleurisy  Exclusion criteria: patients with skin diseases before treatment, chronic renal failure, and chronic liver diseases, including alcoholic hepatitis, fatty liver disease, liver cirrhosis, and carriers of the hepatitis B or C virus, and non-adherence to the treatment | 221 | ATD-induced MPE |
| Kim 2012b (GI: KIM) | Korea | Case-control | NR | All patients with pulmonary TB were treated with a combination regimen including INH (300-400 mg daily), RIF (450-600 mg daily), EMB (600-800 mg daily), and PZA (1000-1500 mg daily) for 2 months and then with the same regimen, without PZA, for at least 4 more months. Doses of each drug were adjusted based on the body weight of each patient | Inclusion criteria: patients newly diagnosed with pulmonary TB and/or TB pleurisy  Exclusion criteria: patients with active or chronic hepatitis including alcoholic hepatitis, fatty liver disease, liver cirrhosis, and carriers of the hepatitis B or C virus | 226 | ATDH |
| Kim 2012c (GI: KIM) | Korea | Case-control | NR | The treatment consisted of an initial phase of 2 months and a subsequent continuation phase of 4 or more months. During the initial phase, 4 drugs were administered, including INH (300 mg daily), RIF (450–600 mg daily), EMB (600–800 mg daily) and PZA (1000–1500 mg daily). Doses of each drug were adjusted based on the body weight of each patient. In the continuation phase, PZA was discontinued, whereas the other 3 drugs were continued. The period of the continuation phase in each patient was determined by the clinician, based on clinical features and treatment response | Inclusion criteria: patients with newly diagnosed pulmonary TB and/or TB pleuritis who were treated with first-line ATD, including INH, RIF, EMB and PZA  Exclusion criteria: (1) an abnormal LFT result at baseline (elevated serum AST, ALT or bilirubin above normal range); (2) active or chronic hepatitis, including alcoholic hepatitis, fatty liver disease and cirrhosis; (3) carriers of the hepatitis B or C virus; (4) heavy alcohol intake; (5) decreased renal function; (6) other chronic medical conditions requiring medication and (7) non-adherence to the treatment | 306 | ATDH |
| Kim 2015 (GI: KIM) | Korea | Case-control | NR | All patients with pulmonary TB were treated daily with a combination regimen including INH (300-400 mg daily), RIF (450-600 mg daily), EMB (600-800 mg daily), and PZA (1,000-1,500 mg daily) for 2 months, and then without PZA for 4 or more months. Doses of each drug were adjusted based on body weight of the patient | Inclusion criteria: newly treated patients with first-line ATD  Exclusion criteria: 1) an abnormal LFT result at baseline; 2) history of liver disease (active and chronic hepatitis, fatty liver disease and cirrhosis); 3) decreased renal function; 4) other serious medical conditions requiring medication; 5) non-adherence to the treatment | 321 | ATD-hepatitis |
| Kwon 2012 | Korea | Case-control | The time between the initiation of the causative drug and symptom onset was ≤14 days | Anti-TB drugs | Inclusion criteria: patients with DILI  Exclusion criteria: NR | 238 (not all on TB drugs) | DILI |
| Leiro 2008 | Spain | Case-control | NR | Treatment with regimens that included at least INH, RIF and PZA at the usual drug dosages (INH 5 mg/kg/day – maximum 300 mg/day, RIF 10 mg/kg/day – maximum 600 mg/day and PZA 25–30 mg/kg/day – maximum 2500 mg/day) | Inclusion criteria: (i) age between 15 and 75 years; (ii) microbiological demonstration of active TB; (iii) treatment as described in “Drugs and dosage”; (iv) adequate compliance  Exclusion criteria: (i) increased baseline serum transaminases (ALT or AST, normal value ≤40 IU/L); (ii) positive serological testing for HIV, hepatitis B virus or hepatitis C virus; (iii) regular alcohol intake or concomitant use of hepatotoxic drugs; (iv) a history of chronic liver disease; (v) pregnancy; (vi) default or poor adherence to treatment | 95 | ATDH |
| Li 2012 | China | Case-control | NR | RIF treatment | Inclusion criteria: i) 18-65 years of age, and ii) RIF treatment prescribed for >3 months  Exclusion criteria: pre-existing cardiovascular, renal, hepatic, hematologic or immunologic diseases | 273 | DILI |
| Liu 2014 | China | Case-control | NR | Standard anti-TB therapy: INH 10–20 mg/kg/day (up to a maximum of 300 mg/day), RIF 10–20 mg/kg/day (up to a maximum of 450 mg/day), PZA 20–30 mg/kg/day(up to a maximum of 1500 mg/day), EMB 15–25 mg/kg/day, and SM 20–30 mg/kg/day (up to a maximum of 750 mg/day) | Inclusion criteria: (i) Chinese Han children aged between 0 and 16 years; (ii) diagnosis of active TB by clinical examination, radiological and microbiological investigations; (iii) standard anti-TB treatment has been started for at least 2 weeks; (iv) serum transaminases were normal before treatment (ALT: 40 IU/L, AST: 40 IU/L)  Exclusion criteria: pre-existing liver disease, viral hepatitis, chronic alcoholism, or history of intake of other hepatotoxic drugs | 163 | ATDH |
| Monteiro 2012 | Brazil | Prospective cohort | NR | NR | Inclusion criteria: (i) signed written consent; (ii) sputum smear with acid-fast bacilli or culture positive for Mycobacterium tuberculosis; (iii) ongoing TB treatment; (iv) HIV, Hepatitis B and C virus serology results; (v) laboratory LFTs  Exclusion criteria: (i) no more than 1 visit registered; (ii) pregnancy; (iii) age <18 years | 177 | ATD-induced liver injury |
| Nanashima 2012 | Japan | Prospective Cohort | NR | Treatment including INH (400 mg/day) and RIF (450 mg/day) for 6-9 months | Inclusion criteria: new onset of pulmonary TB with treatment as described in “Drugs and dosage”  Exclusion criteria: liver cirrhosis, acute hepatitis, chronic hepatitis, alcoholic liver disease, other chronic liver diseases | 100 | ATDH |
| Rana 2014 (GI: RANA) | India | Prospective cohort | NR | Daily ATT for the first 2 months included INH (300 mg), RIF (600 or 450 mg for body weight/50 kg), PZA (20 mg/kg body weight) and EMB (25 mg/kg body weight). After 2 months, EMB and PZA were discontinued, whereas INH and RIF were continued for an additional 4 months | Inclusion criteria: Patients with pulmonary and extra-pulmonary TB  Exclusion criteria: patients with history of alcohol abuse and/or any other liver disease; patients who had received other potentially hepatotoxic drugs in addition to antitubercular drugs; patients who had abnormal serum ALT, AST or bilirubin before starting ATT; patients who developed viral hepatitis during ATT; patients with renal failure or cancer; and patients who refused blood sampling or providing informed written consent | 300 | Hepatotoxicity |
| Roy 2001 | India | Case-control | NR | Cases and controls were treated with a daily dose of the same uniform drug regimen for the first 2 months: INH (300 mg), RIF (450 mg), PZA (1.5 g) and EMB (800 mg). Subsequently, INH and RIF were continued for a further 4 months | Inclusion criteria: (i) diagnosis of ATDH based on clinical history, laboratory findings (serum bilirubin > 3.0 mg/dL and raised ALT level at least 2-fold the ULN); (ii) aged between 18 and 75 years; and (iii) informed consent from the patients.  Exclusion criteria: patients with pre-existing liver disease, evidence of viral hepatitis, chronic alcoholism, history of intake of other hepatotoxic drugs, concurrent medical illness and pregnancy | 66 | ATDH |
| Sharma 2014 | India | Case-control | NR | INH, RIF, PZA, EMB; dosages administered to patients according to body weight:  RIF, body weight (kg) and mg/day <=35: 300  36–50: 450 >50: 600  INH, body weight (kg) and mg/day <=35: 200 >35: 300  PZA, body weight (kg) and g/day <=50: 1.0 >50: 1.5  EMB, 15 mg/kg/day | Cases: patients who developed clinical and/or laboratory evidence of DIH while on ATT  Controls: TB patients without DIH  Exclusion criteria: patients whose serum samples tested positive for markers of viral hepatitis and/or who were receiving other potentially hepatotoxic drugs or had ultrasonography evidence of chronic liver disease; HIV-infected and chronic alcohol-dependent patients who had consumed >48 g of alcohol/day for at least 1 year; patients receiving other potentially hepatotoxic drugs (e.g., methotrexate, phenytoin, valproate, fluconazole); pregnant women; subjects who did not provide written informed consent | 314 | DIH |
| Singla 2014 | India | Prospective cohort | NR | NR | Inclusion criteria: newly diagnosed patients with TB  Exclusion criteria: history of heavy use of alcohol or chronic liver diseases or liver cirrhosis; patients who were settled in the area for a minimum of 3 generations; people infected with HIV | 408 | ATDH |
| Sotsuka 2011 | Japan | Prospective cohort | 3 months | INH, RIF and PZA, plus EMB or SM during the first 2 months, followed by administration of INH and RIF plus EMB or SM during the final 4 months | Inclusion criteria: inpatients with active pulmonary TB who were treated with the standard Japanese chemotherapy regimen followed up for more than 3 months after treatment, and who consented to this study | 144 | Hepatotoxicity |
| Tang 2012 (GI: ADACS) | China | Case-control | Patients were monitored for 6–9 months according to the treatment episode. | All patients took INH 600 mg, RIF 600 mg (or 450 mg if body weight was < 50 kg), PZA 2000 mg and EMB 1250 mg every other day for the first 2 months (re-treatment patients were injected with SM 750 mg each time simultaneously). PZA and EMB were then discontinued for primary patients, whereas INH and RIF were continued for another 4 months. PZA and SM were discontinued for re-treatment patients, whereas INH, RIF and EMB continued. | Inclusion criteria: sputum smear-positive patients who received standard short-course chemotherapy recommended by WHO  Exclusion criteria: patients using other potentially hepatotoxic medications; patients with positive serum hepatitis B virus surface antigen, alcohol drinking, liver diseases or abnormal liver function before ATT | 445 | ATDH |
| Teixeira 2011 | Brazil | Case-control | NR | Anti-TB drug regimens that include INH at the usual dosage (400 mg/day) | Inclusion criteria: (i) age above 18 years, (ii) diagnosis of active TB, (iii) treatment with anti-TB drug regimens that include INH at the usual dosage (400 mg/day) and (iv) normal baseline serum transaminases (ALT and AST) before treatment  Exclusion criteria: i) positive serological test for the HIV, hepatitis B virus or hepatitis C virus, (ii) alcohol abuse, (iii) history of chronic liver disease and (iv) pregnancy | 167 | Hepatitis |
| Wang 2010 | China | Case-control | NR | INH, RIF, PZA, and EMB for 2 months followed by INH and RIF for 4 months (2HRZE/4HR) | TB patients with (cases) and without (controls) ADIH  Exclusion criteria: patients with chronic hepatitis B virus infection, alcohol consumption, obesity, senility and poor nutritional status | 215 | ADIH |
| Wang 2015a (GI: ADACS) | China | Case-control | NR | All primary/retreatment patients with pulmonary TB were treated with a combination regimen including INH (600 mg), RIF (600 mg, or 450 mg if body weight was < 50 kg), PZA (2000 mg) and EMB (1250 mg) for the first 2 months (retreatment patients were injected with SM 750 mg each time simultaneously) and then with the same regimen, without PZA and EMB, for another 4 months for primary patients and with the same regimen, without PZA and SM, for another 6 months for retreatment patients | Inclusion criteria: newly diagnosed patients with sputum smear positive pulmonary TB  Exclusion criteria: 1) abnormal serum ALT, AST or total bilirubin levels before anti-TB treatment; 2) positive serological testing for hepatitis B virus; 3) concomitant use of hepatotoxic drugs or regular alcohol intake; 4) history of liver disease or systemic diseases that may cause liver dysfunction | 445 | ATDH |
| Wang 2015b | Taiwan | Prospective cohort | 6 months | Daily INH, RIF, EMB, and PZA in the first 2 months, and daily INH and RIF for the next 4 months  The daily dosage of each drug was calculated by weight. | Adult Taiwanese (>16 years) patients with culture-confirmed pulmonary TB  Subjects were excluded if they were pregnant, had a life expectancy <6 months, had abnormal baseline LFT, or had Mycobacterium tuberculosis isolates resistant to INH, RIF, or both | 355 in the derivation cohort, 182 in the validation cohort | HATT |
| Wang 2015c (GI: ADACS) | China | Case-control | NR | All primary/retreatment patients received standard anti-TB treatment recommended by WHO for 6–9 months. Patients were given INH (600 mg), RIF (600 or 450 mg if body weight was <50 kg), PZA (2000 mg) and EMB (1250 mg) for the first 2 months (retreatment patients were injected with SM 750 mg each time simultaneously) and then with the same regimen, without PZA and EMB, for another 4 months for primary patients and with the same regimen, without PZA and SM, for another 6 months for retreatment patients | Inclusion criteria: patients with sputum smear-positive pulmonary TB  Exclusion criteria: (i) abnormal serum ALT, AST or total bilirubin levels before treatment; (ii) positive serological testing for hepatitis B virus; (iii) concomitant use of hepatotoxic drugs or regular alcohol intake; and (iv) history of liver disease or systemic diseases that may cause liver dysfunction | 445 | ATDH |
| Xiang 2014 | China | Prospective cohort | 2 months | All patients were prescribed INH (600 mg), RIF (600 mg, or 450 mg if the body weight was less than 50 kg), PZA (2,000 mg), and EMB (1,250 mg) every other day in the first 2 months. After 2 months, INH and were continued for a further 4 to 6 months. Retreatment patients in addition received SM (750 mg) every other day in the first 2 months and continued receiving EMB for another 6 months | Inclusion criteria: newly diagnosed pulmonary TB patients belonging to the Uyghur ethnic group, who were receiving standard short-course chemotherapy recommended by WHO, who attended for a 2-month assessment, and any patients attending clinic with suspected liver disease after the start of treatment, prior to the 2-month visit  Exclusion criteria: patients who had signs of abnormal liver function when they started treatment (jaundice or elevated ALT, AST or bilirubin levels), or disease associated with liver dysfunction | 2244 | ATDILI |
| Yimer 2011 | Ethiopia | Prospective cohort | followed up for development of DILI for up to 56 weeks | All study participants received RIF based short-course chemotherapy for TB following the national TB treatment guideline. ART was then initiated. | Newly diagnosed ART and anti-TB treatment naïve adult TB and HIV co-infected patients. The eligibility criteria were age >18 years, CD4 count, 200 cells/UL, not pregnant and not on other known hepatotoxic drugs concurrently (except co-trimoxazole, 960 mg per day, which was given to all participants before enrolment and during the follow-up period according to the treatment guideline). | 353 | Anti-tubercular and antiretroviral drugs induced liver injury |
| Zazuli 2015 | Indonesia | Prospective cohort | ALT and AST serum levels were obtained 5 times: before ATD treatment, after completing the second, the fourth, and the sixth months of ATD therapy and 1 month after the patient finished the last ATD therapy | All patients received FDC-ATD category I intensive phase (RIF 150 mg, INH 75 mg, PZA 400 mg and EMB 275 mg per tablet) and FDC-ATD category I continuation phase (RIF 150 mg and INH 150 mg) in 6 months of therapy. The dosage of ATDs was selected according to the patients’ weight | Inclusion criteria: (i) adult patients (over 18 years of age) newly diagnosed with active lung TB; (ii) treated with fixed-dose combination of anti-TB drugs (FDC-ATD) category I using the DOTS strategy; (iii) written informed consent  Exclusion criteria: (i) positive HIV/AIDS; (ii) abnormal serum ALT and AST levels over twice the ULN value before treatment with ATD; (iii) hepatitis or history of hepatitis; (iv) haemoglobin serum <8 mg/dL; (v) not having ATDs for over 2 weeks; (vi) history of kidney disease and (vii) refusal to blood collection | 106 | ATDILI |

ADIH=anti-tuberculosis drug-induced hepatitis; ALP=alkaline phosphatase; ALT=alanine aminotransferase; AST=aspartate aminotransferase; ATD=anti-tuberculosis drug; ATDH=anti-tuberculosis drug-induced hepatotoxicity; ATDILI=anti-tuberculosis drug-induced liver injury; ATT=anti-tuberculosis treatment; DIH=drug-induced hepatotoxicity; DILI=drug-induced liver injury; DOTS=directly observed treatment short-course; EMB=ethambutol; FDC=fixed-dose combination; HATT=hepatotoxicity during anti-tuberculosis treatment; HIV=human immunodeficiency virus; IgM=immunoglobulin M; INH=isoniazid; LFT=liver function test; MPE=maculopapular eruption; NR=not reported; PZA=pyrazinamide; RIF=rifampicin; SM=streptomycin; TB=tuberculosis; ULN=upper limit of normal; WHO=World Health Organization
