## Additional File 4 for "Association of variants within the *GST* and other genes with anti-tubercular agents related toxicity: a systematic review and meta-analysis"

**Additional file 4** Definitions of hepatotoxicity

| Study | Outcome and definition |
| --- | --- |
| Brito 2014 | Criteria for the diagnosis of anti-TB drug-induced hepatitis was an elevation in liver function tests, AST and/or ALT of more than 3-fold the ULN (reference values: 40 and 65 U/L, respectively) and/or total bilirubin up to >2.0 mg/dL in the presence of gastrointestinal symptoms such as anorexia, nausea, vomiting and/or jaundice, with serum ALT level normalisation after anti-TB drug discontinuation. |
| Chang 2012 | ATDH was “defined according to the classification of the CIOMS (1).” No further information was provided. |
| Chatterjee 2010 | ATDH was defined as: a rise of serum (i) ALT level ≥3 x ULN or (ii) total bilirubin >1.0 mg⁄dL or, (iii) ALT <3 x ULN but associated with severe anorexia, nausea, vomiting. |
| Chen 2015,  Tang 2012, Wang 2015a, Wang 2015c (all GI: ADACS) | ATDH was defined as: (i) an increase in ALT levels greater than 2 x ULN with/without a combined increase in AST and total bilirubin levels provided one of them was greater than 2 x ULN during the treatment (1);(ii) causality assessment result was certain, probable or possible based on the WHO Uppsala Monitoring Center system (2). |
| Feng 2014, Teixeira 2011, | Anti-TB drug-induced hepatitis (Teixeira 2011)/Anti-TB drug-induced hepatic injury (Feng 2014): An increase in serum transaminase values to higher than 3 x ULN values (40 IU/L ALT in Feng) and symptoms compatible with hepatitis. |
| Gupta 2013  (GI: GUPTA), Nanashima 2012 | Increase in ALT over 2 x ULN or a combined increase in AST and bilirubin levels, provided one of them is above 2 x ULN, was defined as ATDH according to the international consensus meeting (1). Nanashima (2012) specified the ULN ranges (ALT: normal ≤42 IU/L; AST: normal ≤33 IU/L; total bilirubin: normal ≤1.5 mg/dL). |
| He 2015 | ADLI was defined according to the Danan criteria promulgated in 1990 (1, 3). No further information was provided. |
| Huang 2007 | The inclusion criteria of DILI patients were based on the suggestion of Drug-induced Liver Injury Network (4) as follows: (i) an increase in serum ALT or AST level >5 x ULN, or an elevation in ALP greater than twice the ULN, confirmed on at least two consecutive blood draws; (ii) if baseline ALT, AST or ALP are known and elevated, then ALT or AST greater than 5 x baseline value, or ALP greater than twice baseline level on at least two consecutive blood draws; (iii) any elevation of serum ALT, AST, or ALP, associated with an increased serum total bilirubin (>2.5 mg/dL), in the absence of prior diagnosis of liver disease, Gilbert’s syndrome, or evidence of haemolysis; (iv) a causality assessment score (5) greater than five, cases of which categorized as probable or highly probable DILI. |
| Kim 2009, Kim 2012b (both GI: KIM) | Anti-TB drug-induced hepatitis was defined as an elevation in the serum levels of ALT above 2 x ULN range (≤40 U/mL) during treatment and normalisation of these values after cessation of medication according to the criteria from the international consensus meeting (1). |
| Kim 2010 (GI: KIM), Kim 2012c (GI: KIM), Leiro 2008 | Anti-TB drug-induced hepatitis was defined as an elevation in the serum level of ALT or AST >3 x ULN range ( 40 U/L) during treatment, according to the American Thoracic Society guidelines (6). |
| Kim 2015 (GI: KIM) | Anti-TB drug-induced hepatitis was defined as an elevation in the serum levels of AST or ALT of more than 3 x ULN during treatment and normalisation of these values after cessation of treatment. |
| Kwon 2012 | DILI was defined as drug-induced acute hepatocellular injury with an AST level ≥ULN and an (AST/ULN)/(ALP/ULN) ratio ≥5. |
| Li 2012 | DILI was defined as serum ALT levels 3 or more x ULN, and/or serum bilirubin levels 2 or more x ULN. |
| Liu 2014 | The diagnostic criteria of ATDH was based on the international consensus (1, 5, 7): (i) serum ALT >2 x ULN (40 IU/L); or (ii) serum direct bilirubin >2 x ULN (6.8 μmol/L); or (iii) increases of serum AST (40 IU/L), ALP (220 IU/L), and total bilirubin (19.0 μmol/L); moreover, one of them >2 x ULN; or (iv) any index mentioned above >1 x ULN and associated with liver damage symptoms, such as skin or sclera yellow dye, severe anorexia, nausea, vomiting, fever, rash, itching. |
| Monteiro 2012 | Liver injury was defined as an increase in serum ALT levels beyond twice the ULN (ALT ≥42 IU/L), or at least a 2-fold increase in ALT initial levels for those patients with a baseline ALT of >84 IU/L, during the treatment period. |
| Rana 2014  (GI: RANA) | ATDH was defined according to international consensus criteria (1). Patients with a rise in serum AST or ALT levels more than or equal to 5 x ULN, irrespective of symptoms and serum bilirubin levels, or patients with rise in serum AST or ALT levels more than or equal to 2 x ULN with hyperbilirubinaemia and an absence of serological evidence of infection with hepatitis viruses (A, B, C and E) were considered as having ATDH. |
| Roy 2001 | ATDH was defined according to the international consensus criteria (1) with regard to chronology and causation for drug-induced liver diseases. However, only icteric hepatitis cases (serum bilirubin >3.0 mg/dL), among those fulfilling the above criteria, were included in the study. No further information was provided. |
| Sharma 2014 | ATDH was diagnosed if any one of criteria i), ii) or iii) were present along with criteria iv) and v). The criteria were: i) an increase of 5 x ULN (50 IU/L) of serum AST and/or ALT levels on one occasion or more than 3 times (>150 IU/L) on 3 consecutive occasions; ii) serum total bilirubin level >1.5 mg/dL; iii) any increase in serum AST and/or ALT above pre-treatment values, together with anorexia, nausea, vomiting and jaundice; iv) absence of serological evidence of infection with hepatitis viruses A, B, C or E; and v) improvement in liver function (serum bilirubin <1 mg/dL, AST and ALT <100 IU/l) after the withdrawal of anti-TB drugs. |
| Singla 2014 | International consensus criteria (1) define ATDH as development of more than 2 x ULN value of ALT and AST. The ULN values used in this study were 35 U/L ALT and 40 U/L AST. |
| Sotsuka 2011 | The severity of hepatotoxicity (hepatotoxicity A-D) was judged by the increase in either AST or ALT levels from the ULN range (AST, 33 U/L; ALT, 42 U/L): hepatotoxicity A, above the upper limit and less than 2-fold increase; hepatotoxicity B, 2- to 3-fold increase; hepatotoxicity C, 3- to 4-fold increase; hepatotoxicity D, greater than 4-fold increase. Results for grades B–D of hepatotoxicity were used in this review as clinical opinion was that the hepatotoxicity A patients would not have met the criteria for hepatotoxicity in many of the other studies included in this review. |
| Wang 2010 | The selection criteria for ATDH were as follows: (i) ALT ≥2 x ULN; (ii) increased AST/ALT/serum proteins (i.e. liver damage based on an increase in ALT or bilirubin ≥2 x ULN, or an increase in AST, ASP and total bilirubin with at least one of these being ≥2 x ULN); (iii) negative for hepatitis A antibody, hepatitis B surface antigen and hepatitis C marker; (iv) no other factors influencing the levels of AST/ALT/serum proteins, such as alcohol-induced liver disease, hypoxia, auto-immune disease, congestive heart failure and bacteraemia; and (v) causality assessment score >5. |
| Wang 2015b | Hepatitis during anti-TB treatment was defined as increased serum AST and/or ALT >3 x ULN in symptomatic patients, or >5 x ULN in asymptomatic patients. The diagnosis of INH- or RMP-induced hepatitis required a positive re-challenge test (at least doubling of serum AST or ALT levels and recurrence of clinical symptoms of hepatitis after re-challenge), whereas PZA-induced hepatitis was diagnosed either by a positive re-challenge test or by exclusion. Results are presented for overall drug-induced HATT and INH-induced HATT separately. In this review, we used the results for overall drug-induced HATT as our review focusses on hepatotoxicity induced by any anti-tuberculosis drug. |
| Xiang 2014 | ADLI was defined as an ALT, AST or bilirubin value more than 2 x ULN value. The ULN used in the study was 40 U/L for ALT, 40 U/L for AST and 19 μmol/L for total bilirubin. |
| Yimer 2011 | DILI in patients was defined according to the international consensus criteria (1). Liver biochemical parameters more than 2 x ULN value were considered as hepatotoxicity. |
| Zazuli 2015 | Hepatotoxicity was defined as ALT and/or AST levels above the normal threshold on the second, fourth and sixth months of monitoring during tuberculosis treatment. |

ADLI: anti-tuberculosis drug-induced liver injury; ALP: alkaline phosphatase; ALT: alanine aminotransferase; AST: aspartate aminotransferase; ATDH: anti-tuberculosis drug-induced hepatotoxicity; CIOMS: Council for International Organizations of Medical Sciences; DILI: drug-induced liver injury; GI: group identifier; HATT: hepatitis during anti-tuberculosis treatment; INH: isoniazid; PZA: pyrazinamide; RMP: rifampicin; TB: tuberculosis; ULN: upper limit of normal; WHO: World Health Organization
