## Additional File 5 for "Association of variants within the *GST* and other genes with anti-tubercular agents related toxicity: a systematic review and meta-analysis"

| Outcome | Study | Outcome definition |
| --- | --- | --- |
| ADRs | Costa (2012) | The presence of at least one of the following symptoms during the follow-up period: gastric, joint, neuromuscular, or skin reactions; and hepatotoxicity (in accordance with the criteria of drug-induced liver injuries developed by the international consensus meeting) (1). |
| ATD-induced MPE | Kim 2012a (GI: KIM) | The development of MPE after receiving first-line ATD and the disappearance of MPE after discontinuing ATD. |
| ATD-induced cutaneous reactions | Kim 2010 (GI: KIM) | The development of any cutaneous symptom or skin lesion after receiving ATD medication. |

**Additional file 5** Definitions of other toxicity outcomes

ADR: adverse drug reaction; ATD: anti-tuberculosis drug; GI: group identifier; MPE: macropapular eruption

1.	Saukkonen JJ, Cohn DL, Jasmer RM, Schenker S, Jereb JA, Nolan CM, et al. An official ATS statement: hepatotoxicity of antituberculosis therapy. Am J Respir Crit Care Med. 2006;174(8):935-52.
