## Additional File 6 for "Association of variants within the *GST* and other genes with anti-tubercular agents related toxicity: a systematic review and meta-analysis"

**Additional file 6** Results of the sensitivity analyses

**Sensitivity analysis 1** Pairwise comparisons for the *GSTM1* null polymorphism

As only one study reported on each genotype group separately for the *GSTM1* gene, no meta-analysis was performed. Instead, we calculated odds ratios and corresponding 95% confidence intervals, as shown in Table 1 below.

**Table 1** *GSTM1* null polymorphism and anti-tuberculosis drug-induced hepatotoxity: results of pairwise comparisons

| Study | Country | Ethnicity | Comparison | OR (95% CI) | # cases | # controls | I^2^ |
| --- | --- | --- | --- | --- | --- | --- | --- |
| Liu (2014) | China | NR | Het vs Hom present | 0.42 (0.02, 8.18) | 6 | 47 | N/A |
|  |  |  | Hom null vs Hom present | 0.97 (0.35, 2.71) | 20 | 136 | N/A |

CI: confidence interval; Het: heterozygous genotype; Hom: homozygous; NR: not reported; OR: odds ratio

**Sensitivity analysis 2** Pairwise comparisons for the *GSTT1* null polymorphism

*Heterozygous genotype versus homozygous present genotype*

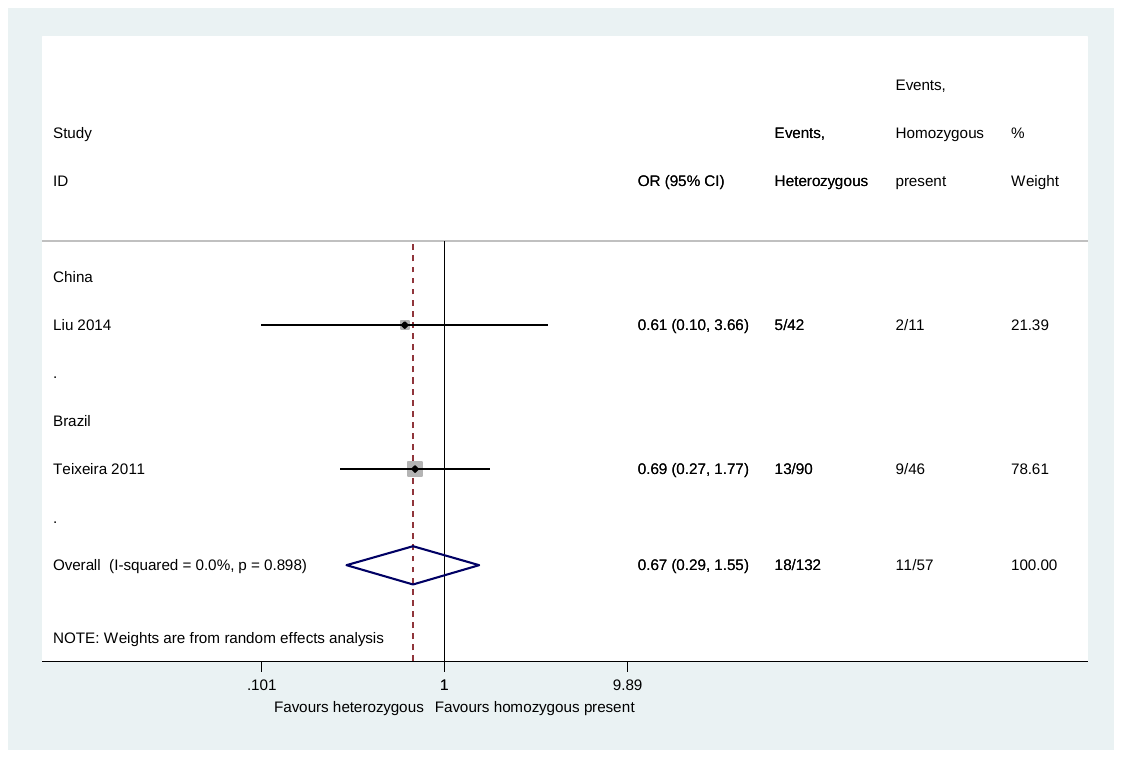

**Fig. 1** *GSTT1* null polymorphism and anti-tuberculosis drug-induced hepatotoxicity: heterozygous genotype versus homozygous present genotype

CI: confidence interval; OR: odds ratio

*Homozygous null genotype versus homozygous present genotype*

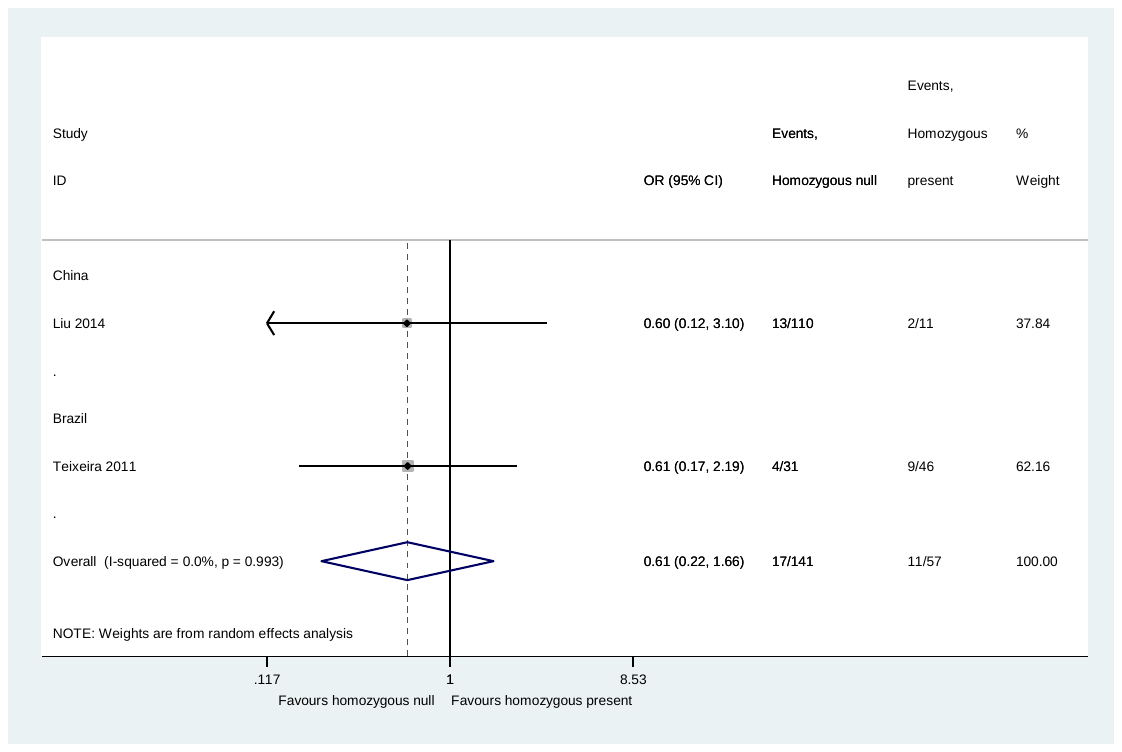

**Fig. 2** GSTT1 null polymorphism and anti-tuberculosis drug-induced hepatotoxicity: homozygous null genotype versus homozygous present genotype

CI: confidence interval; OR: odds ratio
