## Additional File 7 for "Association of variants within the *GST* and other genes with anti-tubercular agents related toxicity: a systematic review and meta-analysis"

**Additional file 7** Funnel plots for the primary analyses

GSTM1 *null polymorphism*

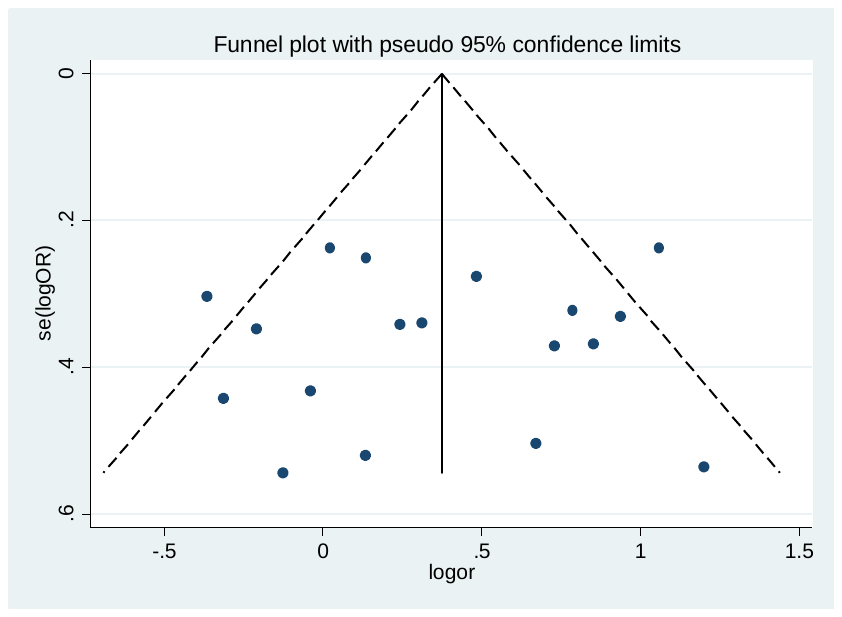

**Fig. 1** Funnel plot for the analysis of GSTM1 null polymorphism and anti-tuberculosis drug-induced hepatotoxicity

GSTT1 *null polymorphism*

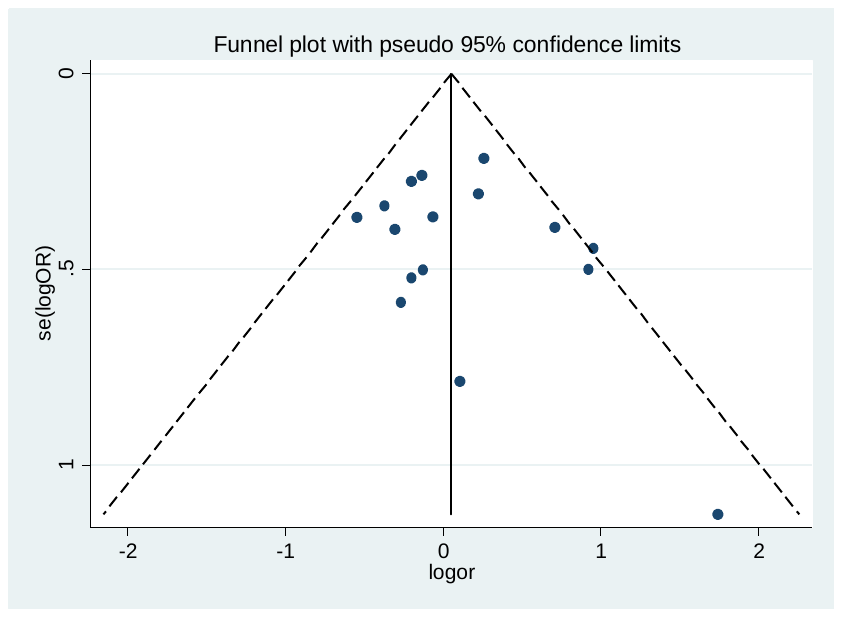

**Fig. 2** Funnel plot for the analysis of GSTT1 null polymorphism and anti-tuberculosis drug-induced hepatotoxicity
