## Additional File 8 for "Association of variants within the *GST* and other genes with anti-tubercular agents related toxicity: a systematic review and meta-analysis"

**Additional file 8** Other genetic variants and hepatotoxicity

| Gene | Variant | Comparison | Country (no. of studies) | Ethnicity | OR (95% CI) | # cases | # controls | I^2^ value |
| --- | --- | --- | --- | --- | --- | --- | --- | --- |
| *ABCB1* | Rs1045642^a^ | Het (CT) vs Hom WT (CC) | South Korea (1 study) | NR | 1.13 (0.61, 2.09) | 58 | 137 | N/A |
|  |  |  | Ethiopia (1 study) | NR | 0.43 (0.18, 1.06) | 36 | 154 | N/A |
|  |  |  | All (2 studies) | | 0.74 (0.29, 1.89) | 94 | 291 | 67.0% |
|  |  | Hom MT (TT) vs Hom WT (CC) | South Korea (1 study) | NR | 1.01 (0.40, 2.56) | 35 | 88 | N/A |
|  |  |  | Ethiopia (1 study) | NR | 2.84 (0.81, 10.00) | 34 | 105 | N/A |
|  |  |  | All (2 studies) | | 1.55 (0.57, 4.23) | 69 | 193 | 41.0% |
|  | Rs10261685 | Het (TG) vs Hom WT (TT) | South Korea (1 study) | NR | 0.72 (0.32, 1.62) | 65 | 159 | N/A |
|  |  | Hom MT (GG) vs Hom WT (TT) | South Korea (1 study) | NR | 6.93 (0.28, 172.71) | 57 | 130 | N/A |
| *ABCC2* | 1774 G-del^b^ | Het (G/-) vs Hom WT (GG) | South Korea (1 study) | NR | 0.60 (0.32, 1.14) | 58 | 137 | N/A |
|  |  | Hom MT (-/-) vs Hom WT (GG) | South Korea (1 study) | NR | 0.69 (0.26, 1.79) | 33 | 63 | N/A |
|  | Rs1885301 | Het (GA) vs Hom WT (GG) | South Korea (1 study) | NR | 1.18 (0.65, 2.15) | 61 | 149 | N/A |
|  |  | Hom MT (AA) vs Hom WT (GG) | South Korea (1 study) | NR | 1.87 (0.56, 6.29) | 39 | 96 | N/A |
|  | Rs717620 | Het (CT) vs Hom WT (CC) | South Korea (1 study) | NR | 1.10 (0.61, 2.01) | 64 | 149 | N/A |
|  |  | Hom MT (TT) vs Hom WT (CC) | South Korea (1 study) | NR | 0.81 (0.16, 4.18) | 40 | 98 | N/A |
|  | Rs2804400 | Het (CT) vs Hom WT (CC) | South Korea (1 study) | NR | 1.27 (0.70, 2.30) | 63 | 152 | N/A |
|  |  | Hom MT (TT) vs Hom WT (CC) | South Korea (1 study) | NR | 1.53 (0.42, 5.56) | 38 | 98 | N/A |
|  | Rs2273697 | Het (GA) vs Hom WT (GG) | South Korea (1 study) | NR | 0.48 (0.17, 1.32) | 66 | 157 | N/A |
|  |  | Hom MT (AA) vs Hom WT (GG) | South Korea (1 study) | NR | 0.44 (0.02, 9.25) | 61 | 136 | N/A |
|  | Rs3740070 | Het (GA) vs Hom WT (GG) | South Korea (1 study) | NR | 1.00 (0.34, 2.97) | 66 | 159 | N/A |
|  |  | Hom MT (AA) vs Hom WT (GG) | South Korea (1 study) | NR | Data excluded^c^ | 61 | 147 | N/A |
|  | Rs3740066 | Het (CT) vs Hom WT (CC) | South Korea (1 study) | NR | 1.21 (0.66, 2.21) | 61 | 147 | N/A |
|  |  | Hom MT (TT) vs Hom WT (CC) | South Korea (1 study) | NR | 1.44 (0.45, 4.61) | 40 | 100 | N/A |
| *BACH1* | rs2300301 | Het (AG) vs Hom WT (AA) | Japan (1 study) | NR | 0.40 (0.13, 1.29) | 14 | 73 | N/A |
|  |  | Hom MT (GG) vs Hom WT (AA) | Japan (1 study) | NR | 1.33 (0.31, 5.72) | 11 | 30 | N/A |
|  | rs1153285 | Het (GA) vs Hom WT (GG) | Japan (1 study) | NR | 1.21 (0.39, 3.75) | 15 | 76 | N/A |
|  |  | Hom MT (AA) vs Hom WT (GG) | Japan (1 study) | NR | 2.83 (0.55, 14.54) | 9 | 40 | N/A |
|  | rs2070401 | Het (TC) vs Hom WT (TT) | Japan (1 study) | NR | 1.13 (0.37, 3.50) | 15 | 81 | N/A |
|  |  | Hom MT (CC) vs Hom WT (TT) | Japan (1 study) | NR | 17.00 (1.59, 182.14) | 12 | 52 | N/A |
| *GSTP1* | Ile105Val | Hom WT vs Het or Hom MT | China (1 study) | NR | 1.84 (1.10, 3.07) | 127 | 127 | N/A |
| *HMOX1* | rs2071746 | Het (TA) vs Hom WT (TT) | Japan (1 study) | NR | 2.67 (0.69, 10.31) | 15 | 70 | N/A |
|  |  | Hom MT (AA) vs Hom WT (TT) | Japan (1 study) | NR | 2.33 (0.41, 13.26) | 6 | 40 | N/A |
|  | rs2071749 | Het (GA) vs Hom WT (GG) | Japan (1 study) | NR | 2.44 (0.77, 7.67) | 16 | 78 | N/A |
|  |  | Hom MT (AA) vs Hom WT (GG) | Japan (1 study) | NR | 4.10 (0.59, 28.38) | 7 | 45 | N/A |
|  | rs5755720 | Het (AG) vs Hom WT (AA) | Japan (1 study) | NR | 0.81 (0.26, 2.55) | 16 | 61 | N/A |
|  |  | Hom MT (GG) vs Hom WT (AA) | Japan (1 study) | NR | 0.32 (0.06, 1.76) | 8 | 41 | N/A |
| *HSPA1L* | rs2227956 | Het (TC) vs Hom WT (TT) | China (1 study) | 100% Chinese | 1.98 (1.21, 3.23) | 82 | 321 | N/A |
|  |  | Hom MT (CC) vs Hom WT (TT) | China (1 study) | 100% Chinese | 1.16 (0.45, 3.00) | 42 | 223 | N/A |
| *IL4* | rs2243289 | Het (GA) vs Hom WT (GG) | China (1 study) | 100% Chinese | 0.84 (0.50, 1.41) | 85 | 341 | N/A |
|  |  | Hom MT (AA) vs Hom WT (GG) | China (1 study) | 100% Chinese | 1.43 (0.37, 5.54) | 64 | 240 | N/A |
|  | rs2243250 | Het (TC) vs Hom WT (TT) | China (1 study) | 100% Chinese | 0.82 (0.49, 1.38) | 86 | 346 | N/A |
|  |  | Hom MT (CC) vs Hom WT (TT) | China (1 study) | 100% Chinese | 1.62 (0.41, 6.46) | 65 | 242 | N/A |
|  | rs2070874 | Het (TC) vs Hom WT (TT) | China (1 study) | 100% Chinese | 0.80 (0.47, 1.35) | 85 | 343 | N/A |
|  |  | Hom MT (CC) vs Hom WT (TT) | China (1 study) | 100% Chinese | 1.62 (0.41, 6.44) | 65 | 241 | N/A |
| *IL6* | rs2066992 | Het (GT) vs Hom WT (GG) | China (1 study) | 100% Chinese | 1.15 (0.69, 1.92) | 79 | 335 | N/A |
|  |  | Hom MT (TT) vs Hom WT (GG) | China (1 study) | 100% Chinese | 2.35 (1.03, 5.36) | 60 | 242 | N/A |
|  | rs2069837 | Het (AG) vs Hom WT (AA) | China (1 study) | 100% Chinese | 1.41 (0.85, 2.33) | 87 | 335 | N/A |
|  |  | Hom MT (GG) vs Hom WT (AA) | China (1 study) | 100% Chinese | 0.57 (0.13, 2.57) | 59 | 259 | N/A |
|  | rs1524107 | Het (CT) vs Hom WT (CC) | China (1 study) | 100% Chinese | 1.18 (0.71, 1.96) | 80 | 333 | N/A |
|  |  | Hom MT (TT) vs Hom WT (CC) | China (1 study) | 100% Chinese | 1.99 (0.85, 4.63) | 59 | 241 | N/A |
| *IL10* | rs1800896 | Het (AG) vs Hom WT (AA) | China (1 study) | 100% Chinese | 0.90 (0.43, 1.87) | 88 | 353 | N/A |
|  |  | Hom MT (GG) vs Hom WT (AA) | China (1 study) | 100% Chinese | 3.96 (0.25, 64.0) | 79 | 310 | N/A |
|  | rs1800871 | Het (TC) vs Hom WT (TT) | China (1 study) | 100% Chinese | 1.11 (0.68, 1.82) | 79 | 316 | N/A |
|  |  | Hom MT (CC) vs Hom WT (TT) | China (1 study) | 100% Chinese | 1.13 (0.52, 2.46) | 52 | 213 | N/A |
|  | rs1800872 | Het (AC) vs Hom WT (AA) | China (1 study) | 100% Chinese | 1.12 (0.68, 1.84) | 78 | 314 | N/A |
|  |  | Hom MT (CC) vs Hom WT (AA) | China (1 study) | 100% Chinese | 1.18 (0.54, 2.57) | 51 | 210 | N/A |
| *KEAP1* | rs1048290 | Het (GC) vs Hom WT (GG) | Japan (1 study) | NR | 1.12 (0.30, 4.13) | 12 | 64 | N/A |
|  |  | Hom MT (CC) vs Hom WT (GG) | Japan (1 study) | NR | 1.92 (0.47, 7.83) | 10 | 41 | N/A |
|  | rs11545829 | Het (CT) vs Hom WT (CC) | Japan (1 study) | NR | 0.97 (0.31, 3.06) | 14 | 71 | N/A |
|  |  | Hom MT (TT) vs Hom WT (CC) | Japan (1 study) | NR | 1.82 (0.45, 7.39) | 11 | 46 | N/A |
| *MAFF* | rs2413508 | Het (CG) vs Hom WT (CC) | Japan (1 study) | NR | 1.04 (0.32, 3.41) | 13 | 72 | N/A |
|  |  | Hom MT (GG) vs Hom WT (CC) | Japan (1 study) | NR | 2.83 (0.71, 11.27) | 11 | 44 | N/A |
|  | rs2267373 | Het (TC) vs Hom WT (TT) | Japan (1 study) | NR | 1.86 (0.55, 6.27) | 13 | 67 | N/A |
|  |  | Hom MT (CC) vs Hom WT (TT) | Japan (1 study) | NR | 2.40 (0.60, 9.52) | 10 | 51 | N/A |
|  | rs2235264 | Het (AG) vs Hom WT (AA) | Japan (1 study) | NR | 1.49 (0.36, 6.14) | 11 | 67 | N/A |
|  |  | Hom MT (GG) vs Hom WT (AA) | Japan (1 study) | NR | 3.73 (0.83, 16.71) | 10 | 39 | N/A |
|  | rs4821765 | Het (TC) vs Hom WT (TT) | Japan (1 study) | NR | 0.56 (0.17, 1.88) | 17 | 82 | N/A |
|  |  | Hom MT (CC) vs Hom WT (TT) | Japan (1 study) | NR | 11.89 (0.46, 308.42) | 14 | 53 | N/A |
| *MAFK* | rs4720833 | Het (GA) vs Hom WT (GG) | Japan (1 study) | NR | 4.03 (1.30, 12.52) | 18 | 74 | N/A |
|  |  | Hom MT (AA) vs Hom WT (GG) | Japan (1 study) | NR | 0.49 (0.02, 9.64) | 5 | 53 | N/A |
|  | rs3808337 | Het (TC) vs Hom WT (TT) | Japan (1 study) | NR | 2.28 (0.76, 6.81) | 17 | 74 | N/A |
|  |  | Hom MT (CC) vs Hom WT (TT) | Japan (1 study) | NR | 0.85 (0.09, 8.09) | 7 | 49 | N/A |
| *NFE2L2* | rs2886161 | Het (CT) vs Hom WT (CC) | Japan (1 study) | NR | 1.25 (0.41, 3.75) | 16 | 65 | N/A |
|  |  | Hom MT (TT) vs Hom WT (CC) | Japan (1 study) | NR | 0.54 (0.10, 2.88) | 9 | 49 | N/A |
|  | rs4243387 | Het (TC) vs Hom WT (TT) | Japan (1 study) | NR | 0.76 (0.24, 2.37) | 17 | 76 | N/A |
|  |  | Hom MT (CC) vs Hom WT (TT) | Japan (1 study) | NR | 0.68 (0.07, 6.20) | 13 | 55 | N/A |
|  | rs6726395 | Het (GA) vs Hom WT (GG) | Japan (1 study) | NR | 0.41 (0.12, 1.40) | 15 | 72 | N/A |
|  |  | Hom MT (AA) vs Hom WT (GG) | Japan (1 study) | NR | 1.04 (0.24, 4.44) | 14 | 48 | N/A |
|  | rs2001350 | Het (AG) vs Hom WT (AA) | Japan (1 study) | NR | 0.64 (0.19, 2.19) | 16 | 76 | N/A |
|  |  | Hom MT (GG) vs Hom WT (AA) | Japan (1 study) | NR | 1.39 (0.25, 7.76) | 14 | 56 | N/A |
| *NOS2A* | rs10459953 | Het (CG) vs Hom WT (CC) | Japan (1 study) | NR | 0.55 (0.17, 1.79) | 14 | 59 | N/A |
|  |  | Hom MT (GG) vs Hom WT (CC) | Japan (1 study) | NR | 0.52 (0.13, 2.04) | 11 | 44 | N/A |
|  | rs3794764 | Het (GA) vs Hom WT (GG) | Japan (1 study) | NR | 1.05 (0.36, 3.04) | 17 | 80 | N/A |
|  |  | Hom MT (AA) vs Hom WT (GG) | Japan (1 study) | NR | 2.40 (0.20, 29.10) | 11 | 50 | N/A |
|  | rs12944039 | Het (GA) vs Hom WT (GG) | Japan (1 study) | NR | 1.22 (0.42, 3.58) | 16 | 80 | N/A |
|  |  | Hom MT (AA) vs Hom WT (GG) | Japan (1 study) | NR | 5.50 (0.67, 44.90) | 10 | 46 | N/A |
|  | rs11080344 | Het (CT) vs Hom WT (CC) | Japan (1 study) | NR | 0.40 (0.13, 1.19) | 17 | 69 | N/A |
|  |  | Hom MT (TT) vs Hom WT (CC) | Japan (1 study) | NR | 0.20 (0.02, 1.74) | 12 | 42 | N/A |
|  | rs2314810 | Het (GC) vs Hom WT (GG) | Japan (1 study) | NR | 0.91 (0.32, 2.58) | 18 | 80 | N/A |
|  |  | Hom MT (CC) vs Hom WT (GG) | Japan (1 study) | NR | 0.83 (0.04, 18.41) | 11 | 49 | N/A |
|  | rs3729966 | Het (CT) vs Hom WT (CC) | Japan (1 study) | NR | 0.73 (0.25, 2.14) | 16 | 76 | N/A |
|  |  | Hom MT (TT) vs Hom WT (CC) | Japan (1 study) | NR | 1.33 (0.23, 7.89) | 10 | 38 | N/A |
|  | rs944722 | Het (TC) vs Hom WT (TT) | Japan (1 study) | NR | 0.84 (0.29, 2.42) | 17 | 79 | N/A |
|  |  | Hom MT (CC) vs Hom WT (TT) | Japan (1 study) | NR | 1.43 (0.13, 15.26) | 11 | 46 | N/A |
|  | rs2255929 | Het (AT) vs Hom WT (AA) | Japan (1 study) | NR | 0.68 (0.23, 2.02) | 16 | 73 | N/A |
|  |  | Hom MT (TT) vs Hom WT (AA) | Japan (1 study) | NR | 0.84 (0.15, 4.59) | 11 | 43 | N/A |
|  | rs3794756 | Het (CT) vs Hom WT (CC) | Japan (1 study) | NR | 0.31 (0.08, 1.20) | 15 | 79 | N/A |
|  |  | Hom MT (TT) vs Hom WT (CC) | Japan (1 study) | NR | 3.67 (0.65, 20.54) | 15 | 47 | N/A |
| *NQO1* | 609C-T  (rs1800566) | Het (CT) vs Hom WT (CC) | Taiwan (1 study) | NR | 0.93 (0.40, 2.16) | 53 | 49 | N/A |
|  |  |  | Japan (1 study) | NR | 1.45 (0.47, 4.41) | 16 | 71 | N/A |
|  |  |  | All (2 studies) | | 1.09 (0.56, 2.14) | 69 | 120 | 0.0% |
|  |  | Hom MT (TT) vs Hom WT (CC) | Taiwan (1 study) | NR | 0.63 (0.22, 1.83) | 27 | 29 | N/A |
|  |  |  | Japan (1 study) | NR | 1.00 (0.18, 5.70) | 8 | 44 | N/A |
|  |  |  | All (2 studies) | | 0.72 (0.29, 1.78) | 35 | 73 | 0.0% |
|  | rs689452 | Het (CG) vs Hom WT (CC) | Japan (1 study) | NR | 1.29 (0.44, 3.82) | 16 | 71 | N/A |
|  |  | Hom MT (GG) vs Hom WT (CC) | Japan (1 study) | NR | 0.91 (0.17, 4.91) | 10 | 51 | N/A |
|  | rs2917669 | Het (CT) vs Hom WT (CC) | Japan (1 study) | NR | 1.37 (0.46, 4.05) | 16 | 71 | N/A |
|  |  | Hom MT (TT) vs Hom WT (CC) | Japan (1 study) | NR | 0.93 (0.17, 5.03) | 10 | 52 | N/A |
|  | rs10517 | Het (CT) vs Hom WT (CC) | Japan (1 study) | NR | 1.71 (0.56, 5.22) | 16 | 71 | N/A |
|  |  | Hom MT (TT) vs Hom WT (CC) | Japan (1 study) | NR | 1.09 (0.19, 6.20) | 8 | 47 | N/A |
| *PXR* | Rs3814055 | Het (CT) vs Hom WT (CC) | Taiwan (2 studies)^a^ | NR | 1.19 (0.75, 1.88) | 99 | 414 | 0.0% |
|  |  |  | Indonesia (1 study) | NR | 1.07 (0.43, 2.66) | 30 | 69 | N/A |
|  |  |  | All (3 studies) | | 1.16 (0.77, 1.75) | 129 | 483 | 0.0% |
|  |  | Hom MT (TT) vs Hom WT (CC) | Taiwan (2 studies)^d^ | NR | 1.19 (0.43, 3.32) | 68 | 298 | 0.0% |
|  |  |  | Indonesia (1 study) | NR | 5.88 (1.05, 32.85) | 25 | 49 | N/A |
|  |  |  | All (3 studies) | | 1.86 (0.65, 5.33) | 93 | 347 | 23.4% |
|  | Rs12488820 | Het (CT) vs Hom WT (CC) | Taiwan (2 studies)^d^ | NR | Data excluded^c^ | | | |
|  |  | Hom MT (TT) vs Hom WT (CC) | Taiwan (2 studies)^d^ | NR | 0.93 (0.31, 2.81) | 103 | 432 | 0.0% |
|  | Rs2461823 | Het (GA) vs Hom WT (GG) | Taiwan (2 studies)^d^ | NR | 0.84 (0.52, 1.36) | 83 | 377 | 0.0% |
|  |  | Hom MT (AA) vs Hom WT (GG) | Taiwan (2 studies)^d^ | NR | 1.60 (0.86, 2.98) | 59 | 211 | 0.0% |
|  | Rs7643645 | Het (AG) vs Hom WT (AA) | Taiwan (2 studies)^d^ | NR | 1.29 (0.59, 2.80) | 74 | 334 | 48.6% |
|  |  | Hom MT (GG) vs Hom WT (AA) | Taiwan (2 studies)^d^ | NR | 1.64 (0.89, 3.04) | 52 | 226 | 0.0% |
|  | Rs6785049 | Het (GA) vs Hom WT (GG) | Taiwan (2 studies)^d^ | NR | 1.12 (0.70, 1.80) | 94 | 360 | 0.0% |
|  |  | Hom MT (AA) vs Hom WT (GG) | Taiwan (2 studies)^d^ | NR | 0.56 (0.26, 1.19) | 44 | 213 | 0.0% |
|  | Rs3814057 | Het (AC) vs Hom WT (AA) | Taiwan (2 studies)^d^ | NR | 1.98 (1.06, 3.69) | 78 | 343 | 0.0% |
|  |  | Hom MT (CC) vs Hom WT (AA) | Taiwan (2 studies)^d^ | NR | 2.18 (1.07, 4.44) | 40 | 193 | 0.0% |
| *SLC10A1* | rs4646285 | Het (GA) vs Hom WT (GG) | China (1 study) | NR | 1.05 (0.61, 1.80) | 86 | 348 | N/A |
|  |  | Hom MT (AA) vs Hom WT (GG) | China (1 study) | NR | 2.05 (0.50, 8.41) | 67 | 268 | N/A |
| *SLCO1B1* | Rs4149013 | Het (AG) vs Hom WT (AA) | China (1 study) | NR | 1.61 (0.95, 2.73) | 88 | 349 | N/A |
|  |  |  | South Korea (1 study) | NR | 1.76 (0.93, 3.31) | 66 | 150 | N/A |
|  |  |  | All (2 studies) | | 1.67 (1.12, 2.50) | 154 | 499 | 0.0% |
|  |  | Hom MT (GG) vs Hom WT (AA) | China (1 study) | NR | 0.89 (0.10, 7.78) | 63 | 282 | N/A |
|  |  |  | South Korea (1 study) | NR | 0.24 (0.01, 4.46) | 43 | 120 | N/A |
|  |  |  | All (2 studies) | | 0.56 (0.10, 3.19) | 106 | 402 | 0.0% |
|  | Rs4149014^e^ | Het (TG) vs Hom WT (TT) | China (1 study) | NR | 0.86 (0.54, 1.39) | 87 | 309 | N/A |
|  |  |  | South Korea (1 study) | NR | 0.78 (0.43, 1.45) | 61 | 147 | N/A |
|  |  |  | All (2 studies) | | 0.83 (0.57, 1.21) | 148 | 456 | 0.0% |
|  |  | Hom MT (GG) vs Hom WT (TT) | China (1 study) | NR | 0.15 (0.03, 0.63) | 48 | 197 | N/A |
|  |  |  | South Korea (1 study) | NR | 1.21 (0.38, 3.87) | 43 | 92 | N/A |
|  |  |  | All (2 studies) | | 0.44 (0.05, 3.82) | 91 | 289 | 81.5% |
|  | Rs2306283^f^ | Het (GA) vs Hom WT (GG) | China (2 studies) | NR | 1.10 (0.76, 1.57) | 188 | 471 | 0.0% |
|  |  |  | South Korea (1 study) | NR | 1.12 (0.61, 2.06) | 59 | 145 | N/A |
|  |  |  | Ethiopia (1 study) | NR | 0.69 (0.32, 1.50) | 32 | 140 | N/A |
|  |  |  | All (4 studies) | | 1.03 (0.77, 1.38) | 279 | 756 | 0.0% |
|  |  | Hom MT (AA) vs Hom WT (GG) | China (2 studies) | NR | 1.31 (0.71, 2.41) | 135 | 330 | 0.0% |
|  |  |  | South Korea (1 study) | NR | 1.40 (0.48, 4.11) | 39 | 96 | N/A |
|  |  |  | Ethiopia (1 study) | NR | 1.59 (0.60, 4.21) | 24 | 73 | N/A |
|  |  |  | All (4 studies) | | 1.38 (0.87, 2.21) | 198 | 499 | 0.0% |
|  | Rs4149056 | Het (TC) vs Hom WT (TT) | China (2 studies) | NR | 1.58 (0.42, 5.89) | 204 | 505 | 88.8% |
|  |  |  | South Korea (1 study) | NR | 1.23 (0.65, 2.32) | 66 | 153 | N/A |
|  |  |  | Ethiopia (1 study) | NR | 1.05 (0.50, 2.21) | 40 | 156 | N/A |
|  |  |  | All (4 studies) | | 1.35 (0.74, 2.43) | 310 | 814 | 69.4% |
|  |  | Hom MT (CC) vs Hom WT (TT) | China (2 studies) | NR | 3.98 (0.64, 24.66) | 158 | 417 | 0.0% |
|  |  |  | South Korea (1 study) | NR | 0.35 (0.02, 6.88) | 46 | 116 | N/A |
|  |  |  | Ethiopia (1 study) | NR | 0.99 (0.11, 9.23) | 28 | 111 | N/A |
|  |  |  | All (4 studies) | | 1.62 (0.45, 5.79) | 232 | 644 | 0.0% |
|  | Rs2291075 | Het (CT) vs Hom WT (CC) | China (1 study) | NR | 0.95 (0.54, 1.65) | 68 | 262 | N/A |
|  |  | Hom MT (TT) vs Hom WT (CC) | China (1 study) | NR | 0.81 (0.42, 1.56) | 45 | 185 | N/A |
| *SOD1* | rs2070424 | Hom MT (GG) or Het (GA) vs Hom WT (AA) | South Korea (1 study) | 100% Korean | 2.28 (1.16, 4.48) | 84 | 236 | N/A |
| *SOD2 (MnSOD)* | 47 T-C (rs4880) | Hom MT (CC) or Het (CT) vs Hom WT (TT) | Taiwan (1 study) | NR | 2.40 (1.12, 5.16) | 63 | 63 | N/A |
|  |  |  | South Korea (1 study) | 100% Korean | 1.27 (0.71, 2.27) | 83 | 237 | N/A |
|  |  |  | All (2 studies) | | 1.66 (0.89, 3.08) | 146 | 300 | 40.9% |
|  |  | Het (TC) vs Hom WT (TT) | Taiwan (1 study) | NR | 2.56 (1.12, 5.81) | 59 | 60 | N/A |
|  |  | Hom MT (CC) vs Hom WT (TT) | Taiwan (1 study) | NR | 1.78 (0.37, 8.44) | 40 | 51 | N/A |
| *SOD3* | rs1799895 | Hom MT (AA) or Het (GA) vs Hom WT (GG) | South Korea (1 study) | 100% Korean | 1.03 (0.62, 1.71) | 83 | 234 | N/A |
|  | rs2536512 | Hom MT (GG) or Het (CG) vs Hom WT (CC) | South Korea (1 study) | 100% Korean | 1.81 (0.72, 4.54) | 84 | 237 | N/A |
| *STAT3* | rs1053004 | Het (TC) vs Hom WT (TT) | China (1 study) | 100% Chinese | 0.90 (0.54, 1.48) | 78 | 300 | N/A |
|  |  | Hom MT (CC) vs Hom WT (TT) | China (1 study) | 100% Chinese | 0.75 (0.36, 1.58) | 49 | 191 | N/A |
|  | rs1053023 | Het (AG) vs Hom WT (AA) | China (1 study) | 100% Chinese | 1.26 (0.72, 2.19) | 64 | 293 | N/A |
|  |  | Hom MT (GG) vs Hom WT (AA) | China (1 study) | 100% Chinese | 2.15 (1.14, 4.07) | 49 | 187 | N/A |
|  | rs1053005 | Het (AG) vs Hom WT (AA) | China (1 study) | 100% Chinese | 1.00 (0.61, 1.63) | 80 | 310 | N/A |
|  |  | Hom MT (GG) vs Hom WT (AA) | China (1 study) | 100% Chinese | 0.79 (0.36, 1.76) | 49 | 199 | N/A |
| *TNF-alpha* | -308 G-A | Het (GA) vs Hom WT (GG) | South Korea (1 study) | 100% Korean | 1.80 (0.95, 3.43) | 75 | 228 | N/A |
|  |  | Hom MT (AA) vs Hom WT (GG) | South Korea (1 study) | 100% Korean | 6.81 (0.61, 76.44) | 59 | 195 | N/A |
| *UGT1A1* | 211 G-A  (rs4148323) | Het (GA) vs Hom WT (GG) | Taiwan (1 study) | NR | 0.99 (0.31, 3.12) | 17 | 81 | N/A |
|  |  |  | South Korea (1 study) | NR | 1.03 (0.55, 1.93) | 61 | 145 | N/A |
|  |  |  | All (2 studies) | | 1.02 (0.59, 1.77) | 78 | 226 | 0.0% |
|  |  | Hom MT (AA) vs Hom WT (GG) | Taiwan (1 study) | NR | Data excluded^c^ | | | |
|  |  |  | South Korea (1 study) | NR | 1.20 (0.39, 3.73) | 45 | 106 | N/A |
|  | rs3755319 | Het (CA) vs Hom WT (AA) | South Korea (1 study) | NR | 1.55 (0.86, 2.80) | 63 | 150 | N/A |
|  |  | Hom MT (CC) vs Hom WT (AA) | South Korea (1 study) | NR | 0.83 (0.16, 4.21) | 33 | 97 | N/A |
|  | rs2003569 | Het (GA) vs Hom WT (GG) | South Korea (1 study) | NR | 0.96 (0.49, 1.87) | 64 | 155 | N/A |
|  |  | Hom MT (AA) vs Hom WT (GG) | South Korea (1 study) | NR | 1.20 (0.11, 13.53) | 49 | 117 | N/A |
|  | 686 C-A | Het (CA) vs Hom WT (CC) | Taiwan (1 study) | NR | 5.27 (0.69, 40.35) | 17 | 81 | N/A |
|  |  | Hom MT (AA) vs Hom WT (CC) | Taiwan (1 study) | NR | Data excluded^c^ | | | |
|  | TA_6_→TA_7_ at the promoter region | Het (TA_7_TA_6_) vs Hom WT (TA_6_TA_6_) | Taiwan (1 study) | NR | 1.12 (0.28, 4.46) | 17 | 81 | N/A |
|  |  | Hom MT (TA_7_TA_7_) vs Hom WT (TA_6_TA_6_) | Taiwan (1 study) | NR | Data excluded^c^ | | | |
|  | 1091 C-T | Het (CT) vs Hom WT (CC) | Taiwan (1 study) | NR | 5.00 (0.30, 84.17) | 17 | 81 | N/A |
|  |  | Hom MT (TT) vs Hom WT (CC) | Taiwan (1 study) | NR | Data excluded^c^ | | | |
| *UGT1A3* | rs2008584 | Het (GA) vs Hom WT (AA) | South Korea (1 study) | NR | 1.62 (0.89, 2.93) | 63 | 149 | N/A |
|  |  | Hom MT (GG) vs Hom WT (AA) | South Korea (1 study) | NR | 1.26 (0.31, 5.17) | 34 | 98 | N/A |
|  | rs6431625 | Het (TC) vs Hom WT (TT) | South Korea (1 study) | NR | 1.36 (0.64, 2.89) | 54 | 143 | N/A |
|  |  | Hom MT (CC) vs Hom WT (TT) | South Korea (1 study) | NR | 8.42 (0.34, 210.83) | 42 | 116 | N/A |
| *UGT2B7* | Rs7662029^g^ | Het (GA) vs Hom WT (AA) | Ethiopia (1 study) | NR | 0.69 (0.32, 1.50) | 36 | 124 | N/A |
|  |  | Hom MT (GG) vs Hom WT (AA) | Ethiopia (1 study) | NR | 0.38 (0.12, 1.15) | 19 | 74 | N/A |
| *XPO1* | rs7606167 | Het (GC) vs Hom WT (GG) | Japan (1 study) | NR | 0.90 (0.29, 2.80) | 15 | 73 | N/A |
|  |  | Hom MT (CC) vs Hom WT (GG) | Japan (1 study) | NR | 1.56 (0.35, 6.92) | 12 | 51 | N/A |
|  | rs11125883 | Het (AC) vs Hom WT (AA) | Japan (1 study) | NR | 0.33 (0.10, 1.07) | 16 | 64 | N/A |
|  |  | Hom MT (CC) vs Hom WT (AA) | Japan (1 study) | NR | 0.27 (0.05, 1.38) | 13 | 45 | N/A |
|  | rs1050567 | Het (GA) vs Hom WT (GG) | Japan (1 study) | NR | 0.59 (0.18, 1.91) | 14 | 76 | N/A |
|  |  | Hom MT (AA) vs Hom WT (GG) | Japan (1 study) | NR | 2.89 (0.67, 12.42) | 13 | 45 | N/A |

CI: confidence interval; Het: heterozygous; Hom: homozygous; MT: mutant-type; N/A: not applicable; NR: not reported; OR: odds ratio; WT: wild-type

^a^ One of the studies (Kim 2012b [GI: KIM]) reports WT to be T and MT to be C, but the other study (Yimer 2011), and the data, suggest that WT is C and MT is T

^b^ The study (Kim 2012b [GI: KIM]) reports WT to be del (-) and MT to be G, but the data suggests that WT is G and MT is del (-)

^c^ Data excluded due to zero counts

^d^ Reported in the same paper (Wang 2015b), but two separate cohorts of patients

^e^ One of the studies (Kim 2012b [GI: KIM]) reports WT to be G and MT to be T, but the other study (Chen 2015 [GI: ADACS]), and the data, suggest that WT is T and MT is G

^f^ Three of the studies (Chen 2015 [GI: ADACS], Li 2012 and Yimer 2011) report WT to be A and MT to be G, but the other study (Kim 2012b [GI: KIM]), and the data, suggest that WT is G and MT is A

^g^ The paper (Yimer 2011) reports WT to be G and MT to be A, but the data suggest that WT is A and MT is G
