## Additional File 9 for "Association of variants within the *GST* and other genes with anti-tubercular agents related toxicity: a systematic review and meta-analysis"

Other genetic variants and hepatotoxicity meta-analyses

### Pairwise comparisons for *ABCB1* rs1045642

*Heterozygous genotype (CT)* versus *homozygous wild-type (CC)*

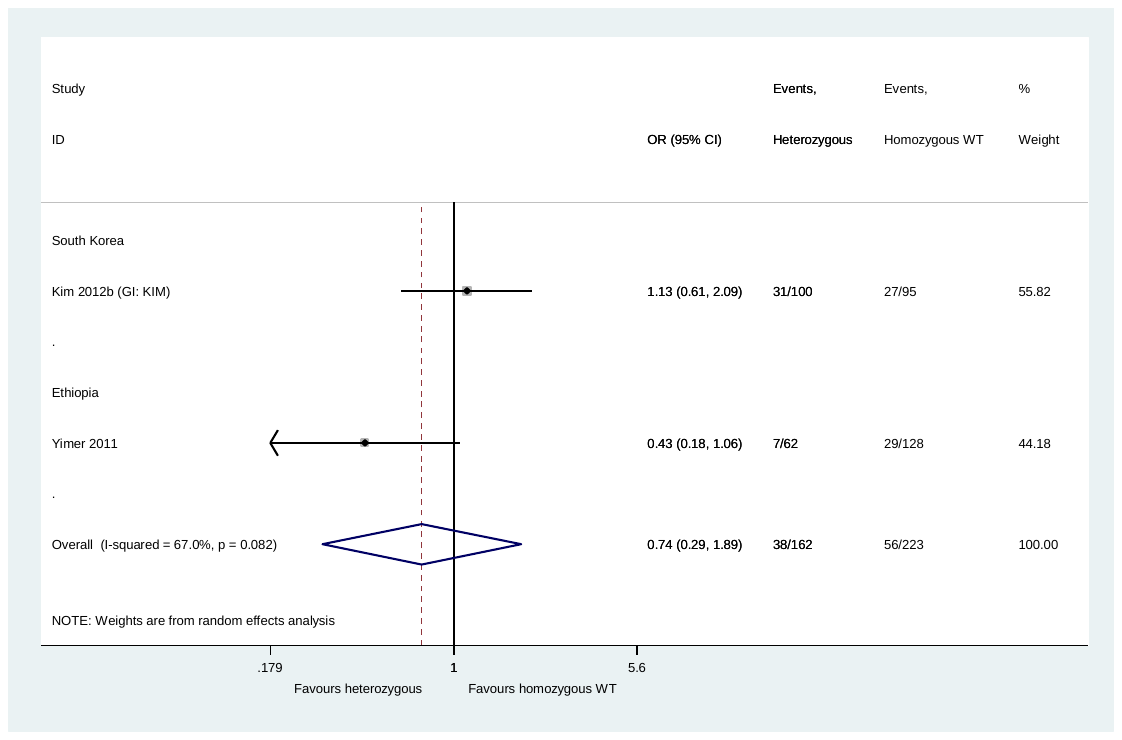

**Fig. 1** *ABCB1* rs1045642 and hepatotoxicity: heterozygous genotype (CT) versus homozygous wild-type (CC)

CI: confidence interval; GI: group identifier; OR: odds ratio; WT: wild-type

One of the studies (Kim 2012b [GI: KIM]) reports WT to be T and MT to be C, but the other study (Yimer 2011), and the data, suggest that WT is C and MT is T

*Homozygous mutant-type (TT)* versus *homozygous wild-type (CC)*

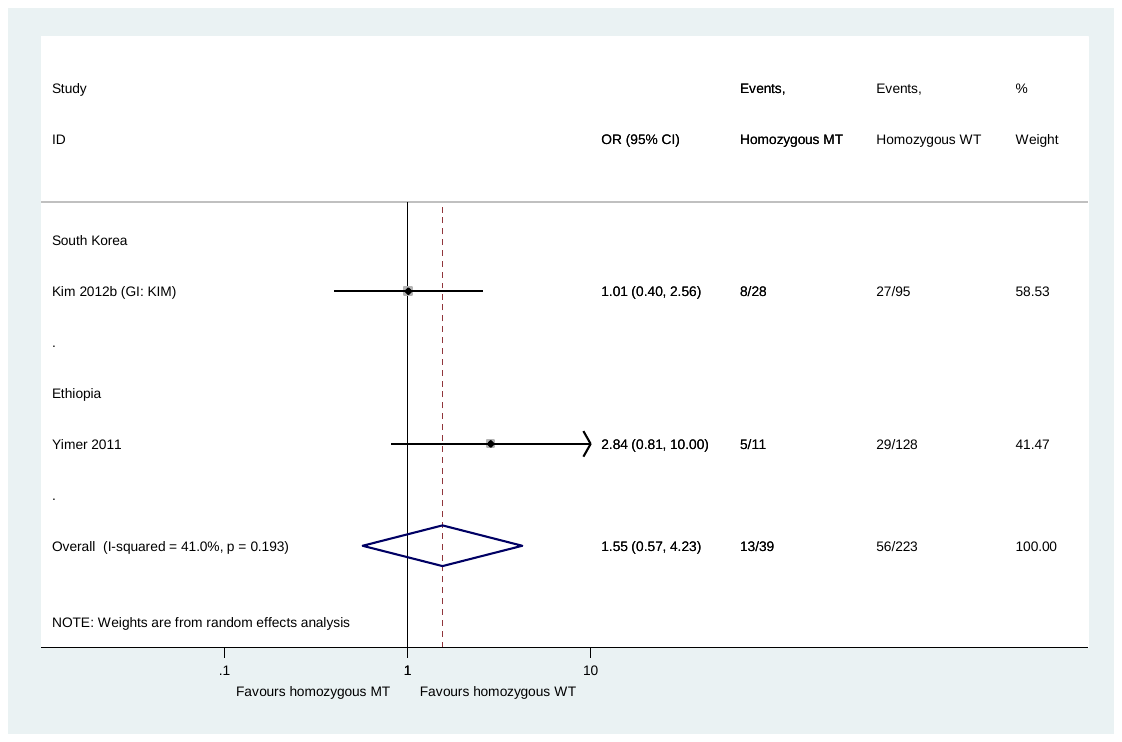

**Fig. 2** *ABCB1* rs1045642 and hepatotoxicity: homozygous mutant-type (TT) versus homozygous wild-type (CC)

CI: confidence interval; GI: group identifier; MT: mutant-type; OR: odds ratio; WT: wild-type

One of the studies (Kim 2012b [GI: KIM]) reports WT to be T and MT to be C, but the other study (Yimer 2011), and the data, suggest that WT is C and MT is T

### Pairwise comparisons for *NQO1* rs609C-T (rs1800566)

*Heterozygous genotype (CT)* versus *homozygous wild-type (CC)*

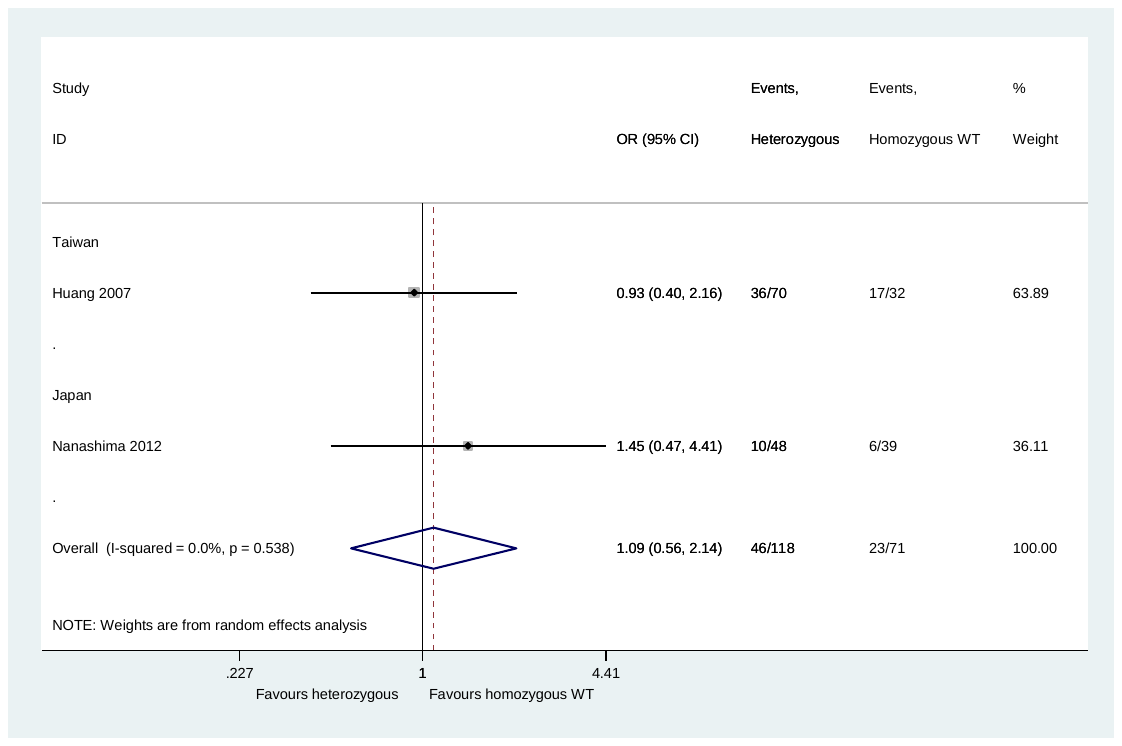

**Fig. 3** *NQO1* rs609C-T (rs1800566) and hepatotoxicity: heterozygous genotype (CT) versus homozygous wild-type (CC)

CI: confidence interval; OR: odds ratio; WT: wild-type

*Homozygous mutant-type (TT)* versus *homozygous wild-type (CC)*

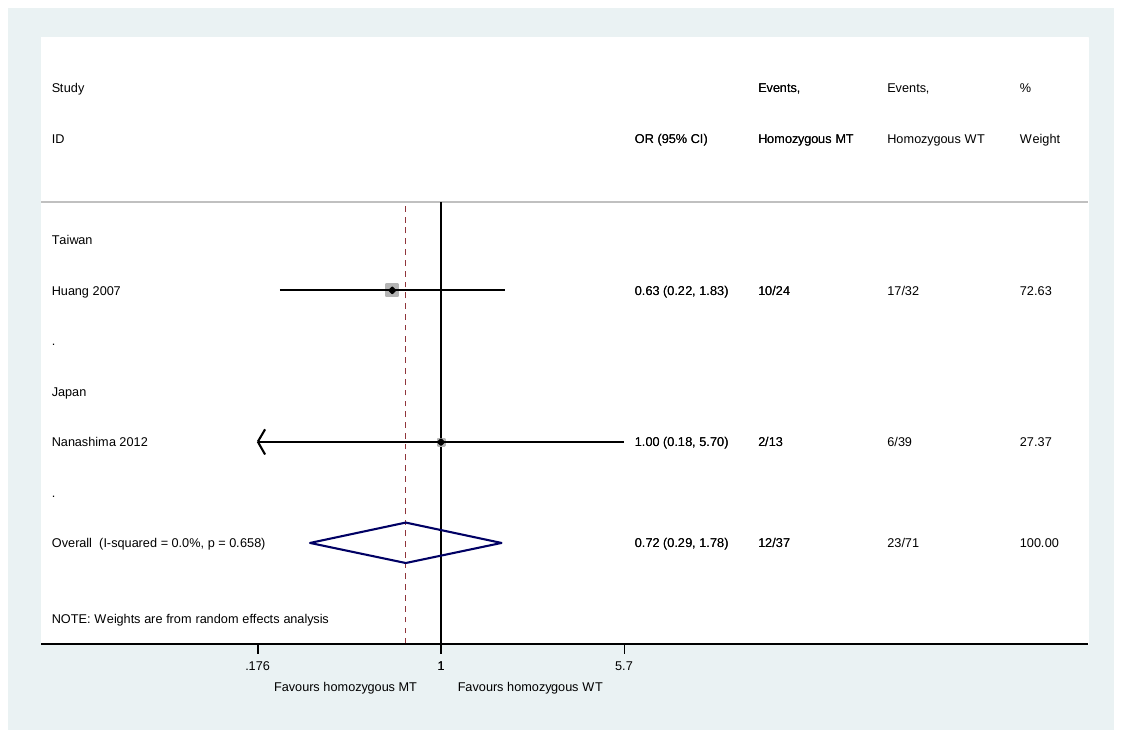

**Fig. 4** *NQO1* rs609C-T (rs1800566) and hepatotoxicity: homozygous mutant-type (TT) versus homozygous wild-type (CC)

CI: confidence interval; MT: mutant-type; OR: odds ratio; WT: wild-type

### Pairwise comparisons for *PXR* rs3814055

*Heterozygous genotype (CT)* versus *homozygous wild-type (CC)*

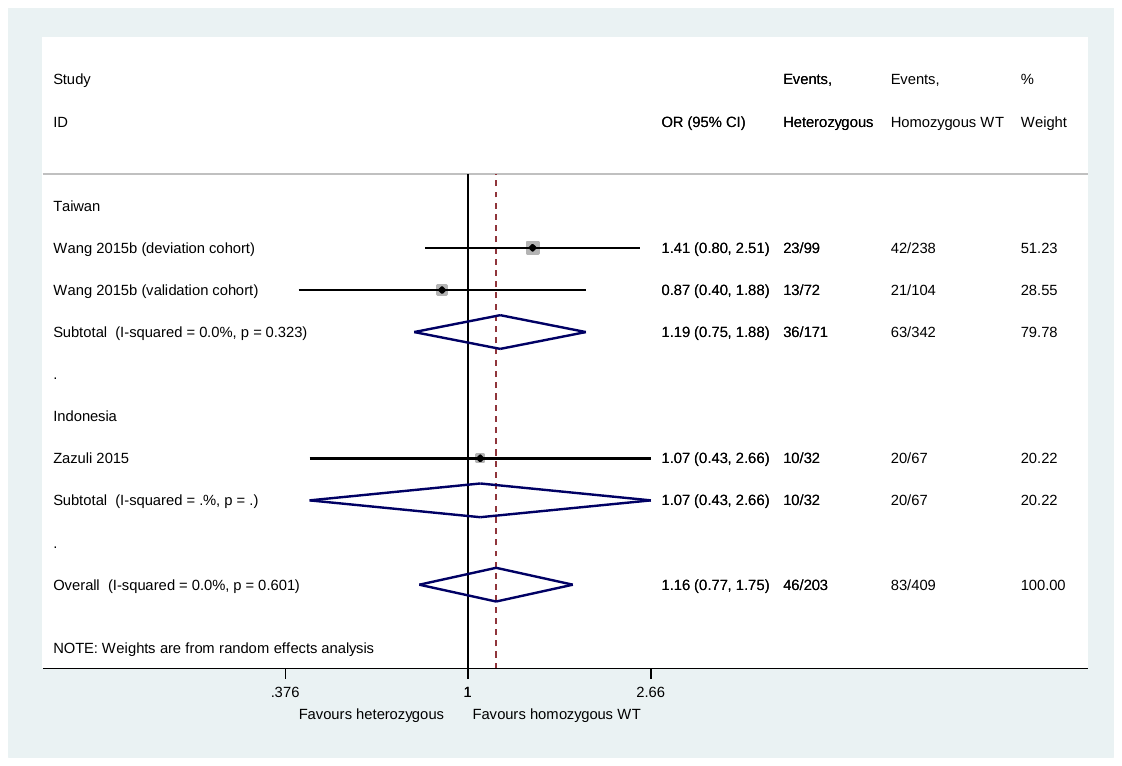

**Fig. 5** *PXR* rs3814055 and hepatotoxicity: heterozygous genotype (CT) versus homozygous wild-type (CC)

CI: confidence interval; OR: odds ratio; WT: wild-type

*Homozygous mutant-type (TT)* versus *homozygous wild-type (CC)*

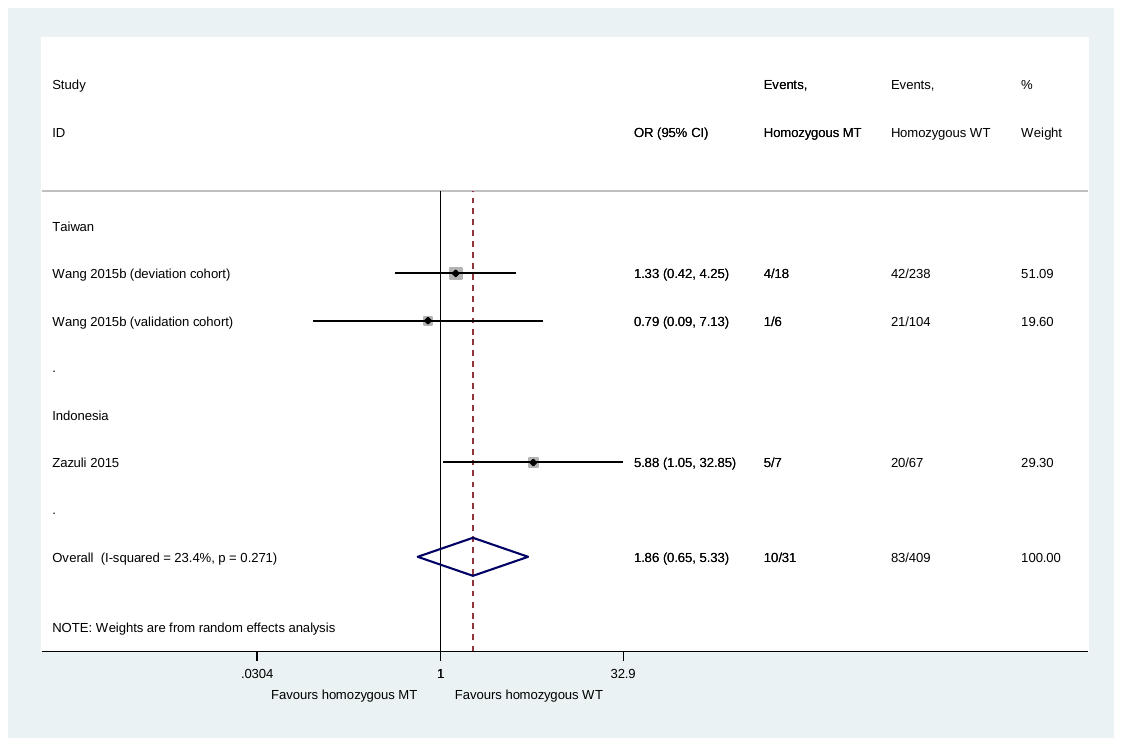

**Fig. 6** *PXR* rs3814055 and hepatotoxicity*:* homozygous mutant-type (TT) versus homozygous wild-type (CC)

CI: confidence interval; MT: mutant-type; OR: odds ratio; WT: wild-type

### Pairwise comparisons for *PXR* rs12488820

*Heterozygous genotype (CT)* versus *homozygous wild-type (CC)*

No meta-analysis performed as no patients had heterozygous genotype in either patient cohort.

*Homozygous mutant-type (TT)* versus *homozygous wild-type (CC)*

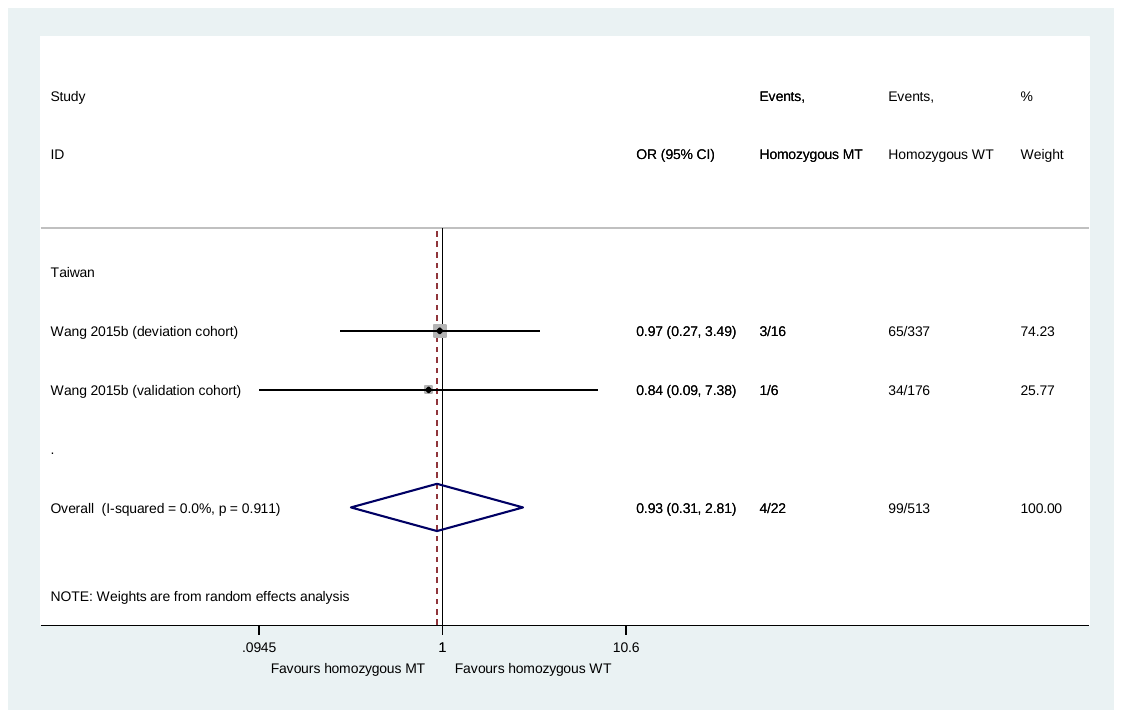

**Fig. 7** *PXR* rs12488820 and hepatotoxicity*:* homozygous mutant-type (TT) versus homozygous wild-type (CC)

CI: confidence interval; MT: mutant-type; OR: odds ratio; WT: wild-type

### Pairwise comparisons for *PXR* rs2461823

*Heterozygous genotype (GA)* versus *homozygous wild-type (GG)*

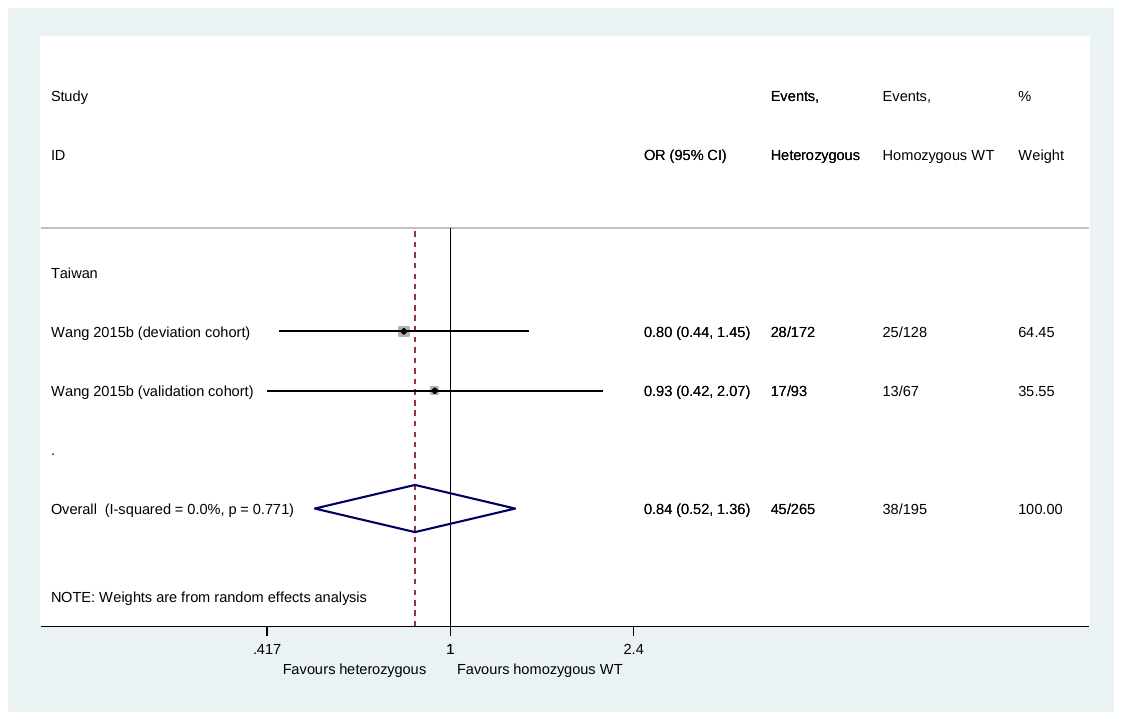

**Fig. 8** *PXR* rs2461823 and hepatotoxicity: heterozygous genotype (GA) versus homozygous wild-type (GG)

CI: confidence interval; OR: odds ratio; WT: wild-type

*Homozygous mutant-type (AA)* versus *homozygous wild-type (GG)*

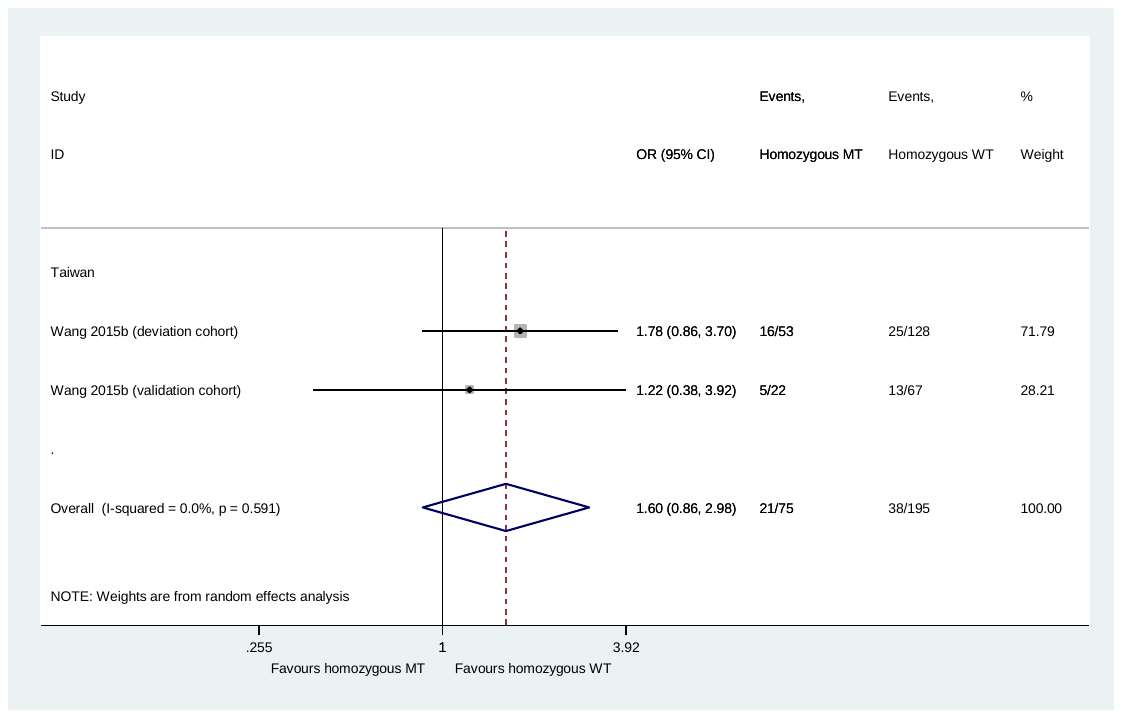

**Fig. 9** *PXR* rs2461823 and hepatotoxicity: homozygous mutant-type (AA) versus homozygous wild-type (GG)

CI: confidence interval; MT: mutant-type; OR: odds ratio; WT: wild-type

### Pairwise comparisons for *PXR* rs7643645

*Heterozygous genotype (AG)* versus *homozygous wild-type (AA)*

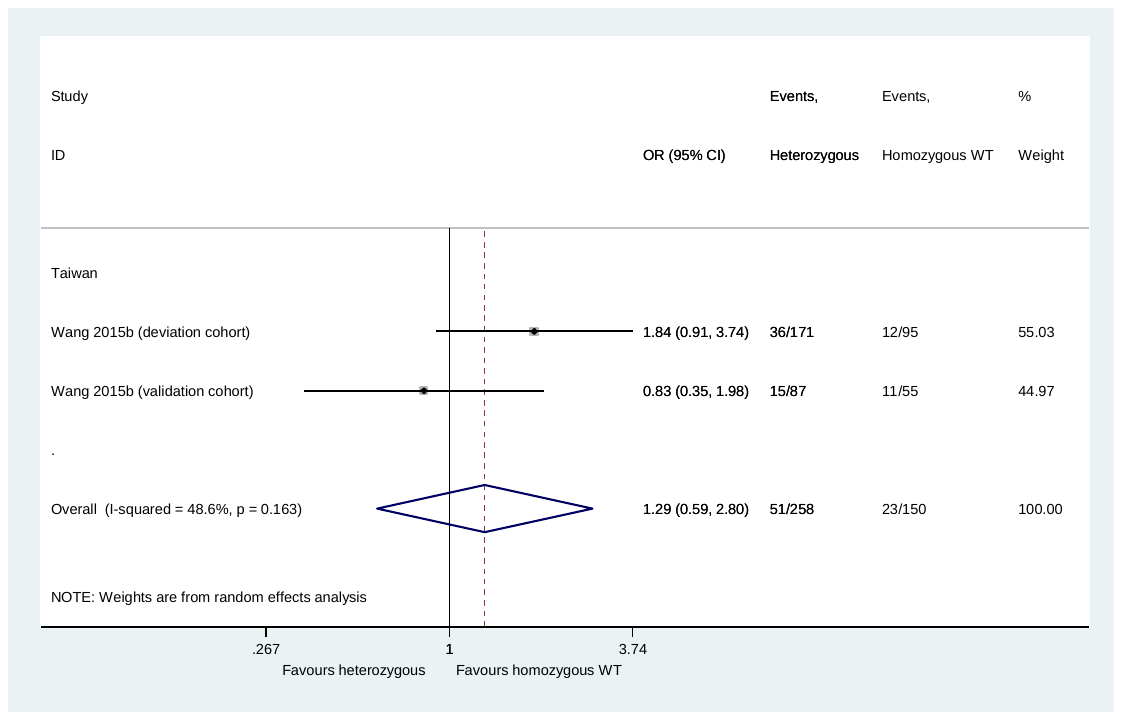

**Fig. 10** *PXR* rs7643645 and hepatotoxicity: heterozygous genotype (AG) versus homozygous wild-type (AA)

CI: confidence interval; OR: odds ratio; WT: wild-type

*Homozygous mutant-type (GG)* versus *homozygous wild-type (AA)*

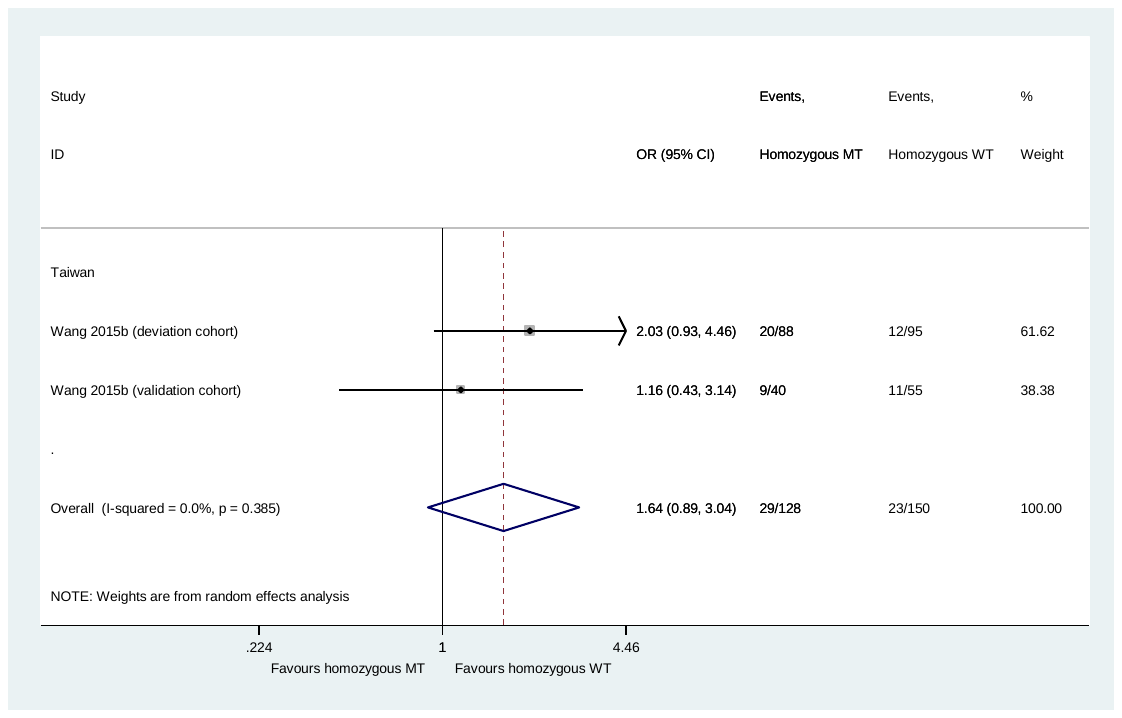

**Fig. 11** *PXR* rs7643645 and hepatotoxicity*:* homozygous mutant-type (GG) versus homozygous wild-type (AA)

CI: confidence interval; MT: mutant-type; OR: odds ratio; WT: wild-type

### Pairwise comparisons for *PXR* rs6785049

*Heterozygous genotype (GA)* versus *homozygous wild-type (GG)*

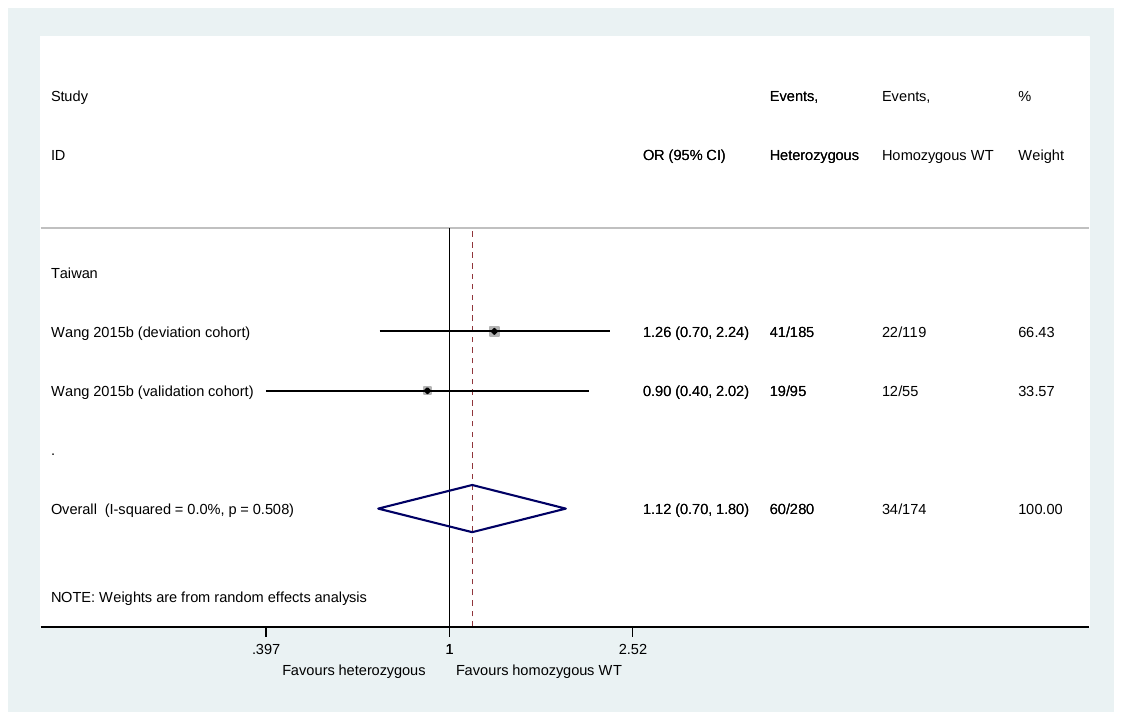

**Fig. 12** *PXR* rs6785049 and hepatotoxicity: heterozygous genotype (GA) versus homozygous wild-type (GG)

CI: confidence interval; OR: odds ratio; WT: wild-type

*Homozygous mutant-type (AA)* versus *homozygous wild-type (GG)*

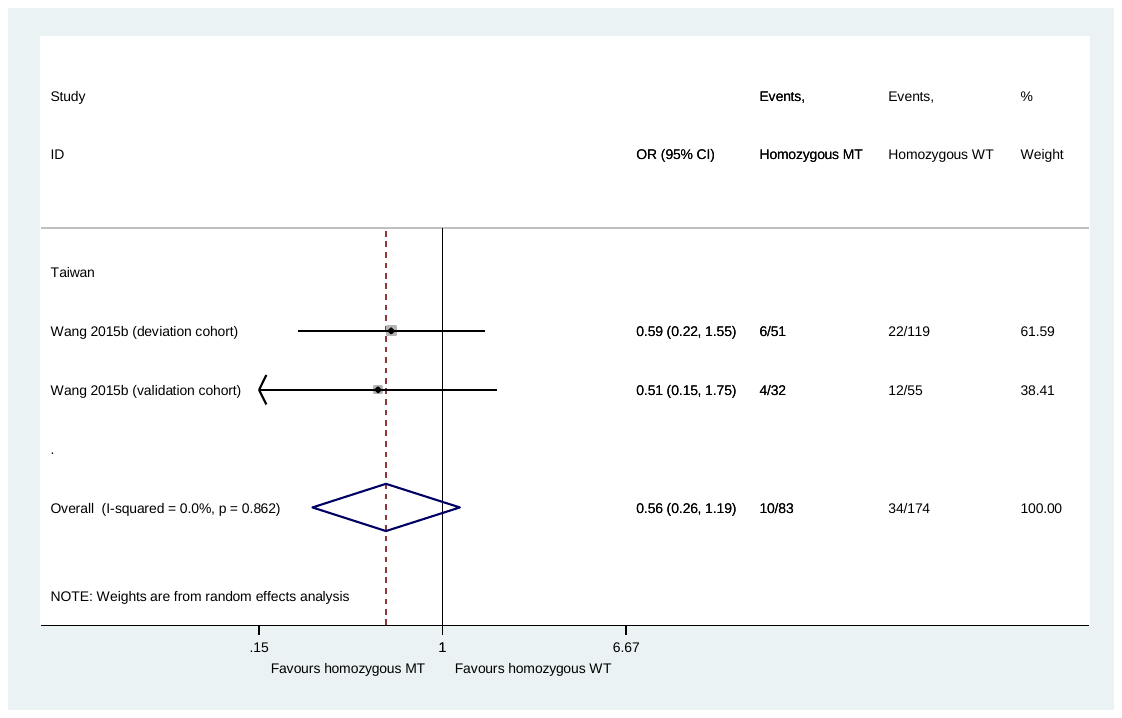

**Fig. 13** *PXR* rs6785049 and hepatotoxicity*:* homozygous mutant-type (AA) versus homozygous wild-type (GG)

CI: confidence interval; MT: mutant-type; OR: odds ratio; WT: wild-type

### Pairwise comparisons for *PXR* rs3814057

*Heterozygous genotype (AC)* versus *homozygous wild-type (AA)*

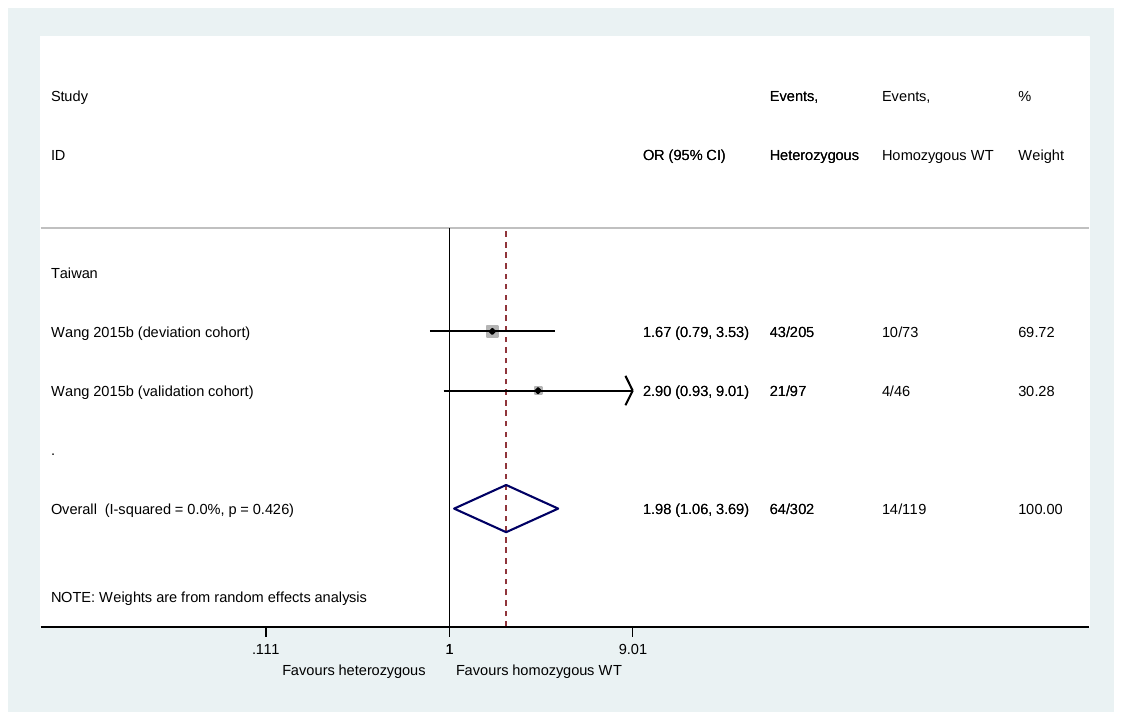

**Fig. 14** *PXR* rs3814057 and hepatotoxicity: heterozygous genotype (AC) versus homozygous wild-type (AA)

CI: confidence interval; OR: odds ratio; WT: wild-type

*Homozygous mutant-type (CC)* versus *homozygous wild-type (AA)*

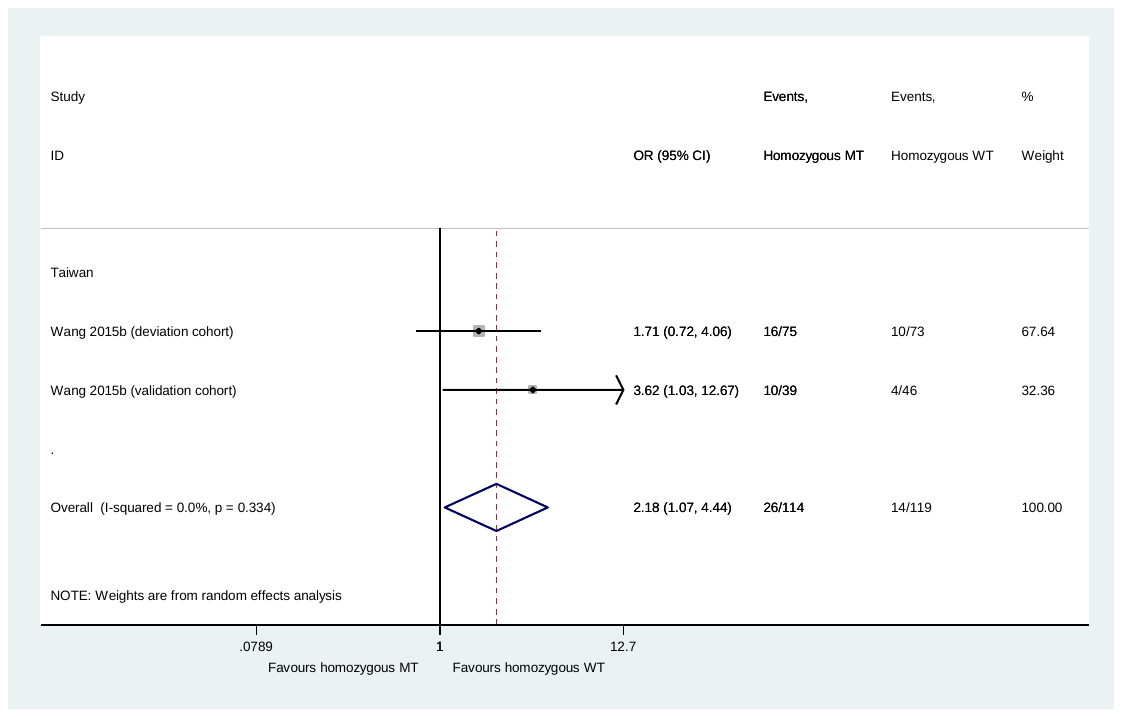

**Fig. 15** *PXR* rs3814057 and hepatotoxicity*:* homozygous mutant-type (CC) versus homozygous wild-type (AA)

CI: confidence interval; MT: mutant-type; OR: odds ratio; WT: wild-type

### Pairwise comparisons for *SLCO1B1* rs4149013

*Heterozygous genotype (AG)* versus *homozygous wild-type (AA)*

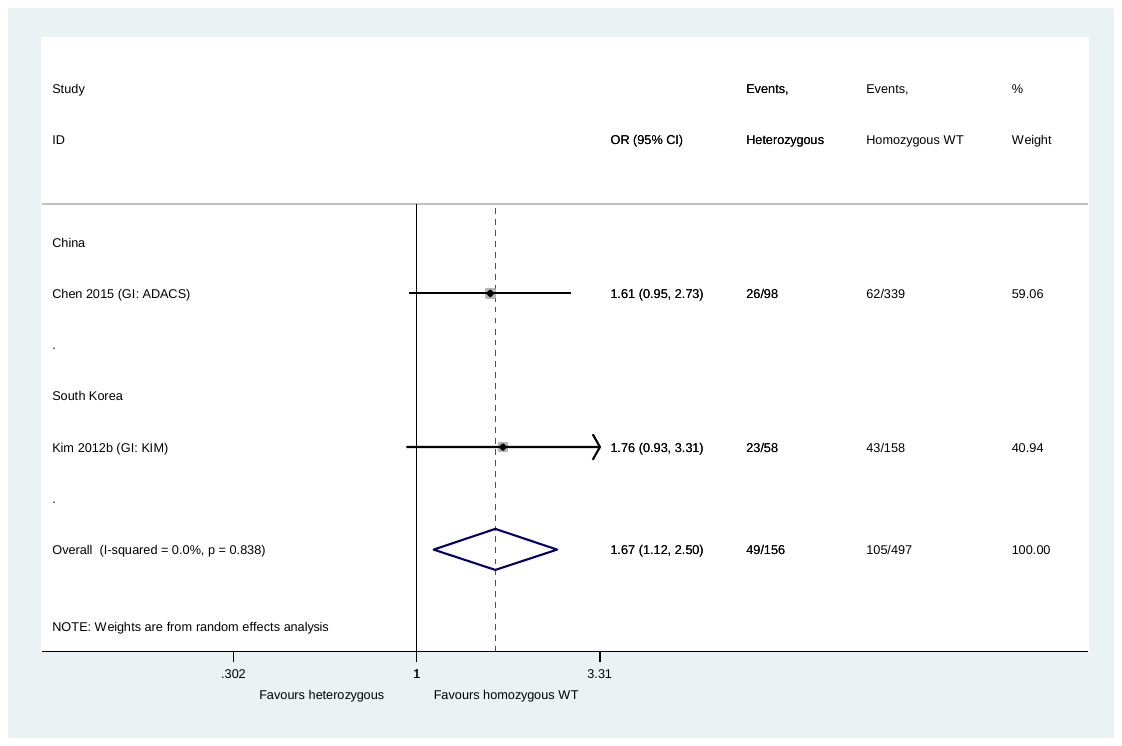

**Fig. 16** *SLCO1B1* rs4149013 and hepatotoxicity: heterozygous genotype (AG) versus homozygous wild-type (AA)

CI: confidence interval; GI: group identifier; OR: odds ratio; WT: wild-type

*Homozygous mutant-type (GG)* versus *homozygous wild-type (AA)*

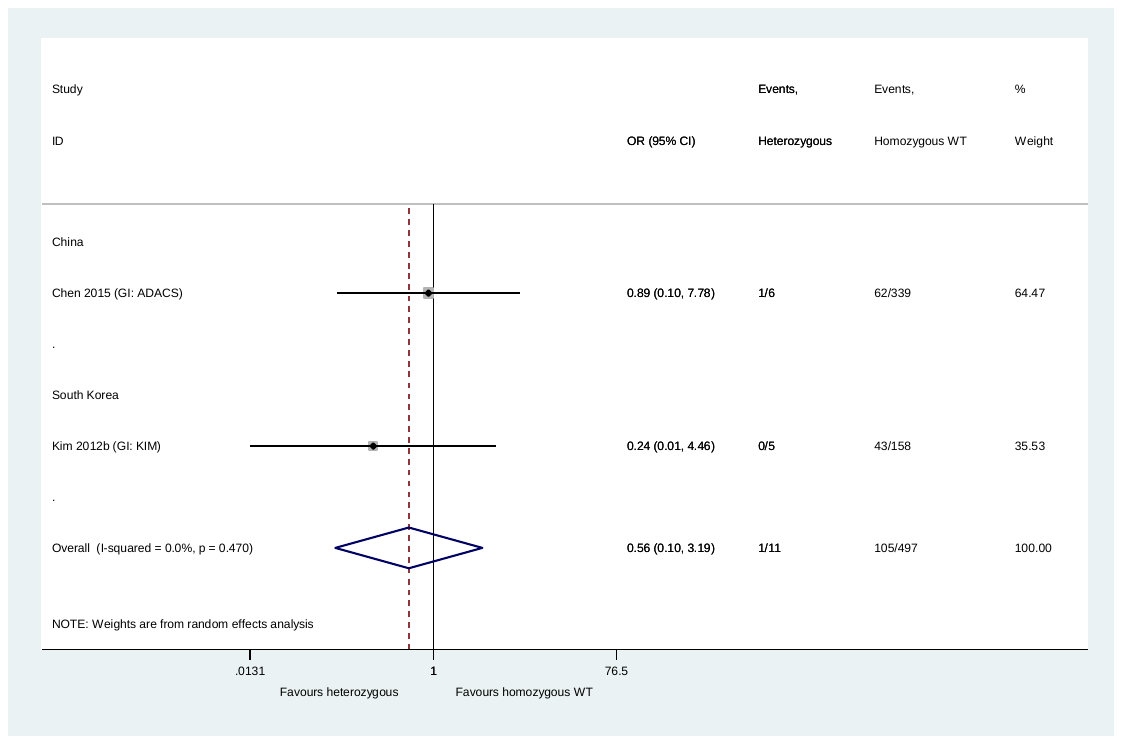

**Fig. 17** *SLCO1B1* rs4149013 and hepatotoxicity*:* homozygous mutant-type (GG) versus homozygous wild-type (AA)

CI: confidence interval; GI: group identifier; MT: mutant-type; OR: odds ratio; WT: wild-type

### **Pairwise comparisons for *SLCO1B1* rs4149014**

*Heterozygous genotype (GT)* versus *homozygous wild-type (TT)*

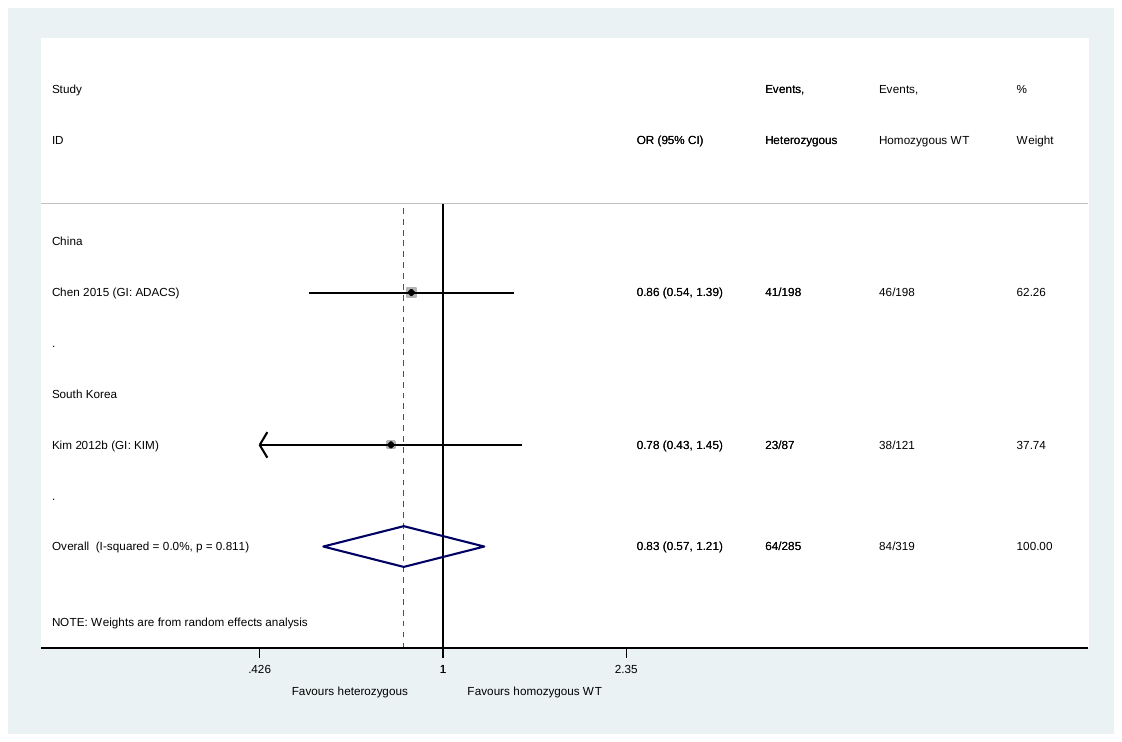

**Fig. 18** *SLCO1B1* rs4149014 and hepatotoxicity: heterozygous genotype (GT) versus homozygous wild-type (TT)

CI: confidence interval; GI: group identifier; OR: odds ratio; WT: wild-type

One of the studies (Kim 2012b [GI: KIM]) reports WT to be G and MT to be T, but the other study (Chen 2015 [GI: ADACS]), and the data, suggest that WT is T and MT is G

*Homozygous mutant-type (GG)* versus *homozygous wild-type (TT)*

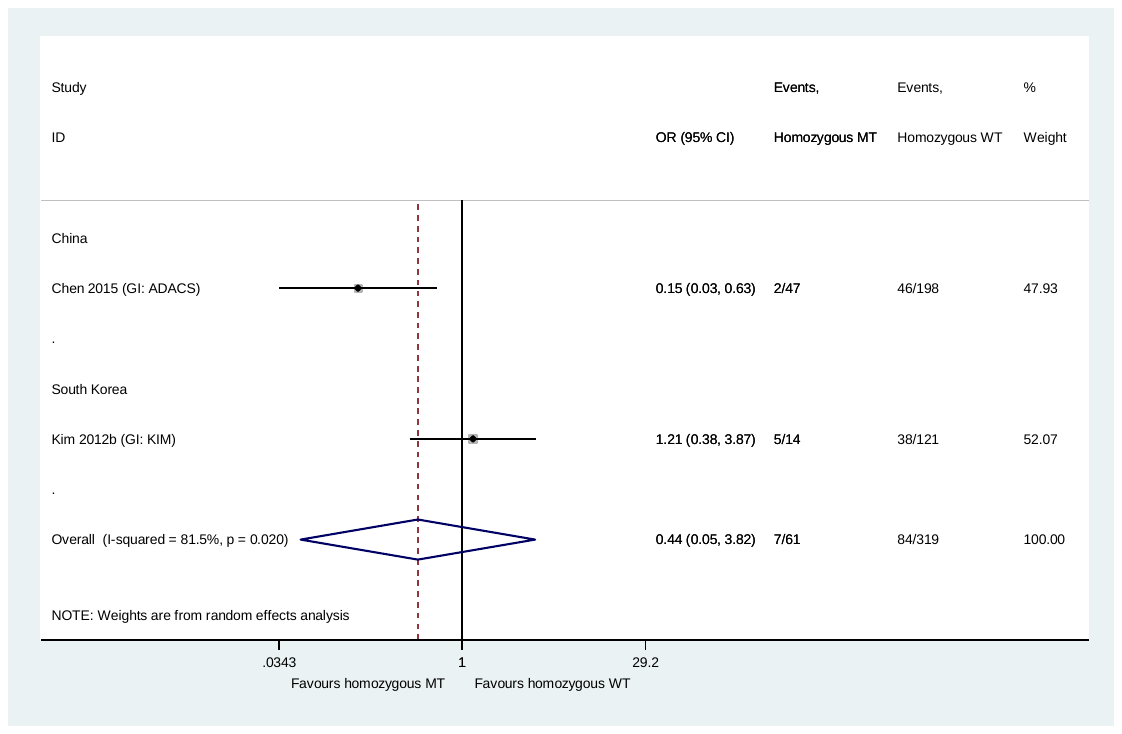

**Fig. 19** *SLCO1B1* rs4149014 and hepatotoxicity*:* homozygous mutant-type (GG) versus homozygous wild-type (TT)

CI: confidence interval; GI: group identifier; MT: mutant-type; OR: odds ratio; WT: wild-type

One of the studies (Kim 2012b [GI: KIM]) reports WT to be G and MT to be T, but the other study (Chen 2015 [GI: ADACS]), and the data, suggest that WT is T and MT is G

### Pairwise comparisons for *SLCO1B1* rs2306283

*Heterozygous genotype (GA)* versus *homozygous wild-type (GG)*

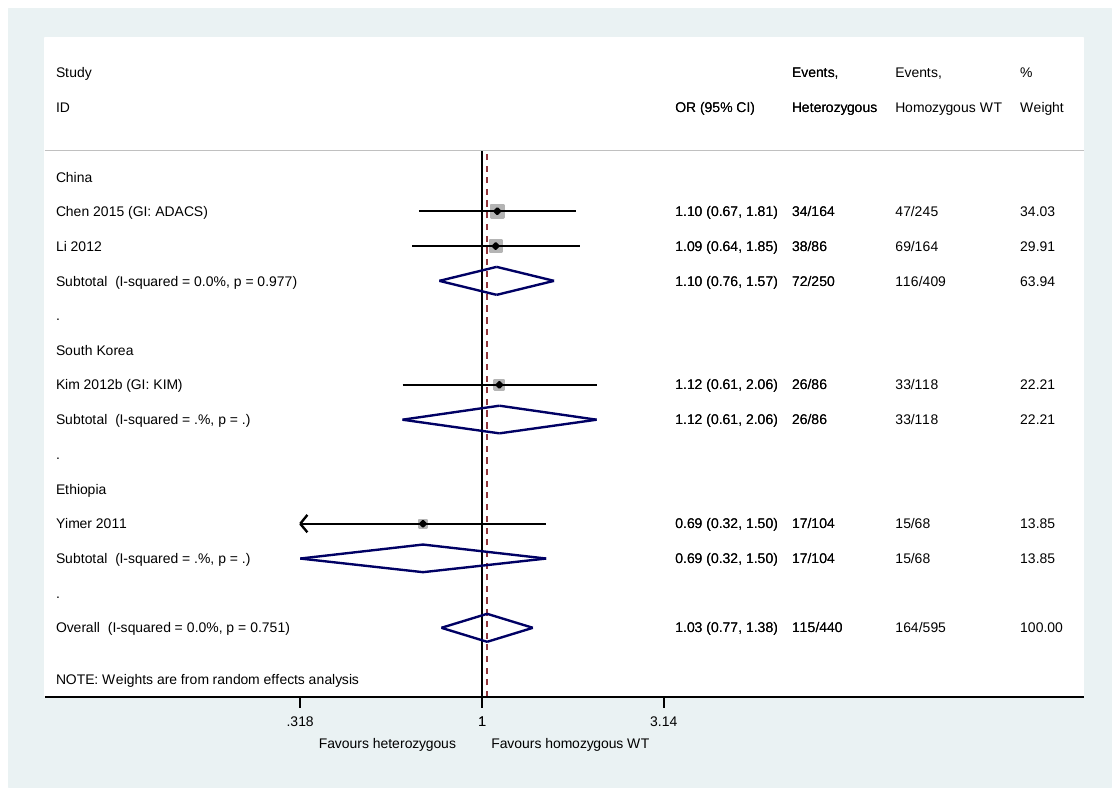

**Fig. 20** *SLCO1B1* rs2306283 and hepatotoxicity: heterozygous genotype (GA) versus homozygous wild-type (GG)

CI: confidence interval; GI: group identifier; OR: odds ratio; WT: wild-type

Three of the studies (Chen 2015 [GI: ADACS], Li 2012 and Yimer 2011) report WT to be A and MT to be G, but the other study (Kim 2012b [GI: KIM]), and the data, suggest that WT is G and MT is A

*Homozygous mutant-type (AA)* versus *homozygous wild-type (GG)*

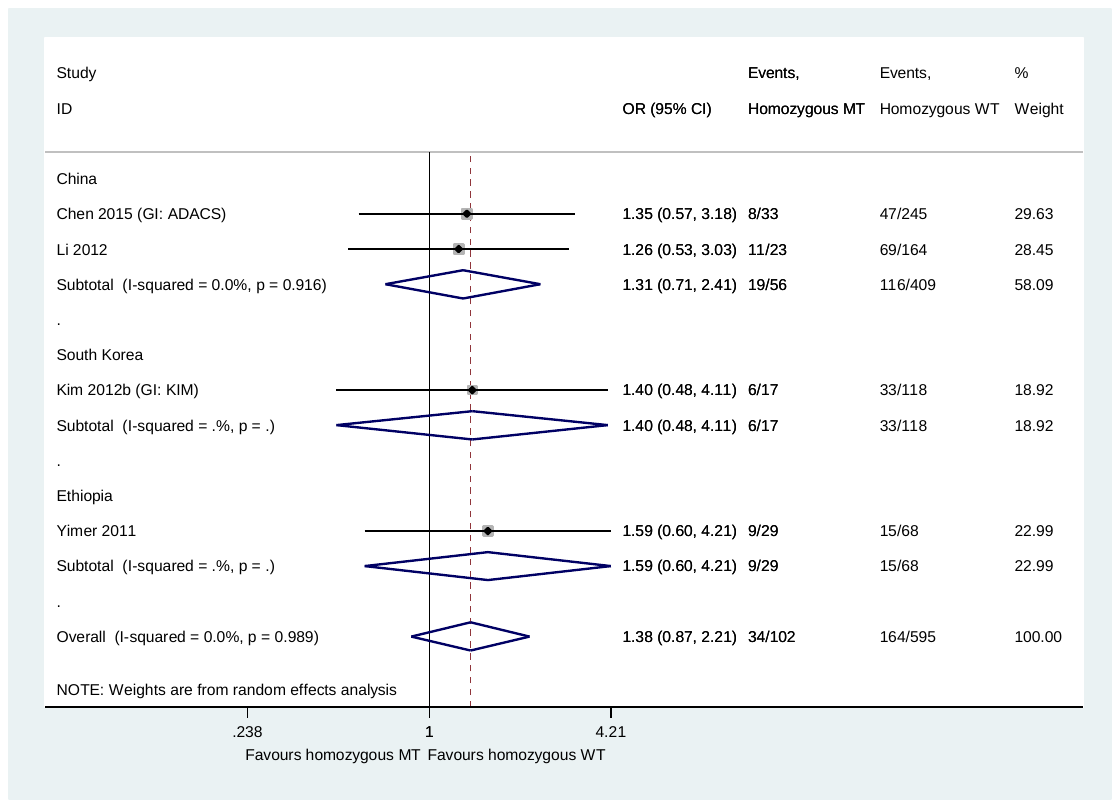

**Fig. 21** *SLCO1B1* rs2306283 and hepatotoxicity*:* homozygous mutant-type (AA) versus homozygous wild-type (GG)

CI: confidence interval; GI: group identifier; MT: mutant-type; OR: odds ratio; WT: wild-type

Three of the studies (Chen 2015 [GI: ADACS], Li 2012 and Yimer 2011) report WT to be A and MT to be G, but the other study (Kim 2012b [GI: KIM]), and the data, suggest that WT is G and MT is A

### Pairwise comparisons for *SLCO1B1* rs4149056

*Heterozygous genotype (TC)* versus *homozygous wild-type (TT)*

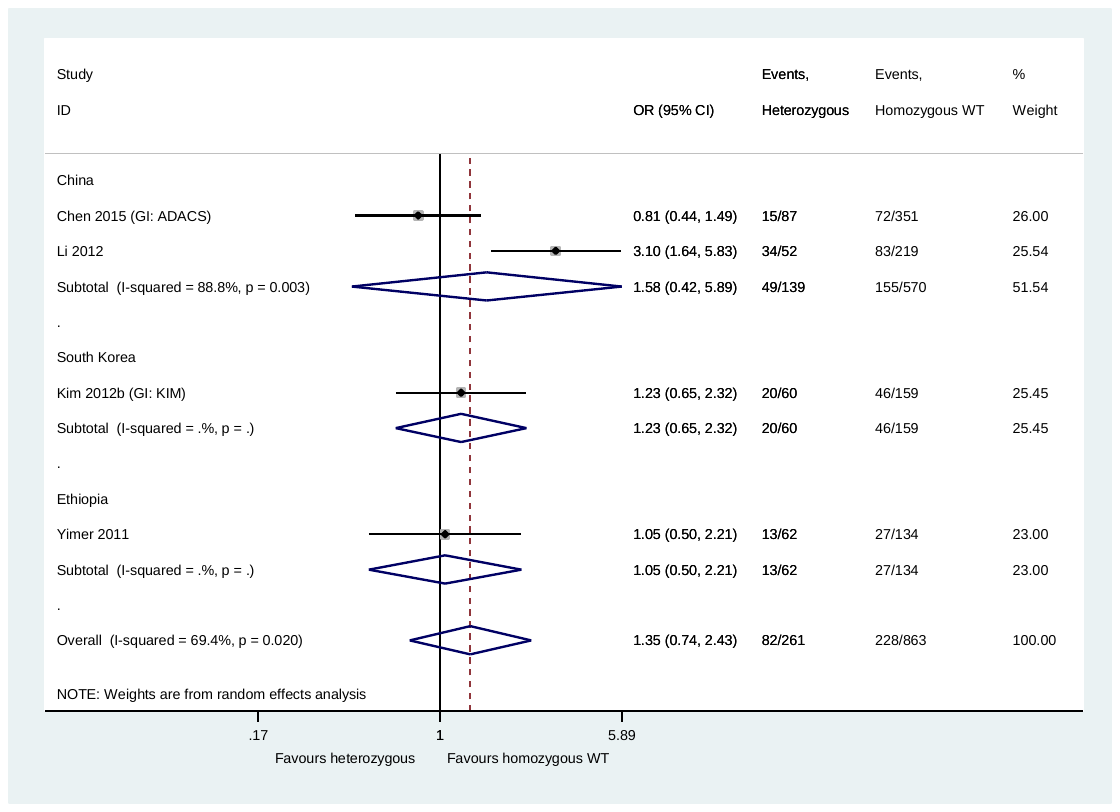

**Fig. 22** *SLCO1B1* rs4149056 and hepatotoxicity: heterozygous genotype (TC) versus homozygous wild-type (TT)

CI: confidence interval; GI: group identifier; OR: odds ratio; WT: wild-type

*Homozygous mutant-type (CC)* versus *homozygous wild-type (TT)*

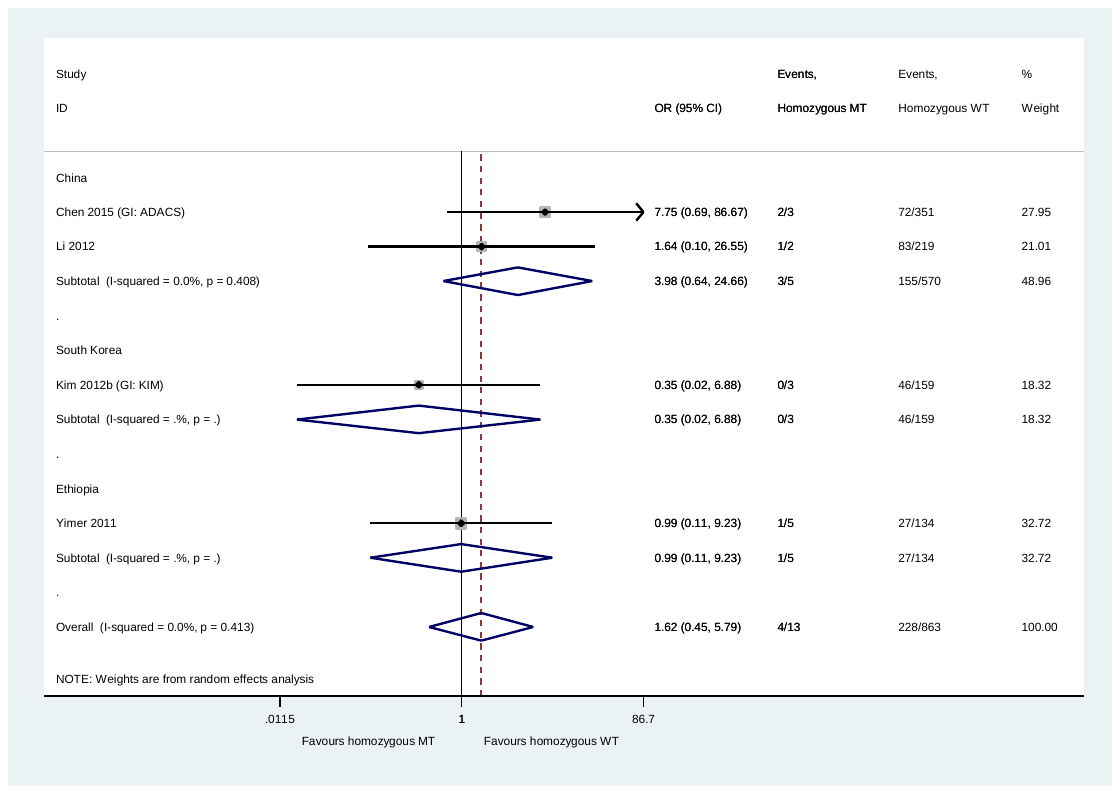

**Fig 23** *SLCO1B1* rs4149056 and hepatotoxicity*:* homozygous mutant-type (CC) versus homozygous wild-type (TT)

CI: confidence interval; GI: group identifier; MT: mutant-type; OR: odds ratio; WT: wild-type

### *SOD2* rs4880: *Homozygous mutant-type (CC) or heterozygous genotype (CT)* versus *homozygous wild-type (TT)*

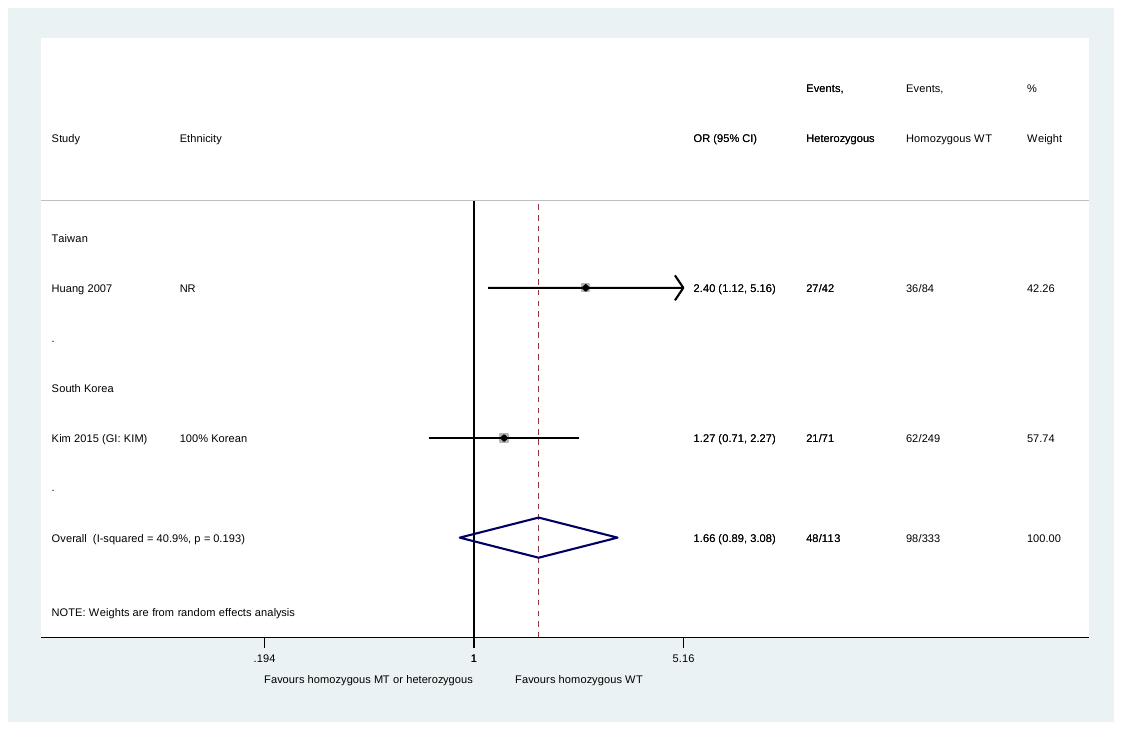

**Fig. 24** *SOD2* rs4880 and hepatotoxicity: homozygous mutant-type (CC) or heterozygous genotype (CT) versus homozygous wild-type (TT)

CI: confidence interval; GI: group identifier; MT: mutant-type; OR: odds ratio; WT: wild-type

### Pairwise comparisons for *UGT1A1* rs4148323

*Heterozygous genotype (GA)* versus *homozygous wild-type (GG)*

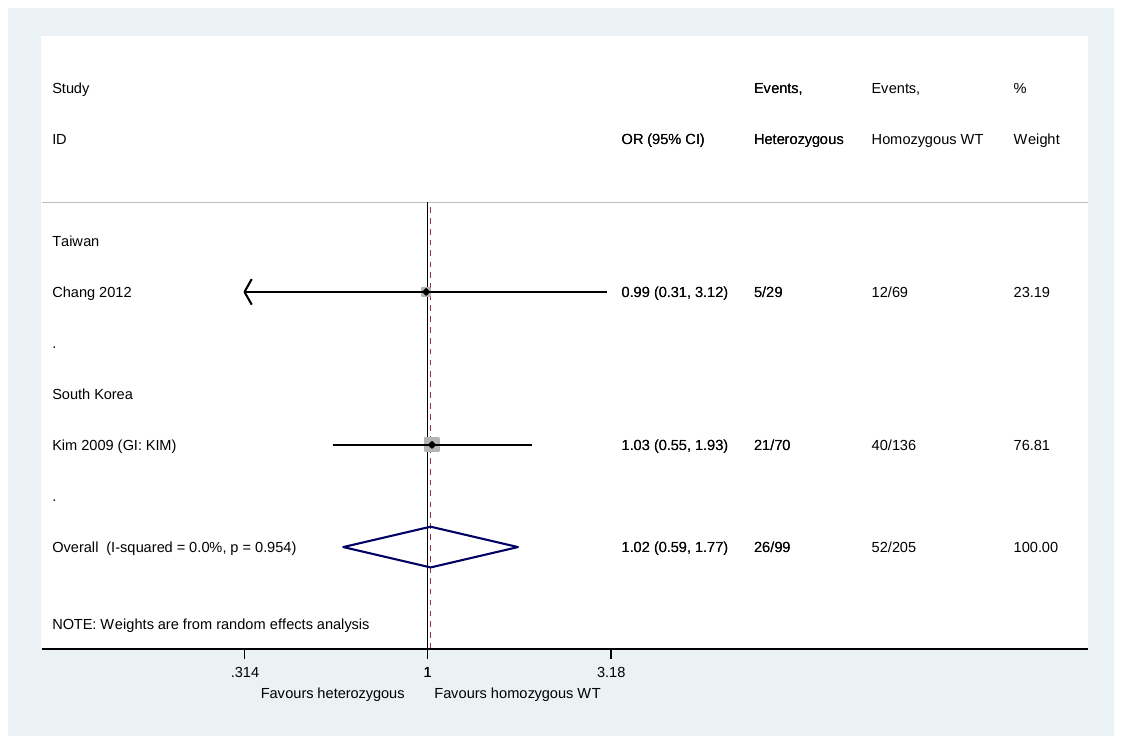

**Fig. 25** *UGT1A1* rs4148323 and hepatotoxicity*:* heterozygous (GA) versus homozygous wild-type (GG)

CI: confidence interval; GI: group identifier; OR: odds ratio; WT: wild-type

*Homozygous mutant-type (TT)* versus *homozygous wild-type (CC)*

No meta-analysis performed as only one study identified patients with homozygous mutant-type genotype.
