## Additional File 10 for "Association of variants within the *GST* and other genes with anti-tubercular agents related toxicity: a systematic review and meta-analysis"

**Additional file 10** *GST* and other genetic variants and toxicity outcomes (other than hepatotoxicity)

| Study | Country | Ethnicity | Outcome | Gene | Variant | Comparison | OR (95% CI) | # cases | # controls |
| --- | --- | --- | --- | --- | --- | --- | --- | --- | --- |
| Costa (2012) | Brazil | 84% Black/mixed race, 16% other | ADRs | *GSTM1* | Null or present status | Null (Hom null) vs not null (Het or Hom present) | 1.06 (0.42, 2.72) | 43 | 45 |
|  |  |  |  | *GSTT1* | Null or present status | Null (Hom null) vs not null (Het or Hom present) | 0.71 (0.29, 1.73) | 43 | 45 |
| Kim 2012a (GI: KIM) | South Korea | NR | ATD-induced MPE | *ABCB1/*  *MDR1* | I1145I (rs1045642) | Het (CT) vs Hom WT (CC) | 1.02 (0.55, 1.89) | 57 | 137 |
|  |  |  |  |  |  | Hom MT (TT) vs Hom WT (CC) | 0.49 (0.15, 1.55) | 32 | 88 |
|  |  |  |  |  | -114918 T-G (rs10261685) | Het (TG) vs Hom WT (TT) | 1.10 (0.52, 2.32) | 61 | 159 |
|  |  |  |  |  |  | Hom MT (GG) vs Hom WT (TT) | 7.91 (0.32, 197.42) | 50 | 130 |
|  |  |  |  | *ABCC2* | 1774 G-del^a^ | Het (G/-) vs Hom WT (GG) | 0.62 (0.32, 1.18) | 54 | 137 |
|  |  |  |  |  |  | Hom MT (-/-) vs Hom WT (GG) | 0.75 (0.29, 1.96) | 31 | 63 |
|  |  |  |  |  | -1549 G-A (rs1885301) | Het (GA) vs Hom WT (GG) | 1.19 (0.63, 2.23) | 54 | 149 |
|  |  |  |  |  |  | Hom MT (AA) vs Hom WT (GG) | 3.39 (1.13, 10.14) | 38 | 96 |
|  |  |  |  |  | -24 C-T (rs717620) | Het (CT) vs Hom WT (CC) | 1.12 (0.60, 2.10) | 56 | 149 |
|  |  |  |  |  |  | Hom MT (TT) vs Hom WT (CC) | 2.79 (0.84, 9.25) | 39 | 98 |
|  |  |  |  |  | IVS3-49 C-T (rs2804400) | Het (CT) vs Hom WT (CC) | 1.19 (0.64, 2.23) | 54 | 152 |
|  |  |  |  |  |  | Hom MT (TT) vs Hom WT (CC) | 3.47 (1.16, 10.36) | 38 | 98 |
|  |  |  |  |  | V417L (rs2273697) | Het (GA) vs Hom WT (GG) | 1.01 (0.44, 2.32) | 61 | 157 |
|  |  |  |  |  |  | Hom MT (AA) vs Hom WT (GG) | 1.29 (0.11, 14.52) | 53 | 136 |
|  |  |  |  |  | S978S (rs3740070) | Het (GA) vs Hom WT (GG) | 0.84 (0.26, 2.73) | 62 | 159 |
|  |  |  |  |  |  | Hom MT (AA) vs Hom WT (GG) | Data excluded^b^ | | |
|  |  |  |  |  | I1324I (rs3740066) | Het (CT) vs Hom WT (CC) | 1.40 (0.75, 2.63) | 54 | 147 |
|  |  |  |  |  |  | Hom MT (TT) vs Hom WT (CC) | 2.79 (0.99, 7.89) | 37 | 100 |
| Kim 2010 (GI: KIM) | South Korea | NR | ATD-induced cutaneous reactions | *GSTM1* | Null or present status | Null (Hom null) vs not null (Het or Hom present) | 1.22 (0.74, 2.01) | 94 | 190 |
|  |  |  |  | *GSTT1* | Null or present status | Null (Hom null) vs not null (Het or Hom present) | 1.19 (0.72, 1.96) | 94 | 190 |

ADR: adverse drug reaction; ATD: anti-tuberculosis drug; CI: confidence interval; GI: group identifier; Het: heterozygous; Hom: homozygous; MPE: maculopapular eruption; MT: mutant-type; NR: not reported; OR: odds ratio; WT: wild-type

^a^ The study (Kim 2012a [GI: KIM]) reports WT to be del (-) and MT to be G, but the data suggests that WT is G and MT is del (-)

^b^ Data excluded due to zero counts
